# Lateral interactions between antiparallel filaments drive self-assembly of the FtsZ ring in bacterial cell division

**DOI:** 10.64898/2026.11.24.769015

**Authors:** M.B.L. McAndrew, Y. Zhang, L. Rohland, J.O.H. Warren, G. Carloni, M. Krstic, L.P. Gregory, D. Kureisaite-Ciziene, A. Sogues, D.M. Roberts, S. Holden, A. Šarić, A. Wehenkel, J. Löwe

**Affiliations:** MRC Laboratory of Molecular Biology, Cambridge CB2 0QH, UK; Institut Pasteur, Université Paris Cité, CNRS UMR 3528, Bacterial Cell Cycle Mechanisms Unit, F-75015 Paris, France; Institute of Science and Technology (ISTA), Austria, Klosterneuburg, 3400, Austria; School of Life Sciences, University of Warwick, Gibbet Hill Campus, Coventry CV4 7AL, UK

**Author notes:** Corresponding authors: Jan Löwe. These authors contributed equally (M.B.L. McAndrew, Y. Zhang, L. Rohland, J.O.H. Warren, G. Carloni, M. Krstic).

## Abstract

The Z-ring is the organiser of cell division in most archaea and bacteria, arriving early at mid-cell and recruiting downstream proteins. While FtsZ protofilaments and their treadmilling dynamics are well understood, how FtsZ assembles into condensed Z-rings remains unknown. Here we determined the *Escherichia coli* in-cell electron cryotomography (cryo-ET) structure of FtsZ to 6.9 Å resolution, showing antiparallel double filaments with the C-terminus facing the membrane. Electron cryomicroscopy (cryo-EM) of purified *E. coli* FtsZ showed these double filaments self-assemble. Cryo-ET of cells showed ordering of FtsZ doublets into bands, which we revealed is regulated by the disordered N-terminal residues. Structures of purified *Bacillus subtilis* and *Thermotoga maritima* FtsZ demonstrated that antiparallel lateral interactions are a conserved feature. We describe how self-assembly of the Z-ring occurs through two types of lateral interactions and propose a new role for FtsZ treadmilling in reorganising antiparallel lateral interactions during cell constriction.

**Highlights:**

- *Escherichia coli* FtsZ forms the same antiparallel double filaments in cells and on its own, in vitro
- Neighbouring *E. coli double* filaments are organised into bands of filaments and are regulated by the disordered N-terminus of FtsZ
- In vitro structures of *Thermotoga maritima* and *Bacillus subtilis* FtsZ reveal partially conserved lateral interactions
- Perturbations to lateral *interactions* disrupt cell division and viability
- FtsZ treadmilling is modulated by lateral interactions and is proposed to enable Z-ring reduction in diameter during cell constriction

## Introduction

Cell division is fundamental to a bacterium’s lifecycle. For most walled bacteria, it requires the complex balancing of septal cell wall synthesis and remodelling, membrane constriction and chromosome segregation. For rod-shaped cells, such as *Escherichia coli* and *Bacillus subtilis*, Gram negative and positive model organisms, these events happen at mid-cell and are organised by the Z-ring, the centrepiece of a multi-protein cell division machine called the divisome^1–4^. First discovered in 1991^5^, the Z-ring is formed of filaments of the tubulin homologue FtsZ^6^, which condense early in cell division, before septal constriction has commenced^7^. The ring then tracks the decreasing diameter of the septal constriction, until just before septal closure, upon which it decondenses to make new Z-rings at the sites of the next cell division events^8^. Septal constriction is driven by the processive movement of peptidoglycan (PG) synthases (core divisome complexes) that lay down layers of crosslinked glycan strands on the outside of the cell membrane, pushing it inwards mechanically. Generally, FtsZ has no direct membrane-binding properties and instead achieves membrane tethering through binding to its membrane anchors^9–12^. The two most conserved of these are the actin homologue FtsA^13,14^, and SepF in gram-positives and Actinomycetes (and archaea)^15–18^, both of which bind membranes through amphipathic helices. *E. coli* and some other bacteria also have ZipA^10^. In all cases, tethering is achieved through binding to the very C-terminal peptide (CTP) of FtsZ^9,12^, which sits at the end of an often long and disordered C-terminal tail (CTT), ranging from ∼50 residues in *E. coli* to over 150 residues in *Caulobacter crescentus*. Because of the length of the CTT, the main body of FtsZ filaments is located nanometres away from the membrane in cells^19,20^.

FtsZ polymerises into tubulin-like protofilaments, single chains of molecules arranged head-to-tail, with the GTPase sites sandwiched between subunits and formed by residues from two subunits so that polymerisation switches on GTP hydrolysis. FtsZ protofilaments exhibit GTP-hydrolysis-driven dynamics known as treadmilling^21^. For this, FtsZ subunits bind GTP to polymerise onto the plus end of a growing and stable protofilament and diffuse away from the minus end following GTP hydrolysis. This is enabled by a conformation change termed the cytomotive switch, which is a feature conserved across all known actin and tubulin homologues^22,23^. FtsZ protofilament treadmilling achieves movement through subunit exchange, without requiring the protofilament or its constituent subunits to move. The purpose of FtsZ treadmilling has been the source of speculation. When it was first observed *in vivo*, a model whereby FtsZ treadmilling drives cell wall synthesis through linking treadmilling to PG synthase movement was proposed^24,25^. However, in Gram positive bacteria studied so far, there appears to be no correlation between processive synthase speeds and FtsZ treadmilling^26–28^. Additionally, in *E. coli*, perturbations to FtsZ treadmilling (FtsZ G105S), had no meaningful effect on septal closure rates, suggesting that there is likely no direct link between septal closure and FtsZ treadmilling^29^. This leaves the precise role of FtsZ treadmilling mechanistically undefined and its roles have been proposed as contributing towards Z-ring dynamics, to early and uniform synthase distribution^30^ and/or to the potential for protofilament alignment and condensation^31,32^.

How protofilaments are arranged to make a condensed Z-ring has remained largely elusive^33^. Several studies have proposed FtsZ lateral interfaces^34–38^, but without an accepted consensus. In *Bacillus subtilis,* depletions of FtsZ-binding proteins were shown to impact Z-ring condensation but showed no effect when removing single Z-binding proteins^39^. Similarly, the FtsZ-associated proteins (Zaps) ZapA, ZapC and ZapD are all implicated in FtsZ crosslinking^40–42^, yet deletion of all three Zaps is required to produce poor viability and cell filamentation phenotypes^42^. A recent single particle electron cryomicroscopy (cryo-EM) structure of FtsZ protofilaments bound to ZapA dimers provides structural insights into how FtsZ can be bridged by ZapA^43^. Similarly, a cryo-EM and electron cryotomography (cryo-ET) study revealed FtsZ bundles bridged by ZapD^44^. Their dispensability however suggests some functional redundancy and perhaps points towards intrinsic FtsZ lateral self-assembly properties that are reinforced by Zaps.

Several cryo-ET studies have sought to visualise the Z-ring *in situ*, consistently detecting filamentous densities positioned several nanometres away from the cytoplasmic membrane. Early tomograms of dividing *C. crescentus* first segmented putative FtsZ filaments at the division site^45^, while subsequent higher-resolution imaging in C*. crescentus* and *E. coli* resolved these assemblies into bands of filaments^19^. More recently, cryogenic Focussed Ion Beam (cryo-FIB) milling was used to thin *B. subtilis* cells to produce higher contrast tilt series and tomograms^20^. These showed ordered arrangements of filaments lining invaginating septa. Despite these *in situ* views, none of these studies achieved the resolution required to determine how protofilaments associate within bands, establish their relative orientations, or confirm their molecular identity as FtsZ. Here we present the intrinsic self-assembly properties of FtsZ protofilaments that enable them to form a Z-ring. We employed in-cell cryo-ET and subtomogram averaging, single particle cryo-EM of purified proteins, coarse-grain simulations of filament dynamics and live cell imaging to tackle the question of how FtsZ protofilaments assemble laterally into a ring. Our results reveal conserved features of Z-ring self-assembly properties and propose a model for how treadmilling modifies FtsZ lateral interactions to allow the Z-ring to track the cell’s decreasing diameter during constriction.

## Results

### FtsZ self-assembles into antiparallel double filaments in *Escherichia coli*

The bacterial Z-ring is a large assembly of FtsZ protofilaments localised to mid-cell. To gain insights into the Z-ring ultrastructure in Gram negative *E. coli* cells, we prepared log phase cultures of *E. coli* C41(DE3) cells for cryogenic focussed ion beam (cryo-FIB) milling under cryogenic conditions, to produce cellular cross-sections of thicknesses required to make them electron transparent. We consistently obtained sub-200 nm thick lamellae, which proved essential for identifying FtsZ filaments in the cellular volumes? obtained subsequently by electron cryotomography (cryo-ET). Filaments were challenging to identify due to their size and curvature but were most clear when their long axis was parallel to the electron beam. To aid in visualising FtsZ, tomogram membranes and ribosomes were segmented using Easymode^46^, while FtsZ filaments were manually segmented. These tomograms revealed FtsZ protofilaments typically arrange into doublets of filaments with some single filaments also observed (Figure 1A-B). We could occasionally observe additional membrane-bound double filament densities between the FtsZ filaments and the membrane. While it is likely these are the actin homologue and cell division protein FtsA that anchors FtsZ to the membrane, we could not conclusively assign its identity.

**Figure 1.**
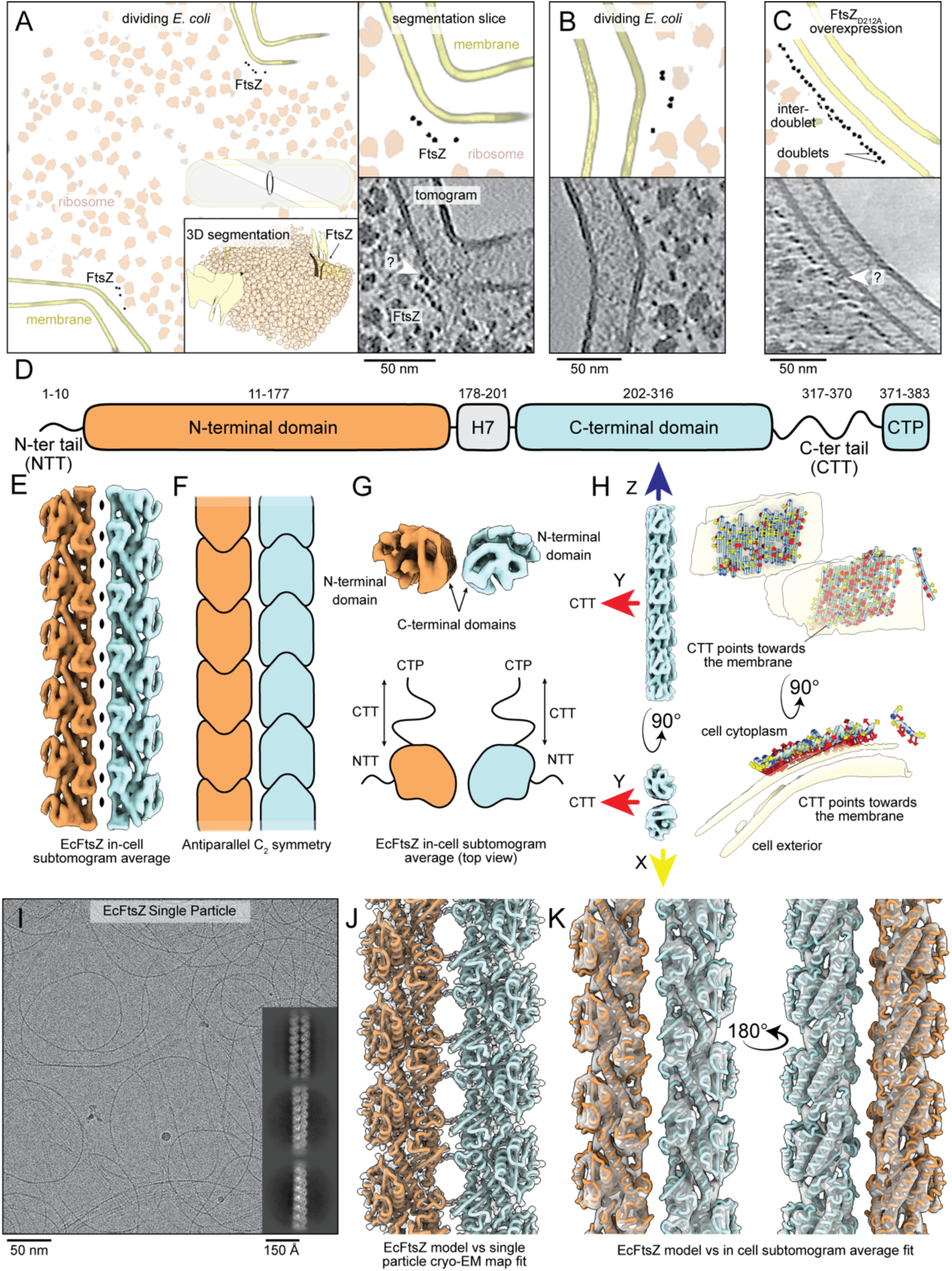
*E. coli* FtsZ double filaments *in situ* and *in vitro*. (A) Tomogram of a cryo-FIB milled dividing *E. coli* cell (“WT”: C41(DE3)). The large panel shows a central slice of the tomogram 3D segmentation (membranes, ribosomes and FtsZ filaments; FtsZ in black). Top right is a magnified 2D slice of one side of the invaginating septum, below is the unsegmented tomogram. An unassigned double filament density is highlighted on the membrane (“?”). Insets show position of the lamella imaged in the cellular context and a 3D rendering of the segmentation, including FtsZ in black (B) A second example of FtsZ filaments in WT dividing *E. coli*. (C) A tomogram slice of overexpressed *E. coli* FtsZ_D212A_ used to subtomogram average FtsZ with an unassigned membrane-anchored density between FtsZ and the membrane (“?”). (D) *E. coli* FtsZ (EcFtsZ) domain architecture. (E) Subtomogram average map of EcFtsZ double filaments at 6.9 Å resolution, with black ovals showing C_2_ symmetry axes. (F) Schematic of antiparallel FtsZ filaments. (G) Top view of the double filament subtomogram average showing the relative positions of the N-terminal and C-terminal domains. Below is a schematic showing the direction of the flexible, unresolved NTT and CTT regions. (H) Refined subtomogram average coordinates mapped back into a cellular tomogram. Arrows represent direction of X, Y and Z axes with the CTT pointing in the direction of Y (red arrows). Positions of refined coordinates shown with respect to the tomogram membrane segmentation. (I) Cryo-EM micrograph of purified EcFtsZ with example 2D classes of double filaments. (J) 2.6 Å single particle cryo-EM map and atomic FtsZ double filament structure obtained from it. (K) EcFtsZ double filament structure docked into the in-cell subtomogram average shown in E.

We attempted to get higher resolution insights into these filaments using subtomogram averaging (STA) where every observed protofilament was annotated in Relion 5^47^. When refined against a cylinder (to eliminate initial model bias), a clear double filament with subunits was resolved, albeit limited to ∼20-30 Å resolution due to the low particle count (Figure S1). To overcome this, we overexpressed a GTP hydrolysis deficient FtsZ_D212A_ in *E. coli* C41(DE3) cells as previously described^19^, thinned cross sections using cryo-FIB milling and performed cryo-ET. The resulting tomograms exposed rows of membrane-aligned protofilament doublets (Figure 1C), from which STA resolved an antiparallel double filament with C_2_ symmetry (Figure 1EF, S2). The symmetry axes runs along the short axis of the doublet (“D1” in Relion convention), meaning the unresolved disordered C-terminal tails (CTT) protrude out from the protofilaments in the same direction (Figure 1G). Given the C-terminal peptide (CTP) at the very end of the CTT has been shown to interact with FtsZ’s membrane anchor FtsA, we tested if overexpressed EcFtsZ double filaments direct the CTT towards the membrane for FtsA binding(citation). For this, refined poses of each subtomogram were placed back into their respective tomograms using ArtiaX^48^, showing the vast majority of membrane adjacent doublets aligned their CTT towards the membrane despite not having overexpressed FtsA and FtsA generally being present in sub stoichiometric amounts in E. coli (Figure 1H).

Although our STA map of EcFtsZ resolved secondary structure features allowing rigid docking of EcFtsZ monomers, its resolution of 6.9 Å was not sufficient to resolve individual residues at the interface between protofilaments within the doublet. Therefore, we purified, plunge froze and collected single particle analysis (SPA) cryo-EM datasets of EcFtsZ polymerised with the non-hydrolysable GTP analogue GMPCPP. To mitigate air-water interface (AWI) adsorption during the vitrification process, we employed the sandwich Langmuir-Blodgett method that avoided FtsZ filaments adhering strongly to the AWIs (manuscript submitted^49^). The micrographs and 2D class averages revealed EcFtsZ had polymerised into double filaments (Figure 1I, S3) and subsequent processing produced a high resolution 2.7 Å reconstruction with the same C_2_ symmetry as the STA map, allowing for confident atomic model building of EcFtsZ protofilaments (Figure 1J). The resulting model fit extremely well into the in-cell STA map, with a model-map Fourier Shell Correlation (FSC) resolution 3of 8.7 Å at the 0.5 threshold (Figure S2I), giving us confidence that interpretations of the structure obtained from purified EcFtsZ accurately reflected FtsZ in a cellular environment. Importantly, this also demonstrated that doublet formation *in situ* is an FtsZ-intrinsic property and not dependent on factors present in the cytoplasm.

### R174 is essential to the doublet interface in *E. coli* FtsZ

With the atomic structure of FtsZ double filaments in hand, we examined the interface between protofilaments. We had seen from the in-cell STA map that the globular C-terminal domains of both protofilaments are positioned inwards towards the centre of the doublet (Figure 1DGK). But we had no information on the residues involved in the interaction. The atomic structure revealed that the protofilament interface is small and consists mostly of a single positively charged arginine (R174) on one protofilament, on the loop between helix 6 (H6) and H7, which protrudes into a pocket of negative charges (D234, E237, E238, E241) on the C-terminal H9 of the other protofilament (Figure 2A). Having noted this residue’s importance had been discovered previously in a genetic screen in 2003 by Koppleman *et al.*^50^, we further probed the physiological importance of the interface, initially using a previously developed FtsZ temperature sensitive (ts) strain where at 42°C (non-permissive temperature) replication of the plasmid carrying the sole copy of FtsZ is inhibited, allowing us to supplement with an additional plasmid of WT or mutant EcFtsZ (Figure S4)^51^. Our spot assays (Figure 2B, S5) showed complete loss of viability when the charge of R174 was either reversed (R174D) or removed (R174A), in agreement with the previous work of Koppelman *et al*^50^. In fact, both mutations to R174 showed a strong dominant negative phenotype, in the presence of WT FtsZ (not shifted to non-permissive temperature). Conversely, when we mutated the other side of the interface, no single point mutations of any of the negatively charged residues resulted in an observable growth phenotype. Only when removing all four negative charges could we reproduce a strong growth defect, yet still without the dominant negative phenotype of R174 mutations. We also confirmed that all non-viable mutants shown in Figure 2B have the typical cell division phenotype of strongly elongated cells by light microscopy (Figure S6).

**Figure 2.**
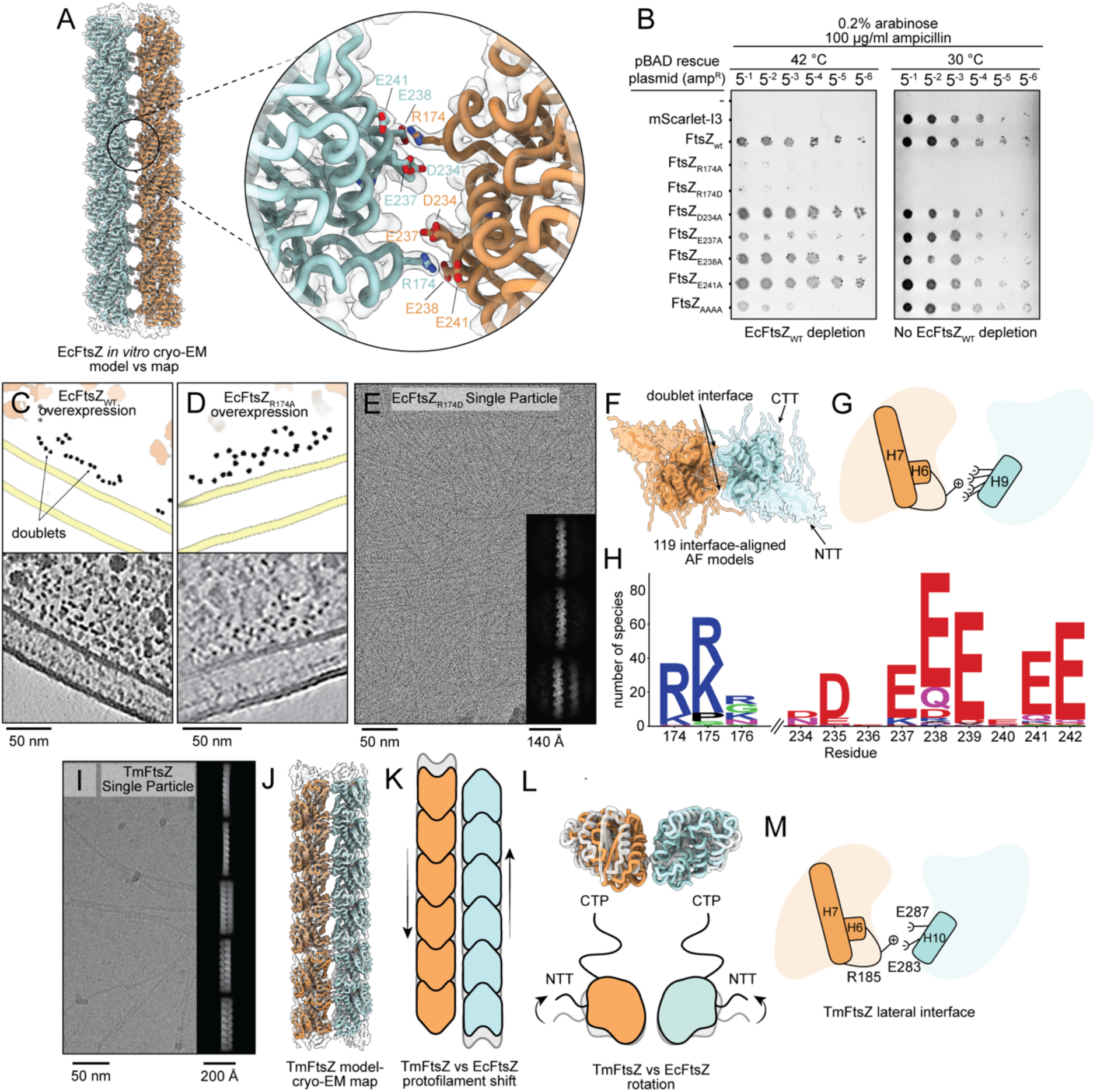
The double filament interface and its conservation. (A) Residues mediating contacts between protofilaments in the EcFtsZ doublet shown alongside the transparent cryo-EM map. (B) Spot dilution assay of temperature sensitive strains for WT FtsZ depletion, supplemented with a pBAD plasmid harbouring FtsZ mutants (and mScarlet-I3 control). (C) Tomogram 2D slice of overexpressed WT FtsZ in *E. coli* C41 (DE3). Top is the 2D slice of the segmentation and bottom is the denoised tomogram. (D) Tomogram 2D slice of overexpressed FtsZ_R174A_ in *E. coli*. (E) A representative micrograph crowded with purified FtsZ_R174D_ with example 2D classes of single filaments, despite the high concentration. (F) 119 FtsZ AF2 monomer predictions from Pseudomonadota, aligned to the doublet interface. (G) Schematic of the EcFtsZ doublet interface per subunit showing motifs probed in panels F and H. (H) Weblogo of AF2 predictions of interfacial residues between FtsZ from Pseudomonadota. (I) A representative EM micrograph and 2D classes showing TmFtsZ double filaments used in the data processing to determine the TmFtsZ single particle cryo-EM structure. (J) Double filament cryo-EM structure of *Thermotoga maritima* FtsZ (TmFtsZ). (K) Schematic of the longitudinal shift of TmFtsZ compared to EcFtsZ. (L) A top view of TmFtsZ vs EcFtsZ double filaments and accompanying schematic showing a rotational difference between the double filaments. (M) Schematic of the TmFtsZ per-subunit double filament interface. The positive charge is provided by an arginine on the H6-H7 loop and likely interacts with negative charges on H10, adjacent to the H9 motif involved in the EcFtsZ doublet interface.

From the spot assays and light microscopy, it was clear R174 was important for cell viability and cell division, but we wanted to understand what happened to mutant protofilaments *in situ* using cryo-ET. To do this we overexpressed EcFtsZ_R174A_ in *E. coli* C41(DE3) cells, to compare with a WT EcFtsZ overexpression, milled lamellae using cryo-FIB and reconstructed tomograms. The annotated FtsZ protofilaments in Figure 2CD show a stark difference in their arrangements. WT FtsZ overexpression appeared to arrange in rows of doublets proximal to the membrane as expected, whereas R174A overexpression showed a distinct absence of doublets with single protofilaments in a disordered bundle. Importantly, this also held true when we purified, polymerised and collected a cryo-EM dataset of FtsZ_R174D._ Despite the intentional high filament density on the grid, 2D classification could only resolve single protofilaments (Figure 2E, S7), demonstrating mutations to R174 abolish double filament formation in *E. coli* FtsZ, *in situ* and *in vitro*.

We then sought to investigate whether the doublet interface is broadly conserved. We obtained 770 different FtsZ AlphaFold2^52^ (AF2) models belonging to the phylum Pseudomonadota (formerly Proteobacteria) and filtered to less than 90% sequence identity. The remaining 119 AF FtsZ models were aligned to each side of the doublet, energy minimised and we then recorded residues within 3 Å of the neighbouring monomers (Figure 2FG). The analysis showed a highly conserved positively charged residue at either position 174 or 175, with corresponding interacting negatively charged residues on H9 (Figure 2H, S8AB). Early attempts to examine other phyla showed variability in this interface suggesting the EcFtsZ doublet interface may not represent a lateral contact for all FtsZs. Given this limitation we conclude that the doublet interface as observed with EcFtsZ is very likely conserved in related organisms and possibly even more broadly.

To test whether the observations for EcFtsZ double filaments held true outside of Pseudomonadota, we turned to a model organism historically explored by the cell division field, *Thermotoga maritima*^53,12,19^. *T. maritima* is a hyperthermophilic Gram-negative bacterium and is evolutionarily distant to *E. coli*, belonging to the phylum Thermotogota. *T. maritima* FtsZ (TmFtsZ) differs substantially from EcFtsZ, with a low sequence identity of 45% and most notably an extremely short CTT^12,19^. Like for EcFtsZ, we purified TmFtsZ and resolved to sub 4 Å resolution, a single particle cryo-EM reconstruction of TmFtsZ antiparallel double filaments (Figure 2IJ, S9). The structure revealed a highly similar double filament to that determined for EcFtsZ, with a small shift in register (less than 10 Å) between protofilaments (Figure 2K) and a ∼20° rotation along the filament long axis (Figure 2L). Although the map suffers from anisotropy, we could determine that the interface between both Tm protofilaments is bridged by R185 in the loop between H6 and H7 (Figure S9I). The main difference between the TmFtsZ and EcFtsZ double filament interface is the position of the negative charge site on H10. In Tm, the helix is positioned closer to the C-terminus in sequence and is tucked just beside H9 (Figure 2M), explaining the protofilament shift described in Figure 2K.

### *Bacillus subtilis* and bacillota FtsZ form divergent antiparallel double filaments

Having found both *E. coli* and *T. maritima* FtsZs form double filaments using a conserved arginine in the H6-H7 loop interacting with helices on the FtsZ C-terminal domain, we set out to structurally explore FtsZ filaments from a key Gram-positive model organism – *Bacillus subtilis*. Where the arginine sits at position 174 in EcFtsZ, we noted a lysine at position 175 in *B. subtilis* FtsZ (BsFtsZ), thus we anticipated a very similar double filament arrangement. Surprisingly, however, SPA of purified BsFtsZ produced 2D classes showing different double filaments to those of the *E. coli* and *T. maritima* FtsZ datasets (Figure 3A, S10A). Despite orientation preference difficulties, we could reconstruct a BsFtsZ antiparallel double filament map to ∼4 Å resolution with some anisotropy (Figure 3B, S10). The map resolution was sufficient to determine the dimerisation interface involved the FtsZ N-terminal domain (Figure 1D). We identified charged residues on H3, where E87 and E90 likely interact with K94 and K86 (Figure 3C). Despite sharing C_2_ symmetry with EcFtsZ and TmFtsZ (Figure 3D), the BsFtsZ interface constituted an almost 90° rotation in the dimer interface (Figure 3E).

**Figure 3.**
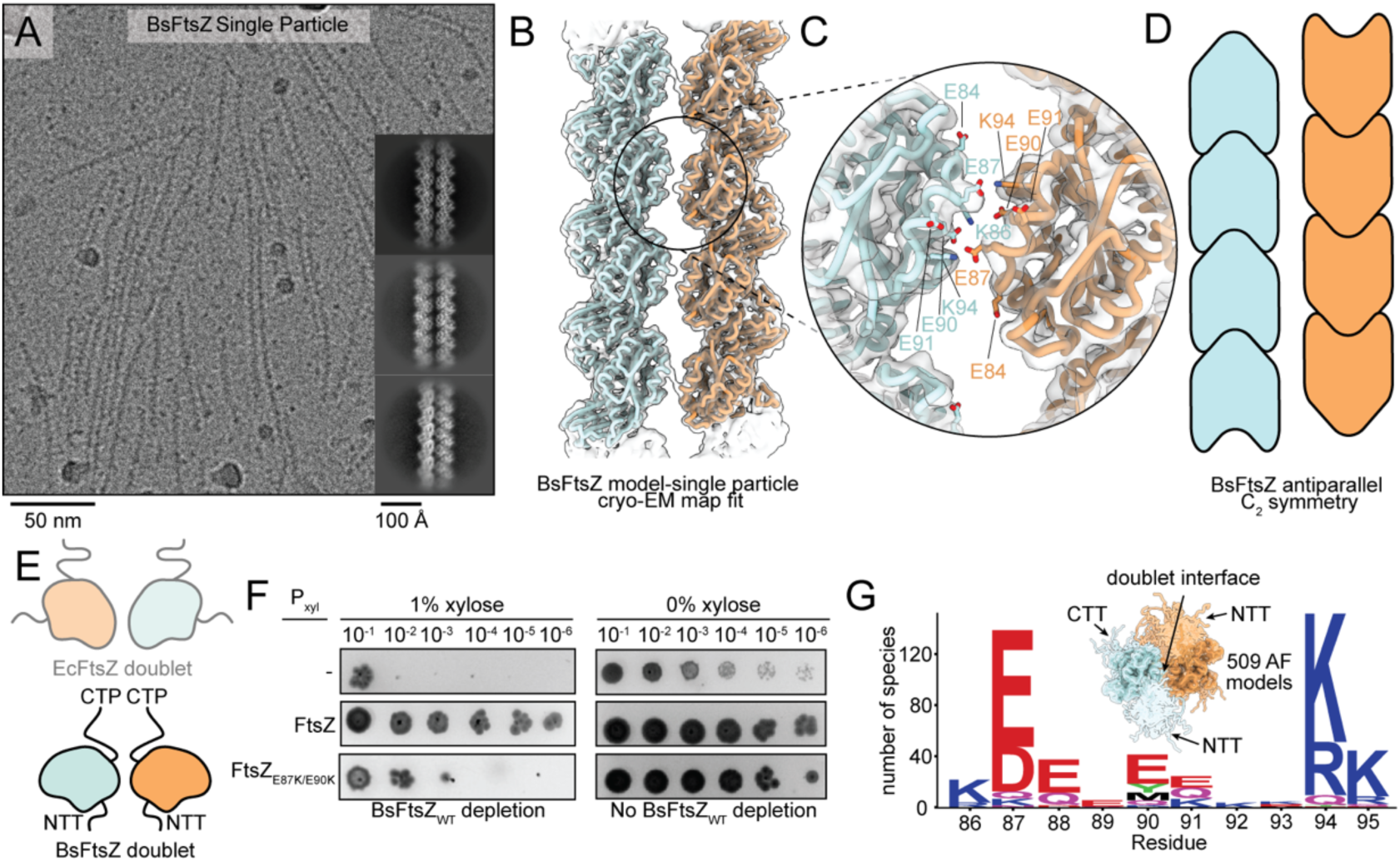
Structure and interface of FtsZ double filaments from *B. subtilis*. (A) Example micrograph and 2D classes of purified *B. subtilis* FtsZ (BsFtsZ). (B) Cryo-EM map and model of BsFtsZ double filaments. (C) Close-up of the BsFtsZ interface. (D) Schematic of double filament orientation and C_2_ symmetry. (E) Schematic of BsFtsZ doublet top view vs EcFtsZ. (F) Spot dilution assay of FtsZ depletion and complementation in *B. subtilis* using a dCas9 knockdown system for WT FtsZ. (G) AF2 model analysis of BsFtsZ doublet interface where AF2 monomers from Bacillota were aligned to both sides of the doublet interface.

To test this interface, we used a xylose inducible dCas9 knockdown system^54^, specifically targeting WT FtsZ (Figure S11AB). Spot dilution assays (Figure 3F) showed a loss of viability when grown in 1% xylose versus no xylose. This was recovered by incorporating a second dCas9-immune *ftsZ* copy at the amyE locus under a xylose inducible promoter. In contrast, WT FtsZ depletion could not be recovered with a E87K/E90K double charge reversal. Imaging performed on this strain also showed elongated cells compared to controls (Figure S11CD), indicative of a cell division phenotype. In agreement with this result, we performed other spot dilution experiments as part of our live cell imaging discussed later (Figure S19B) that show a strong dominant negative effect when the FtsZ mutant is overexpressed. Unlike the R174 mutations in EcFtsZ, it is likely the E87K/E90K mutant increased the lateral association of protofilaments as negative stain EM showed substantial bundling of filaments in the presence of GTP (Figure S12AB vs. S12EF), while pelleting assays revealed increased pelleting with the non-hydrolysable GTP analogue GMPCPP (Figure S12G). Therefore, we conclude the N-terminal lateral interface observed between BsFtsZ protofilaments, while different from EcFtsZ and TmFtsZ C-terminal interfaces, is important for viability in *B. subtilis*.

Given the divergence in interface between *E. coli* and *B. subtilis*, we wanted to test if its doublet interface is conserved within the phylum of Bacillota (formerly known as Firmicutes). Like with the *E. coli* interface, we aligned AF2 models (filtered to less than 90% identity) to the doublet interface to extract conserved features. Our analysis proposed a strongly conserved interaction pair with a negative charge at position 87 and a positive charge at position 94 (Figure 3G, S8CD), suggesting these residues are important for maintaining the FtsZ lateral interface in many species. Therefore, we conclude BsFtsZ protofilaments can assemble into double filaments by an N-terminal interface that is likely widely conserved across the phylum Bacillota.

### The NTT regulates EcFtsZ doublet-doublet distances

While processing the EcFtsZ_D212A_ overexpression tomograms for STA in *E. coli*, we noticed that the doublets appeared to self-align with respect to each other and to the membrane (Figure 1C). These bands of aligned filaments resemble previous cryo-ET results of overexpressed EcFtsZ in *E. coli* and in WT *Caulobacter cresentus*^19^. Similarly, processing of the SPA cryo-EM EcFtsZ dataset, obtained by imaging purified EcFtsZ, revealed 2D classes with neighbouring double filaments (example 2D class in Figure S3B). In both cases, expanding the refinement box revealed a doublet-doublet arrangement that at low resolutions (∼16 Å for STA and ∼6 Å for SPA) appeared consistent between both *in situ* and *in vitro* SPA datasets, with a small difference in the inter-doublet distances (Figure 4A). From the in-cell STA analysis, we knew most EcFtsZ filaments oriented their CTT towards the membrane (Figure 1H). Additionally, we knew doublets were maintained by very specific interactions involving the globular FtsZ C-terminal domain (Figure 1D). This positions the N-terminal domains of EcFtsZ in the direction of neighbouring doublets. With the doublet-doublet spacing being too far apart from each other to be mediated by any residues resolved in our structures, we learned that this doublet-doublet arrangement must involve the unresolved N-terminal tail (NTT) (Figure 1D, 4B). This would mean EcFtsZ filaments have two interfaces, the doublet mediated by C-terminal domain residues (C-C interface) and another N-terminal (N-N interface).

As with the C-C interface (EcFtsZ doublet), we used the temperature sensitive EcFtsZ depletion *E. coli* strain to probe for NTT mutants resulting in strong growth phenotypes (Figure 4C, S13). As shown previously^55^, deletion of the NTT including Δ2-10 abolished growth in the absence of WT EcFtsZ, as did a Δ2-9 mutant. Given the structure of EcFtsZ, we were concerned a deletion including aspartate 10 may have impacted the protofilament longitudinal contact and so made a half NTT deletion (Δ2-6), which showed a similarly strong phenotype. However, deleting one residue less (Δ2-5) showed no growth defect or division phenotype in our microscopy images (Figure S14). Looking at the conservation of the NTT within Pseudomonadota (Figure 4C, top), we noted that aside from the highly conserved F2, negative charges at position 3 and 6 appeared to be conserved. But removing charge from either of these in EcFtsZ showed no clear phenotype in our spot assay (Figure 4C, S13) unless both positions were mutated to alanine simultaneously. In addition, replacing residues 2-9 of the NTT with a putatively disordered region from *E. coli* FtsN (residues 142-149), which happens to have similarly placed negative charges (positions 143 and 147) appears to complement while mutating these charges to alanine (charge removal) does not. We also tested residues 151-159 from *E. coli* FtsN between positions 2-9 that had no negative charges but an abundance of polar residues. We saw a clear division phenotype when imaged (Figure S14), despite a spot assay showing viability, one of two discrepancies between the two assays in our study (FtsZ_2/FtsN151-159/9_ in Figure S13 and S14 and FtsZ_G316/FtsN140-189/K367_ in S6 and S15D). From these data, we conclude the NTT is important for EcFtsZ function. Since it keeps neighbouring doublets together and at a distance, as part of an N-N interface, an intrinsically-disordered, brush-like role for the NTT is most likely.

We also looked at mutations of the much longer CTT, as we cannot formally rule out a role for it in lateral interactions. The results are shown in Figures S6 and S15 but, to summarise, we observed a tolerated CTT replacement with residues 140-189 of FtsN that was temperature sensitive (tolerated at 37 °C but not 42 °C, Figure S15D). Considering the CTT does not point towards other protofilaments laterally and previous work showing tolerated CTT replacements^56^, we focussed our efforts on the NTT.

**Figure 4.**
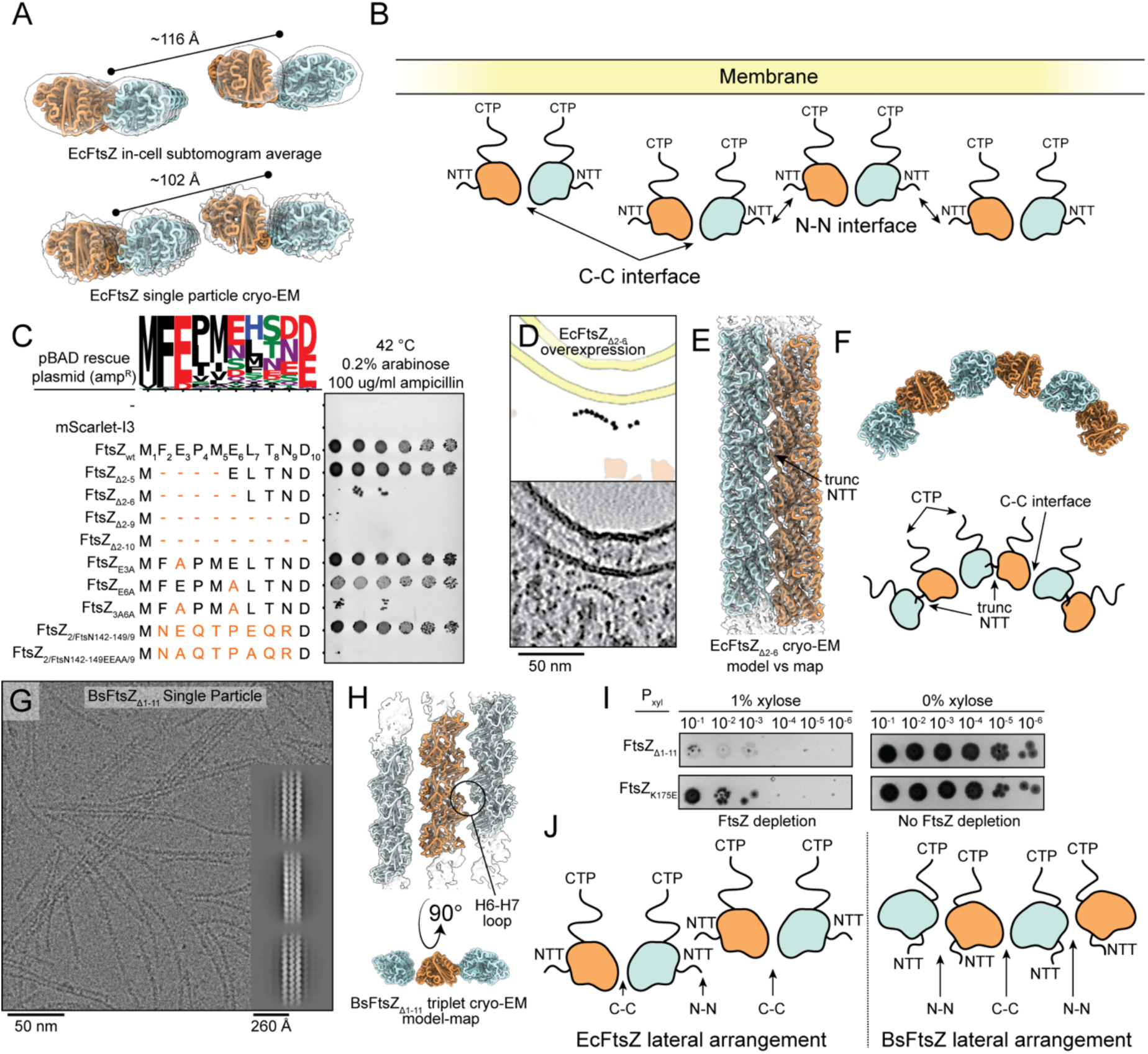
The second FtsZ lateral interface. (A) EcFtsZ doublet-doublet density from both *in situ* and *in vitro* (purified FtsZ) datasets. The EcFtsZ doublet model was docked into both reconstructions. (B) Schematic of laterally assembled doublets in the EcFtsZ *in situ* tomograms (Figure 1A, C), with the orientations of the disordered NTT and CTT labelled. (C) Spot dilution assay of *E. coli* modified to enable temperature sensitive FtsZ depletion, complemented with FtsZ N-terminal mutants and an mScarlet-I3 control in a pBAD vector. Top is a weblogo of a multiple sequence alignment of the NTT from organisms used in the AF2 analysis shown in Figure 2H-J. (D) Tomogram and segmentation 2D slice of overexpressed FtsZ_Δ2-6_ in *E. coli* C41(DE3). (E) Doublet cryo-EM map and model of the EcFtsZ_Δ2-6_ double filament, showing a different and tight interface. (F) Recapitulating the FtsZ arc shown in Figure 4D by combining both N-terminal EcFtsZ_Δ2-6_ interface and the WT doublet C-terminal interface. Top is the model from combining both interfaces and bottom is a schematic describing the model. (G) Representative micrograph and 2D classes of purified BsFtsZ_Δ1-11_ showing extensive lateral interactions. (H) Cryo-EM map and model derived using purified BsFtsZ_Δ1-11_, revealing an additional, second C-terminal interface between BsFtsZ protofilaments. (I) Spot dilution assay of *B. subtilis* dCas9 BsFtsZ depletion strain complemented with K175E at the Bs C-terminal interface. (J) Schematic comparing the arrangement of the two EcFtsZ and BsFtsZ lateral interactions. Different arrangements in *E. coli* and *B. subtilis* produce similar overall bands of FtsZ doublets as shown in Figure 1AC. The interactions producing these can be grouped into C-C and N-N interfaces, but their details are different.

Knowing that the NTT deletions were non-viable in the temperature sensitive *E. coli* FtsZ depletion (Figure 4C), we decided to look at the partial deletion (Δ2-6) when overexpressed *in situ* using cryo-ET. Surprisingly, this revealed protofilaments assembled into membrane-proximal arcs (Figure 4D), distinct from other *in situ* observations of FtsZ. To further characterise this assembly, we determined the structure of EcFtsZ_Δ2-6_ double filaments using single particle cryo-EM to 2.7 Å, revealing a different, tighter interface mediated by the remaining NTT (Figure 4E, Figure S16I). The *in situ* arcs (Figure 4D) could not be observed with purified EcFtsZ_Δ2-6_ on EM grids, possibly due to their shape. Despite this, we found we could recapitulate the *in situ* observations by considering the WT C-C interface in addition to the new EcFtsZ_Δ2-6_ N-N interface. Including both lateral interfaces in a model produced an arc closely resembling that seen *in situ* (Figure 4F). Taken together, these results suggest that the NTT likely generates brush-like inter-doublet interactions (N-N interface).

We also investigated the longer Δ2-10 NTT deletion *in situ* with cryo-ET and found protofilaments bundled near the membrane but now reoriented along the long axis of the cell (Movie S5). Similar to Δ2-6, we purified and determined the structure of EcFtsZ_Δ2-10_ filaments (Figure S17), revealing a hollow cylindrical structure that is composed of antiparallel protofilaments (Figure S17I), with lateral contacts mediated by the WT EcFtsZ doublet interface and the new truncated N-terminal face mediated by a polar contact between R50 and T57 (Figure S17J). We noted that despite the similarity to other cylindrical hollow tubulin assemblies such as microtubules, this EcFtsZ_Δ2-10_ structure differs substantially due to its antiparallel arrangement. This confirmed to us the NTT plays an important role in regulating N-N lateral interfaces between EcFtsZ protofilaments while also demonstrating the diversity of assemblies that can be generated from minimal changes to protein sequence.

### BsFtsZ also has a second lateral interface

With compelling evidence for a second lateral interface in EcFtsZ (N-N interface) required for band formation, we turned our attention back to *B. subtilis*. As shown, the doublet interface for BsFtsZ is an N-N interface (Figure 3), different from the C-C interface of EcFtsZ. To test the effects of an NTT deletion on BsFtsZ protofilaments, we determined the structure of BsFtsZ_Δ1-11_ using cryo-EM. This construct showed an increase in laterally assembled/bundled protofilaments when polymerised with GTP (Figure 4G), in line with pelleting assays showing a similar propensity for BsFtsZ_Δ1-11_ to pellet with GTP as with the non-hydrolysable GMPCPP, unlike for BsFtsZ_WT_ (Figure S12G). This suggested that the NTT also destabilised N-N lateral contacts in *B. subtilis*, despite the interface consisting of residue-residue contacts.

We selected for minority particles within this dataset and classified into 2D classes with three protofilaments (Figure 4G). We refined these particles further in 3D and although the data did not yield high resolution, we could confidently dock models into the three protofilament densities (Figure 4H). This revealed that the protofilaments were again antiparallel with their CTTs pointing in the same general direction while a similar interface to the original BsFtsZ N-N lateral interface (Figure 3BC) was preserved. It appeared that deleting the NTT allowed a more stable N-N lateral interface to scaffold an additional C-C lateral interface, akin to that for EcFtsZ (Figure 4BH). As for EcFtsZ, the C-C interface in BsFtsZ likely involves residues from the H6-7 loop but this time are positioned near the H6-7 loop from the opposite protofilament (Figure 4H). Resolution limitations meant we could not resolve exact residues involved but we suggest residues D174, K175 and N176 are probable candidates.

We probed the importance of the BsFtsZ NTT for *B. subtilis* viability in the dCas9 FtsZ knockdown system, showing no spot growth when BsFtsZ_Δ1-11_ was the only FtsZ copy expressed (Figure 4I), while imaging revealed this strain had an elongated cell phenotype (Figure S11D). We also tested the BsFtsZ C-C interface with a charge reversal of K175 (K175E), the BsFtsZ equivalent of the R174 residue involved in the EcFtsZ C-C interface. This also revealed reduced cell viability (Figure 4I) and a curved cell phenotype (Figure S11D) suggesting K175 likely plays a role in BsFtsZ lateral interactions, like for R174 in EcFtsZ, but is unlikely to be the dominant interface. It became clear that the specific structural differences between EcFtsZ and BsFtsZ lateral interactions were obscuring conserved general properties of FtsZ lateral interactions (Figure 4J), since quite similar bands of antiparallel double filaments can be built in both cases. Each require two different lateral interactions (because of the symmetry of each doublet being “closed”) that we named N-N and C-C and together they condense FtsZ protofilaments into bands, the Z-ring.

### Lateral interactions impact treadmilling rates and Z-ring properties

Our work to this point showed snapshots of Z-ring and FtsZ protofilament self-assemblies but was limited by the static nature of the techniques used. FtsZ exhibits dynamic GTP-driven behaviour known as treadmilling and so we sought to test the relevance of this kinetic phenomenon, factoring for the lateral interactions we had discovered, for Z-ring condensation. We developed a coarse-grained (CG) description of FtsZ protofilaments for molecular dynamics (MD) simulations, adapted from a previously published model^32^. FtsZ monomer lifetimes were modelled on experimental *in vivo* single molecule data (Figure S18)^39^ and were designed to include patches used to model the identified lateral interactions based on the *E. coli* C-C (interface 1) and N-N (interface 2) interfaces (Figure 5A). In addition, we hypothesised that when FtsZ protofilaments were engaged in a C-C lateral interaction (through interface 1), the depolymerisation rate should decrease compared to single protofilaments. This is due to two bonds being broken through depolymerisation if the end monomer has a formed C-C interface, compared to one bond otherwise. We term this additional condition on the treadmilling rates “kinetics modulation”, inspired by the “cozy corner” concept from microtubules^57^. We simulated Z-ring condensation as previously established^32^ for a period of 30 minutes, and saw that the modulation of treadmilling kinetics significantly promoted the formation of condensed, co-linear filaments as they are observed in the Z-ring by tomography (Figure 5B). In contrast, without kinetics modulation we saw no condensation.

**Figure 5.**
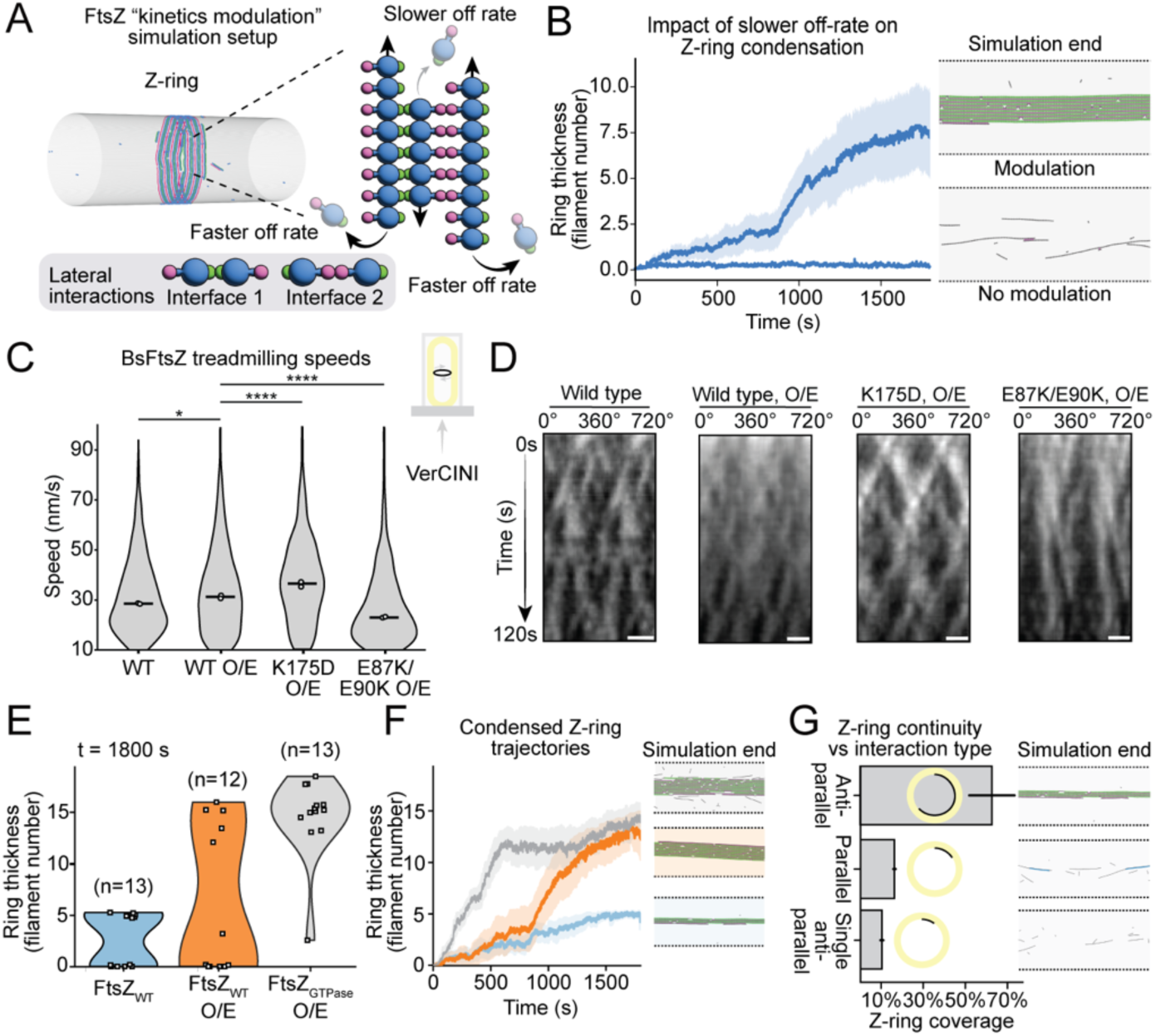
MD simulation and live cell imaging studies on the relationship between lateral interactions and treadmilling. (A) Schematic of the coarse-grained MD simulation models designed to impose two antiparallel lateral interactions. The model assumes a slower off rate of FtsZ subunits when engaged in lateral interactions vs subunits with no lateral interactions. (B) Proof of model concept comparing simulations with modulated off-rates and simulations without. Simulations are compared based on their Z-ring thicknesses, with a representative simulation end snapshot shown on the right-hand side for both conditions. (C) Violin plots of FtsZ mutant treadmilling speeds vs WT/WT overexpression, derived from live cell imaging of *B. subtilis* cells in VerCINI microholes. Speeds below 10 nm/s were excluded. Circles represent the median of both biological replicates and pairwise significance was calculated using the Mann-Whitney U test. See Table S8 for p-values. (D) Representative kymographs of the live cell imaging datasets used to calculate FtsZ treadmilling speeds (scale bar 1 μm). Kymographs were plotted as two complete revolutions around the cell circumference (0-720°) in order to resolve filaments that pass the 0°/360° boundary. (E) Violin plots comparing the effect of FtsZ concentration and GTP hydrolysis rate on Z-ring width (thickness) in MD simulations. (F) Z-ring thicknessed over the course of simulation trajectories for the conditions in panel E. Only repeats resulting in a final condensed Z-ring are plotted. Trajectories and simulation end point snapshots are colour coded as in panel E. (G) In MD simulations, the total length of continuous lateral interactions reached with two antiparallel lateral interactions (62.58%) compared with simulations enforcing parallel interactions only (16.1%) and only one antiparallel interface (10.1%). Lengths are represented as percentages of simulation box widths (equivalent to % septal circumference). Schematics to the right of the box plots illustrate the relative septal circumference (yellow) covered by Z-ring lateral interactions (black).

To verify *in vivo* whether treadmilling speeds are impacted by lateral interactions as suggested by our simulations, we used live-cell imaging in *B. subtilis* to measure treadmilling of BsFtsZ in cells that were vertically trapped in VerCINI microholes^58^ as described previously^24^. We compared treadmilling speeds of WT BsFtsZ and overexpression of either WT or mutants that should affect BsFtsZ lateral interfaces (K175D and E87K/E90K). We postulated that a charge reversal of BsFtsZ residue K175 (from positive to negative) would perturb C-C lateral interactions similarly to the *E. coli* R174 mutant, while negative stain EM and pelleting assays (Figure S12E-G) demonstrated that adding two new positive charges (E87K/E90K) likely enhances BsFtsZ lateral interactions. Imaging experiments revealed populations of both mobile and immobile treadmilling filaments, as reported before^31^, whilst overexpression of either BsFtsZ interface mutants was dominant negative over 24 h (Figure S19B). Examination of the mobile treadmilling filaments (speeds above 10 nm/s) showed a statistically significant increase in treadmilling speed for the BsFtsZ_K175D_ overexpression (lateral disruptor) versus the WT BsFtsZ overexpression, while BsFtsZ_E87K/E90K_ overexpression (lateral enhancer) had a substantially slowed treadmilling speed (Figure 5C-D). That lateral interactions impact treadmilling speeds validated the kinetics modulation regime in our MD simulations (Figure 5B).

We next explored the effects of increased FtsZ levels and treadmilling arrest on Z-ring width using the CG MD simulation setup. We modelled Z ring condensation for 30 mins with WT levels of FtsZ compared to FtsZ and FtsZ_GTPase mutant_ at increased levels. In the WT setup, we limited the maximum number of monomers to a physiologically relevant concentration of 4,000 copies^32^, while in the cases with increased FtsZ levels (“FtsZ overexpression”) there was no limit to the number of monomers in the system. To model the GTPase mutant, we slowed down hydrolysis by a factor of 10 in the simulations. These simulations showed substantial differences between the three conditions (Figure 5E). Our “GTPase mutant” and WT overexpression conditions, when condensed, resulted in similarly thick Z-rings and substantially thicker than those for condensed WT Z-rings. However, both the WT and WT overexpression conditions behaved much more stochastically with many simulations resulting in no condensed Z-rings at all. This produced a bimodal spread of Z-ring widths, with values at either 0 or maximally condensed. To compare condensed Z-rings, we separated trajectories that had a width of at least 0.5 filament widths, from those that did not condense (9/15 WT, 6/12 WT overexpression and 13/13 GTPase mutant overexpression) and looked at their mean Z-rings widths over the course of the simulation trajectories (Figure 5F). This analysis showed that while both GTPase mutant and WT overexpression simulations condensed to similar Z-ring widths, 3x the width of WT rings, the GTPase mutant condensed quicker on average suggesting treadmilling inherently opposes early Z-ring condensation.

A major finding of our work thus far is the antiparallel orientation of protofilaments in lateral FtsZ interactions. With other lateral assemblies of tubulins consisting of parallel protofilaments (e.g. prokaryotic and eukaryotic microtubules), we wanted to test the mechanistic importance of having antiparallel treadmilling protofilaments using our simulations. We setup three simulation regimes, each with different lateral interaction types: antiparallel with two interfaces, parallel with two interfaces and antiparallel with only one lateral interface (only the interface 1 i.e. where a double filament is unable to interact with other doublets). We performed Z-ring condensation simulations for periods of 30 mins, tracking the circumferential coverage of the Z ring. We observed substantially different outcomes for the first condition, antiparallel with two interfaces, compared to the other two scenarios (Movie S6). By approximating Z-ring continuity as the length of the longest stretch of lateral interacting filaments, we revealed that only antiparallel filaments with two lateral interfaces resulted in highly continuous mid-cell circumference coverage (62.58% of the box length vs ∼26.1% and ∼10.1%; Figure 5G). We conclude that antiparallel arrangements with two lateral interfaces are important for generating a denser and more continuous Z-ring.

## Discussion

The question of how protofilaments condense into a Z-ring is at the heart of our understanding of FtsZ-based bacterial cell division. In this study, we have provided *in situ* and *in vitro* evidence that FtsZ protofilaments can self-assemble through defined lateral interactions and arrange themselves antiparallel with respect to each other. We have used *in situ* subtomogram averaging (Figure 1E) and cryo-ET to validate the *in vitro* work, conclusively showing the orientation of filaments with respect to each other and the membrane (Figure 1H). This finding unites FtsZ with its other tubulin homologues, which all have defined lateral interfaces used to assemble higher order filament structures^59^, such as, but not limited to, prokaryotic^60^ and eukaryotic microtubules^61,62,63^. Our *E. coli* double filament structure suggests compelling structural links to eukaryotic tubulins where the position of the tubulin M-loop protrudes from the helix equivalent to the EcFtsZ C-terminal helix 9 (H9) that provides the negative charges at the doublet interface (Figure S20). This may provide clues to the evolutionary pathway from an FtsZ-like last common ancestor to microtubules, although it is important to point out that FtsZ is currently unique in its propensity to form antiparallel protofilament interfaces, a situation that is not completely dissimilar to MreB and FtsA, prokaryotic actin-like proteins that also form antiparallel filaments in contrast to all other actins^14,23,64,65,66^.

**Figure 6.**
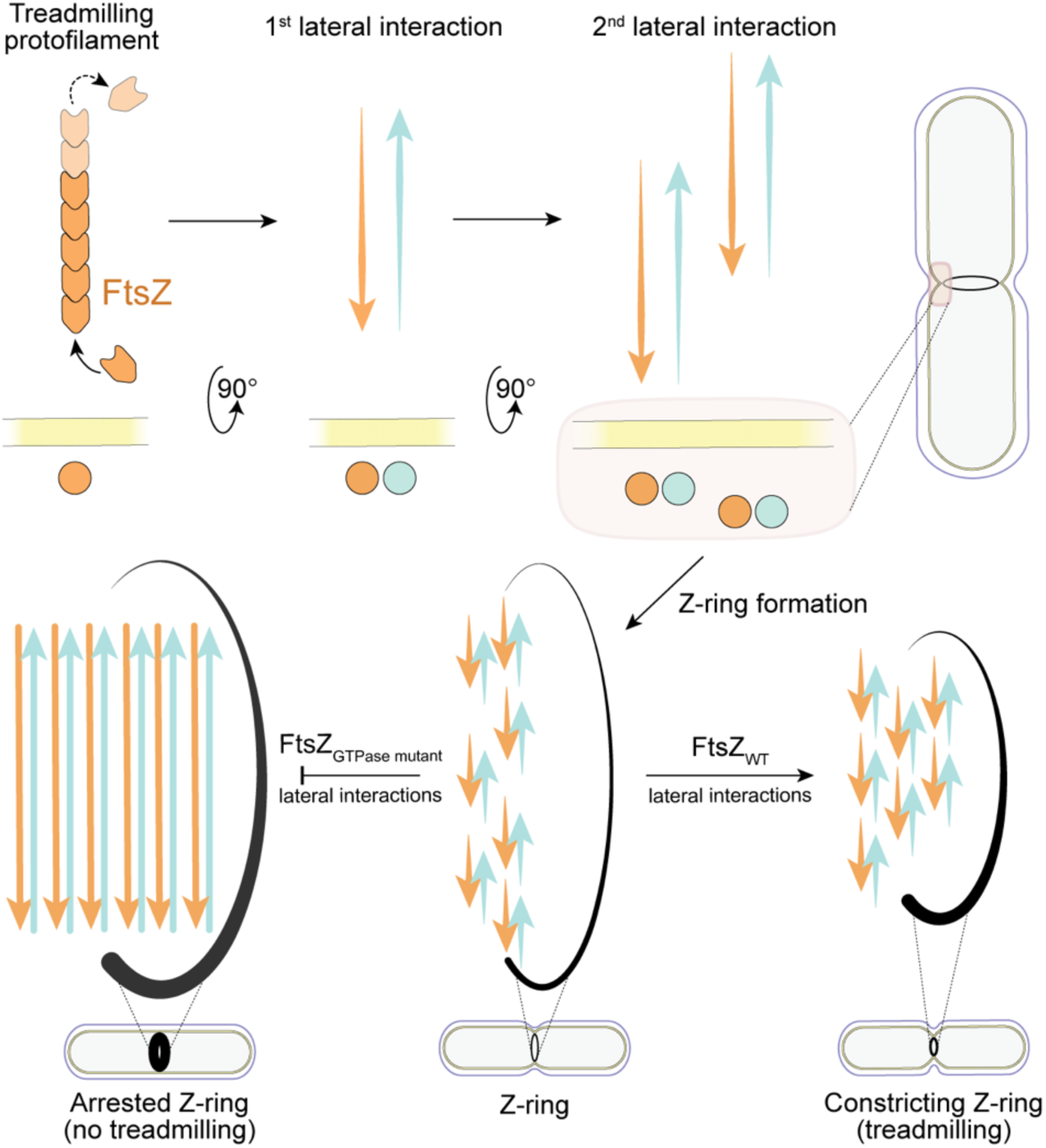
A model for the condensation of FtsZ protofilaments into the Z-ring and the role of treadmilling. Top left to right: a treadmilling FtsZ protofilament engaging in first one (dominant) and a second lateral interaction, leading to two different types of interactions. This allows the treadmilling protofilaments to both extend longitudinally and associate laterally (open symmetry), resulting in an increasingly condensed Z-ring during early division (bottom middle) as more subunits and filaments get pulled in through increasing avidity. Since evidence indicates that septal cell wall synthesis, driven from the outside of the cell by PG synthases, is the primary force driving septal constriction, the Z-ring must track and itself constrict accordingly on the inside of the cell. We propose that this at least requires filament dynamics (pushed to a smaller diameter) but optimally functions if the Z-ring is self-tensioning (pulling to a smaller diameter), as is indicated by the fact the FtsZ filaments are located away from the membrane, despite being anchored to the membrane via the long and disordered CTT. During early division the mid-cell diameter is largest, FtsZ protofilament lateral interactions are determined by how many lateral interactions such a diameter can produce. Progressing through division, the number of lateral interactions increases as the diameter shrinks and the number of FtsZ subunits stays constant (bottom right). Given lateral interactions produce binding energy, the lateral interactions can therefore drive energetically Z-ring tensioning towards smaller diameters as protofilaments maximise their lateral contacts. If treadmilling is not present or disabled (bottom left), all possible lateral interactions will be satisfied (maximal condensation without the ability to remodel or shrink), which produces a static Z-ring. Thus, both lateral interactions and treadmilling are needed to build a Z-ring and reduce its diameter as the cell constricts.

Our FtsZ structures determined here from three different organisms, spread across major clades of the prokaryotic tree of life, suggest the antiparallel arrangement is a broadly conserved feature of FtsZ-based cell division in bacteria. As indicated by the strong, dominant negative growth defects produced by FtsZ interface mutants, it is highly likely that this antiparallel arrangement is essential to FtsZ’s role in cell division. But why? The first answer lies in the fact that the direction of the C_2_ symmetry we discovered is the only one that makes sure the CTTs of all FtsZ filaments point towards the cell membrane. This symmetry is “closed” and requires a second lateral interface to form bands and a Z-ring, which we found in both *E. coli* and *B. subtilis* FtsZs (Figure 4). Filaments could also have been parallel (translational symmetry only) and not treadmill past each other as they do in the antiparallel case. However, our simulations summarised in Figure 5 demonstrate that the antiparallel nature of the filaments significantly helps with Z-ring formation and this we believe can be intuitively understood since treadmilling of protofilaments treadmilling past each other increases the chances of them to overlap, to form lateral interfaces.

The other main finding from our simulations and live-cell imaging was the impact of lateral interactions on treadmilling itself. Our experimental data and simulations suggest that lateral interactions reduce subunit off rates in treadmilling filaments (Figure 5BCD). We borrowed this idea from the “cozy corner” concept in microtubules^57^, where the increased binding surface for a doublet (or laterally interacting tubulin) probably slows subunit depolymerisation compared to a subunit in a singlet. It can thus be expected that this treadmilling modulation serves as both a Z-ring condensing and extending mechanism: the lateral interactions drive Z-ring condensation, while treadmilling ensures only laterally associated protofilaments reach longer filament lengths. This will make filaments appear more in the Z-ring and not elsewhere in the cell.

Considering the relationship between FtsZ lateral interactions and treadmilling points to another important Z-ring mechanism (Figure 6). We have proposed lateral interactions can slow subunit off-rates, but the antiparallel association of protofilaments means treadmilling ultimately opposes lateral interactions by causing protofilaments to move away from one another. These protofilaments can continue treadmilling and encounter other protofilaments resulting in new lateral interactions. A means of achieving more lateral interactions, which must be energetically favoured, would be to shrink the diameter of the Z-ring, reducing the surface area FtsZ protofilaments can occupy. The result would be more protofilament lateral contacts meaning lateral interactions could drive Z-ring constriction because forming more lateral contacts would be energetically favoured. But this cannot be driven by lateral interactions alone. Without treadmilling, protofilaments can maximise lateral interactions without constriction, making treadmilling essential for guiding protofilaments towards smaller Z-ring diameters. Without other mechanisms such as sliding^67^, which would be energetically unfavourable, it is difficult to imagine how the Z-ring can shrink in the absence of treadmilling, due to the stabilising effect of lateral protofilament interactions.

What could this “tensioning” of the Z-ring be good for? It is clear that recent studies showing septal closure with perturbed FtsZ treadmilling strongly favour a synthase-driven constriction model of the cell^29,26–28^, in which PG synthesis on the outside pushes the cell envelope inwards. But this means the Z-ring on the inside will have to get smaller simultaneously. A role for the proposed Z-ring tensioning, driven by lateral interactions and treadmilling, would be to make sure the Z-ring always tracks the cell diameter as it gets smaller. It is even possible that Z-ring tensioning represents an evolutionary relic from a cell wall-less ancestor with low internal turgor pressure^68^. In this context it is also important to point out that some bacteria and many archaea use FtsZ for cell division but do not have a cell wall and it has been suggested in the case of *Haloferax volcanii* that FtsZ might have a more active role in cell constriction^69^.

We found in both the *E. coli* and *B. subtilis* cases that their FtsZs form doublets through different, dominant lateral interfaces, that are supplemented by second, weaker ones. Datasets for purified TmFtsZ presented more sparse filament distributions and without *in situ* investigation, we could not define a similar second interface for this FtsZ.

Residues forming the dominant EcFtsZ doublet have been mentioned previously. We found mentions of the R174D mutation as early as 2003, linking the residue to protofilament bundling^50^. Moreover, the adjacent mutant L169R appears to have a bundling phenotype and can bypass the typically essential ZipA^4,35^, while also rescuing the dominant negative phenotype of FtsZ_R174D_^33,70^. Encouragingly, evolution seems to have targeted this interface through toxin-antitoxin systems. Tre1 is secreted by the type VI secretion system of *Serratia proteamaculans* and can ADP-ribosylate R174 of FtsZ in *E. coli*^71^, while CbtA appears to bind the same H6-7 loop R174 is positioned on^72^, resulting in cell division inhibition and cell lysis^73^. The targeting of this lateral interface by toxins suggests wider conservation within Gammaproteobacteria and possibly beyond. The likeness of our TmFtsZ double filament structure to EcFtsZ points towards this possibility.

The MinCDE system is now well established as a protector of the cell poles from FtsZ polymerisation (the old divisions sites) and also a mid-cell positioner for the Z-ring^74^. Interestingly, the C-terminus of MinC has previously been reported to disrupt FtsZ protofilament lateral interactions^75^ and a recent study reports that MinCD can remove ZapA from the Z-ring in *B. subtilis*^76^. With no structural view of the MinC-FtsZ interaction this remains speculative but preventing or disrupting FtsZ lateral interactions would be a good way of ensuring Z-rings can form only where the MinCDE system allows it.

While the general principles of FtsZ lateral interactions are shared between *E. coli* and *B. subtilis*, the specifics of their lateral interfaces are diverged. We have shown that the dominant lateral interface in BsFtsZ occurs at the N-terminal domain as opposed to a more C-terminal interface in EcFtsZ (Figure 3E). While we lack the resolution to point at specific interactions in the second C-C interface in BsFtsZ, it is clear the same H6-7 loop is involved and likely self-interacts across the protofilament interface. With the findings that the bacterial last common ancestor was likely a diderm^77^, it is possible that the C-C interface in EcFtsZ is closer to a more ancient lateral interface which eventually split and gave rise to modern FtsZ and tubulins (assembled by an M-loop insertion). The BsFtsZ lateral interfaces would then represent a divergence potentially to adapt to different cellular characteristics such as higher turgor pressure^68,78^ and higher ordering required to match the negative curvature of an invaginating Gram positive septum^20^. This may explain the BsFtsZ N-N interface having robust and resolvable residue-residue contacts that are not present in the equivalent EcFtsZ interface.

Our investigation of the N-N interface in EcFtsZ highlighted that we cannot easily resolve or understand interfaces involving intrinsically disordered regions (IDRs). The NTT, recently proposed to destabilise longitudinal contacts^55^, negatively regulates lateral interactions in both organisms (Figure 4, S16), but we should not rule out an attractive contribution in EcFtsZ either since we do know that the filaments form doublets of doublets *in vitro*, with no other factors present. Another difficulty is that FtsZ has another disordered region, the CTT, that could be involved exclusively or additionally in the second *E. coli* interface. Indeed, the much longer CTT has been implicated in lateral interactions previously^33,36,79,80^, but it is worth mentioning that replacing the CTT with an unrelated sequence of the same length is viable, making it perhaps less likely to be involved in specific interactions^56^. Much like the NTT, the CTT could also act as a polymer-brush and our preliminary mutagenesis studies do not rule this out completely (Figure S15). Further studies will be needed to completely explain the contributions of both IDRs to FtsZ lateral interactions.

A recent study revealed tetrameric ZapA bound to antiparallel double filaments of FtsZ from *Klebsiella pneumoniae* (KpFtsZ)^43^. KpFtsZ and EcFtsZ are highly similar, differing by only five residues. The KpFtsZ-ZapA doublet arrangement appears related to the one reported here (Figure 1) at a first glance, but aligning to one protofilament reveals a substantial ∼20 Å (1/2 subunit) shift in both the filament-filament register and a 60-70° rotation around the C-C interface (Figure S21AB). Additionally, the R174-H9 contact is broken, making it likely this conformation is dependent on the ZapA interaction. Our work has revealed specific FtsZ-FtsZ interactions in cells are not dependent on additional proteins as the same contacts occur with purified FtsZ. Therefore, we suggest the roles of such crosslinking proteins are not to facilitate necessary FtsZ-FtsZ lateral interactions but to modify them. This becomes clearer when considering ZapA appears to contact the NTT, drawing it away from the N-N lateral interface (Figure S21BF). Given ZapA is not essential and only produces a growth phenotype when overexpressed^40,81–83^, it is possible ZapA is enhancing or weakening the N-N interface through sequestering the NTT. Similarly, the recently reported toroids made by FtsZ upon addition of ZapD^44^ point towards a modified FtsZ-FtsZ lateral interface. Our work does not contradict such findings on the effects of crosslinking proteins but reveals the basic self-assembly mechanisms onto which FtsZ-binding proteins likely add additional layers of regulation. How each of these proteins achieves this remains to be investigated further.

In summary, we have discovered conserved fundamental properties of how FtsZ protofilaments condense in an antiparallel fashion into a Z-ring by determining the in-cell structure of FtsZ filaments from *E. coli and* recapitulating the interfaces they form with purified FtsZ, *in vitro*. We have validated the physiological relevance of the filament interfaces with genetics and phenotyping. We have tested generality by demonstrating that other bacterial Z-rings likely use partially conserved interfaces between FtsZ protofilaments to form. We have established the importance of antiparallel protofilaments through coarse-grained molecular dynamics simulations and have started to investigate the emergent properties of the system, for example with regards to the modulation of treadmilling dynamics.

Our work positions FtsZ as the building block of a highly self-organising system that orchestrates bacterial cell division. Our findings pave the way towards full understanding of Z-ring condensation and its regulation, potential artificial biology applications and propose a compelling model for Z-ring constriction.

## Resource Availability

The in-cell EcFtsZ STA map was deposited in the Electron Microscopy Data Bank (EMDB) with accession number EMD-100280. EcFtsZ_WT_, EcFtsZ_Δ2-6_ and EcFtsZ_Δ2-10_ cryo-EM structures were deposited in the EMDB under accessions EMD-100276, EMD-100278 and EMD-100277 and the Protein Data Bank (PDB) under accessions 34YW, 34YZ, 34YY respectively. The TmFtsZ cryo-EM structure was deposited in the EMDB under accession EMD-100783 and PDB 50MZ. The BsFtsZ cryo-EM structure was deposited in the EMDB under accession EMD-100880 and PDB 50SS. All code used to generate MD simulations can be found in a GitHub repository at https://github.com/Saric-Group/Doublets.

## Acknowledgements

We thank all members of the MRC Laboratory of Molecular Biology (LMB) Electron Microscopy Facility for help and support with data collection, Scientific Computing for their support with data processing and Mass Spectrometry Facility, in particular Tomos Morgan for purified sample analysis at the LMB. We thank past Löwe group members Victoria Hale and James Hooker for their early efforts towards this project’s aims and for sharing their knowledge within the group. We thank Felix Wodaczek (ISTA) for his help with MD simulations. This work was funded by the Medical Research Council, UK (U105184326 to JL), the Wellcome Trust (227876/Z/23/Z to JL and 227452/Z/23/Z to SH, JL and AS). GC, AŠ and AW were supported in part by grants from the Agence Nationale de la Recherche (ANR-24-CE11-4058 and ANR-21-CE20-0040 (to AW)) and by institutional funding from the Institut Pasteur, the CNRS, and Université Paris Cité. They gratefully acknowledge the NCF, UBI and HPC core facilities at Institut Pasteur. LR was funded by an EMBO postdoctoral fellowship (EMBO ALTF 88-2023) and MK acknowledges funding by the Gesellschaft für Forschungsförderung Niederösterreich (GFF) under the RTI-Dissertations 2024 (FTI24-D-030).

## Author Contributions

J.L. conceived the study.

M.M., L.R. and J.W. performed cryo-FIB, cryo-ET and cryo-ET data analysis.

M.M. performed subtomogram averaging and bioinformatic analysis of FtsZ AF2 models.

Y.Z. performed all SPA cryo-EM experiments for all EcFtsZs including all structure determination and modelling steps except for one dataset where M.M. collected and processed data.

D.K-C. expressed and purified all EcFtsZ proteins and mutants.

L.G. purified and performed all SPA cryo-EM experiments for TmFtsZ including all structure determination and modelling steps.

L.R. performed all *E. coli* genetics, plasmid construction, spot dilution assays and western blots.

J.W. performed *E. coli* imaging experiments for all EcFtsZ mutants.

G.C. expressed, purified and conducted all cryo-EM SPA experiments for BsFtsZ as well as pelleting assays and prepare the reactions for negative stain.

A.S. generated *B. subtilis* dCas9 BsFtsZ depletion strains and performed all spot dilution and imaging experiments of BsFtsZ mutants in these strains, as well as grid preparation and imaging for negative stain EM.

M.K. performed all coarse-grained MD simulations and analysis of datasets.

D.R. generated *B. subtilis* strains for live cell imaging, performed all live cell imaging and analysis and spot dilution assays of strains used in live cell imaging.

M.M. wrote the manuscript first draft. All authors edited the manuscript.

J.L., A.W., A.Š. and S.H. provided supervision and funding.

## Methods

### *E. coli* cryo-ET sample preparation and lamellae production by cryo-FIB milling

Preparation of samples of *E. coli* cells for subsequent tilt series collection varied slightly depending on whether samples were FtsZ_D212A_ overexpression or not. For FtsZ_D212A_ (GTPase mutant) overexpression, C41(DE3) cells, harbouring an arabinose-inducible FtsZ_D212A_ expression plasmid with ampicillin resistance (pBAD vector – see plasmid list in Table S1), were grown overnight in LB media supplemented with 100 μg/mL ampicillin at 37 °C. Overnight cultures were used to inoculate (1:100 dilution) M9 minimal media for a day culture where M9 minimal media consisted of 0.4% glycerol, 0.2% casamino acids, 2 mM MgSO_4_ and 100 μg mL^-1^ CaCl_2_. Day cultures were grown at 37 °C shaking at 200 rpm until optical density at 600 nm wavelength (OD_600_) 0.3 was reached. Expression was then induced with 0.1% arabinose until OD_600_ 0.6, after which cells were pelleted by centrifugation at 1800xg for 5 minutes. Cells were resuspended and concentrated 100-fold to OD_600_ ∼60.

For all other FtsZ overexpression sample preparations, overnight cultures and M9 media were prepared as described and expression vectors used the same arabinose promotor and resistance cassette. During day culturing in M9 media, cultures were grown to OD_600_ 0.6 at 37 °C before expression induction with 0.1% arabinose and were allowed to shake for another 2 hours before centrifugation and concentration to a similar OD_600_ ∼60.

When preparing dividing C41(DE3) cells (without an overexpression plasmid), LB media was used for the day cultures while no selection antibiotic was used in the overnight culture. Cultures were grown until mid-log phase (OD_600_ 0.3-0.6), growth was measured before centrifugation and cell resuspension at a target OD_600_ = 60.

For all preparations, concentrated cell suspensions were plunge frozen in liquid ethane using a Vitrobot Mark IV (Thermo Fisher Scientific) on Quantifoil 200 mesh 2/2 EM grids (either gold or copper support). Prior to plunge freezing, grids were glow discharged using a PELCO easiGlow for 45s at 25 mA. 8 μL of concentrated cell suspension was applied to the carbon face of glow-discharged grids before back blotting for 6 s, blot force 10 at room temperature with humidity control turned off. Back blotting was achieved using a custom Teflon ring in the shape of Vitrobot blot paper on the opposite grid face cells were applied to.

Plunge-frozen grids were clipped and loaded into an Aquilos 2 Dual Beam Cryo-FIB/SEM (Thermo Fisher Scientific) for lamellae production. Due to the extended timeframe over which sample preparations occurred, the exact milling methods evolved over time but remained broadly consistent. Typical milling sessions prepared grids for milling with an initial 40-60 s application of organoplatinum using the Gas Injection System (GIS), followed by 20-40 s of inorganic platinum sputter coating at 30 mA. A gallium source accelerated to 30 kV was used for all subsequent milling steps, typically at a 12° angle and automated using AutoTEM software by Thermo Fisher Scientific. This began with trench milling either side of the target milling area as described previously by Wolff *et al*^84^, followed by rough milling at 1 nA using rectangular milling boxes. Subsequent milling steps, each with a reduced current from the previous step (all using the cleaning cross-section settings) produced increasingly thinner and narrower lamellae until a final polishing step at 30 pA current resulted in a final width of 6 μm and a target thickness of ∼120 nm.

### Cryo-ET Data Collection

All tilt series collections of polished lamellae were carried out on a Krios G4 (Thermo Fisher Scientific) 300 kV Transmission Electron Microscope (TEM) equipped with a Falcon 4i detector, a Selectris X energy filter and a cold field emission gun. All datasets were collected using Tomo5 software from Thermo Fisher Scientific at a magnification of 83,000 and nominal pixel size of 1.514 Å/pixel (later calibrated to 1.504 Å/pixel). All datasets for subtomogram averaging (STA) FtsZ_D212A_ were collected using 3° tilt increments while other collections of dividing bacteria and overexpression of other FtsZ mutants were collected with 2° increments. See Table S2 for more details on STA data collection metrics and Table S3 for all other tilt series collections.

### EcFtsZ Subtomogram averaging (STA)

All processing was carried out using the Relion 5 STA pipeline^47^, across multiple datasets from multiple different collection dates. Tilt series were motion corrected with 5x5 patches using Relion’s implementation of MotionCorr2^85^ and contrast transfer function (CTF) estimation carried out using CTFFind-4.1^86^. Bad tilts were excluded using the Relion 5 Napari viewer and tilt series were aligned with the Relion 5 AreTomo2 wrapper^87^. Tomograms were reconstructed in Relion with an un-binned pixel size of 4000, 4000, 2000 (x, y, z) at 10 Å per pixel. Half tomograms were reconstructed for denoising, where all collected datasets except the final dataset were denoised in Relion 5 using CryoCARE^88^, while the final dataset was de-wedged and denoised using DeepDeWedge^89^. Filament paths were then traced manually using the Relion 5 Napari particle picker with the aim of tracing the centroid of each doublet. Particles were initially extracted from the filament traces every asymmetric unit (∼4 nm spacing) but due to problems with particle duplication during refinement, particles were extracted every 90 Å (roughly two asymmetric units). Filament polarity was kept unknown, allowing subsequent refinements to use bimodal psi searches.

Initially, a smaller subset of particles (mid-picking) was used to generate a *de novo* initial model with helical reconstruction mode selected but without helical symmetry applied^90^. This produced a low-resolution reconstruction of two tubes, enough to align particles to. Particles were extracted initially at 3x binning (4.542 Å/pixel) with a box size of 60 pixels (272.52 Å). The particles were refined using helical reconstruction, starting from some basic assumptions about the symmetry: from the tomograms and initial model, it was clear the doublets had effectively no twist while the rise of FtsZ is typically 44-45 Å^43,91,92^. Thus, helical symmetry parameters consisting of a 45 Å rise and 0° twist were assumed. No other symmetry was enforced initially. With a particle extraction every 90 Å, the number of asymmetric units was set to 2 and an outer cylinder of initially 110 Å was used. The refined map appeared to have 2-fold symmetry corresponding to Relion “D1” symmetry and concurrent refinements of *in vitro* FtsZ showed the same D1 symmetry. Hence all subsequent refinements applied D1 symmetry. Further refinements were conducted with symmetry searching and an outer helical diameter of 98 Å while CTF refinement and polishing steps were used to resolve secondary structure features. An 8 Å map showing secondary structure features was reconstructed with substantial overfitting becoming apparent. Particle duplication was a major problem throughout refinement and inspection of some refined particles in ArtiaX^48^ revealed many picked coordinates failed to align to the nearest filament. This resulted in a re-annotation of filament paths to minimise particle misalignment resulting in fewer but better starting coordinates. Concurrently, a recently developed tilt series alignment tool called Missalignment^93^ was used on the dataset to further refine tilts series and apply a 2D deformation model to tomograms in Relion 5^47^. Newly picked coordinates were refined with D1 symmetry from the beginning and subsequent 3D refinement and CTF refinement steps returned a reconstruction at 6.9 Å resolution, when accounting for pixel size re-calibration. The final map with a twist of -0.116° and rise of 44.37 Å (pixel-calibrated to 1.504 Å/pixel from 1.514 Å/pixel) was sharpened (model-free) using LocScale 2.0 feature enhancement^94^ and deposited in the EMDB under accession EMD-100280. Refinement steps are summarised in brief in Figure S2.

For WT FtsZ STA in dividing *E. coli* C41(DE3) cells, Relion 5 STA pipeline was also used with a slight deviation in preprocessing steps. Tilt series were motion corrected and CTF estimated as above but alignments were imported from AreTomo3^95^, where tilt series were aligned, reconstructed in AreTomo3 and denoised/de-wedged with Isonet2^96^, before processing further in Relion 5. Filaments were picked as described above except single protofilaments were annotated individually over doublets. This was to avoid biasing results and accounting for potential difficulty in seeing the second protofilament of a doublet. Particles were extracted in bin 3 as above and refined against either a cylinder in C1 or a doublet reference with both C1 and D1 symmetry applied. All refinements assumed basic helical symmetry of 0° twist and 45 Å rise. See Figure S1.

### Cryo-ET data processing for FtsZ visualisation

For all tilt series collected that were not used as part of the FtsZ_D212A_ subtomogram averaging, tilt series were motion corrected, CTF estimated aligned and reconstructed entirely using AreTomo3^95^. Reconstructed half-volumes were then used to denoise and de-wedge using Isonet2^96^. Segmentations were used to highlight features, particularly due to the small size of FtsZ protofilaments. Membranes and ribosomes were segmented automatically using Easymode^46^ while Napari tomoslice^97^ was used to manually annotate and fit paths for FtsZ protofilaments. These were converted into spline segmentations in MRC format using an in-house script in order to visualise alongside other segmented membrane and ribosome volumes. Tomograms were typically visualised using IMOD^98^, while segmented volumes were visualised using ChimeraX^99^.

### EcFtsZ expression, purification and polymerisation

*E. coli* FtsZ constructs were commercially synthesised and cloned into either the pET-24a(+) or pET-24b(+) vectors, both containing an N-terminal 10×His-SUMO tag, with the SUMO sequence codon-optimised for expression in *E. coli*. The resulting plasmids (Table S1) were supplied by GenScript and verified by whole-plasmid sequencing upon arrival.

FtsZ WT, Δ2–6 and R174D were expressed in *E. coli* OverExpress™ C41(DE3) cells (LGC, Biosearch Technologies), whereas FtsZ Δ2–10 was expressed in *E. coli* BL21(DE3) cells (MilliporeSigma). Cells were grown in 2×TY medium (16 g L⁻¹ tryptone, 10 g L⁻¹ yeast extract and 5 g L⁻¹ NaCl; pH 7.4) supplemented with kanamycin (50 µg mL⁻¹) and 1% glucose. Cultures were incubated at 37°C with shaking at 200 rpm until an OD₆₀₀ of 0.8 was reached. The temperature was then reduced to 30°C for FtsZ WT, Δ2–6 and R174D, and to 25 °C for FtsZ Δ2–10. Protein expression was induced by adding 0.5 mM isopropyl β-D-1-thiogalactopyranoside (IPTG), followed by overnight incubation.

Cells were harvested by centrifugation and resuspended in lysis/purification buffer (50 mM Tris-HCl, 250 mM NaCl, 50 mM KCl and 3% (v/v) glycerol; pH 8.0) supplemented with cOmplete™ ULTRA Mini, EDTA-free EASYpack Protease Inhibitor Cocktail tablets (Roche, Merck). Ten tablets were added per cell pellet obtained from 1 L of bacterial culture to minimise proteolysis of the FtsZ C-terminus. All purification steps were performed at 4 °C.

Cell lysis was performed using a manual cell grinder, with liquid nitrogen used to pulverise frozen *E. coli* cells while minimising sample heating and protein degradation. The pulverised cells were added to 150–200 mL of lysis/purification buffer, followed by sonication. The lysate was subsequently clarified by centrifugation. The resulting supernatant was incubated with 5 mL of HisPur™ Cobalt Resin (Thermo Scientific, cat. no. 89965) for 45 min with gentle agitation using a batch-binding approach. After the incubation the resin was subsequently transferred to a Bio-Rad Glass Econo-Column (2.5 × 10 cm; cat. no. 7372512) and extensively washed with lysis/purification buffer under gravity-flow conditions.

The bound proteins were released directly from the resin by on-column cleavage of the SUMO tag using His-tagged SENP (SUMO-specific protease). The plasmid encoding SENP protease was a gift from Dmitry G. Vassylyev, and the protease was purified as previously described by Vassylyeva et al^100^. SUMO tag cleavage was performed in 1 mL of cleavage buffer containing 50 mM Tris-HCl, 250 mM NaCl, 50 mM KCl, 3% (v/v) glycerol and 0.5 mM TCEP (pH 8.0), with gentle agitation for 10 min. The purified proteins were confirmed to have the expected molecular masses by intact-mass spectrometry. Samples were also analysed by SDS–PAGE using a 12% Criterion TGX (Tris-Glycine eXtended) Precast Midi Gel (Bio-Rad). The resulting eluates were used immediately for cryo-EM sample preparation without prior concentration.

The eluates were diluted with lysis/purification buffer to estimated final protein concentrations of 0.1, 0.25, 0.2 and 0.9 mg mL⁻¹ for FtsZ WT, R174D, Δ2–6 and Δ2–10, respectively. The diluted protein solutions were immediately subjected to polymerisation without prior freezing. Polymerisation was initiated by adding MgCl₂ to a final concentration of 5 mM and GMPCPP (guanosine-5′-[(β,γ)-methylene]triphosphate) to a final concentration of 0.2 mM, followed by incubation on ice for 1 h.

For single-particle analysis (SPA) by cryo-EM, grids were prepared either using the sandwich Langmuir–Blodgett film method (manuscript submitted^49^) or using a Vitrobot Mk IV (Thermo Fisher Scientific). For Vitrobot-assisted grid preparation, the chamber was maintained at 100% relative humidity and 4 °C, with a blotting time of 3 s and a blot force of 3. Quantifoil copper grids (300 mesh, 1.2/1.3) were glow-discharged at 25 mA for 60 s. A sample volume of 3 µL was applied to each grid, with no waiting time before blotting.

### TmFtsZ expression, purification and polymerisation

FtsZ from *T. maritima* was purified as previously described by Szwedziak *et. al.* with some alterations^12^. In brief, plasmid pSZ7 (Table S1) was transformed into *E. coli* C41(DE3) competent cells with overnight cultures grown in Terrific Broth (TB) medium at 37 °C with shaking. 12 L day cultures were inoculated with overnight cultures and grown in TB medium at 37 °C, shaking at 200 rpm. Protein expression was induced at OD_600_ 0.7 with 1 mM IPTG, followed by a temperature reduction to 20 °C and growth for a further 16 h before centrifugation to collect cell pellets.

All buffers were identical to those previously reported by Szwedziak *et. al*^12^. Cell pellets were resuspended in buffer A (50 mM Tris, 10 mM EDTA, 2 mM 1,10-phenantroline, 1 mM PMSF, pH 8.5) supplemented with cOmplete protease inhibitor tablets and lysed by one passage through a Constant Systems cell disruptor at 25 kpsi. The lysate was de-foamed by a short centrifugation followed by a 30 min ultracentrifugation at 27,000 rpm using a Beckman 45 Ti rotor with temperature set to 4 °C. The supernatant was loaded onto a 100 mL chitin resin column, washed with 150 mL of buffer B (A + 50 mM NaCl), 150 mL of buffer C (B + 50 mM DTT) and eluted in 150 mL buffer B. The eluent was then loaded onto a 5 mL Q XL column (Cytiva) and eluted twice before washing with 100 mL of buffer D (20 mM Tris, 10 mM EDTA, 2 mM 1,10-phenantroline, 1 mM PMSF, pH 7.5) mixed with 5% buffer E (D + 1 M NaCl) and then eluting in buffer E. Peak fractions were identified by SDS-PAGE and concentrated in Amicon Ultra Centrifugal Filter regenerated cellulose MWCO concentrators followed by gel filtration using an Cytiva Sephacryl S-200 10/300 column. Peak fractions were again concentrated to ∼10 mg mL^-1^ and stored at -70 °C for later use.

To prepare filaments for cryo-EM, aliquots were thawed and diluted 1:10 in buffer TEN_7.5_ (50 mM Tris, 100 mM NaCl, 1 mM EDTA, pH 7.5) and supplemented with 1 mM GMPCPP and 5 mM MgCl_2_. Immediately before grid preparation, the sample was further diluted in buffer TEN_7.5_ supplemented with 0.5 mM GMPCPP and 5 mM MgCl_2_ to ∼0.2 mg mL^-1^. Quantifoil 300 mesh R 1.2/1.3 grids were glow discharged and manually back-blotted following application of 3 μL of sample.

### BsFtsZ expression, purification and polymerisation

The BsFtsZ sequence was amplified by PCR from *B. subtilis* (strain 168) (Table S6) and cloned into a pT7 vector containing an N-terminal 6×His-SUMO tag using Gibson assembly (Table S1). FtsZ mutants were constructed using pairs of complementary primers carrying the desired deletion or point mutations. All resulting plasmids were verified by Sanger sequencing. BsFtsZ constructs were expressed in *E. coli* BL21(DE3) using an autoinduction method^101^. After an initial incubation of 4 h at 37°C, the cells were grown for 20 h at 20°C in 2xTY medium, complemented with autoinduction supplement and 50 μg/mL kanamycin. Cells were harvested, flash frozen in liquid nitrogen, and stored at −20°C. Cell pellets were resuspended in lysis buffer (50 mM HEPES, 300 mM KCl, 10 mM imidazole, 5% glycerol, 1 mM MgCl_2_, pH 8.0, benzonase, lysozyme, 0.25 mM Tris (2-carboxyethyl) phosphine hydrochloride (TCEP), EDTA-free protease inhibitor cocktails (Roche)) at 4°C and lysed by sonication. Cell debris were removed by centrifugation (15,000 × g) for 15 min at 4°C, and the supernatant loaded onto a Ni-NTA affinity chromatography column (HisTrap FF, Cytiva) pre-equilibrated in buffer A (50 mM HEPES, 300 mM KCl, 10 mM imidazole, 5 % glycerol, pH 8.0). His-tagged proteins were eluted with a linear gradient of buffer B (50 mM HEPES, 300 mM KCl, and 0.5 M imidazole, pH 8.0). The fractions of interest were pooled and dialysed in the presence of SUMO protease at a 1:100 wt/wt ratio. Dialysis was carried out at 4°C overnight in SEC buffer (25 mM HEPES, 150 mM KCl. pH 8.0). Cleaved His-tags and His-tagged SUMO protease were removed with Ni-NTA agarose resin. The cleaved protein was concentrated and injected onto a Superdex 200 16/60 size exclusion column (GE Healthcare) pre-equilibrated at 4°C in SEC buffer. The peak corresponding to the protein was concentrated, flash frozen in small aliquots in liquid nitrogen, and stored at −80°C.

Frozen aliquots of BsFtsZ were thawed, pre-cleaned by centrifugation at 16,000xg for 30 min to remove any large aggregate, and diluted in polymerisation buffer (100 mM KCl, 10 mM MgCl_2_, 25 mM Pipes pH 6.9) to a final concentration of 10 µM (∼ 0.5 mg/mL). To initiate polymerisation, GTP was added to final concentration of 2 mM. After 5 min incubation at room temperature, 5 µL of sample were applied onto a Quantifoil grid (Cu R2/2 300-mesh), glow discharged at 15 mA for 25 seconds with a Pelco easiGlow device, and plunge frozen in liquid ethane using a Vitrobot Mark VI operating at 18°C and 100% relative humidity. Grids were blotted for 4s and blot force set to 0.

### EcFtsZ single particle cryo-EM data collection, processing and atomic model building

Single-particle datasets were collected on a Titan Krios (Thermo Fisher Scientific) operating at 300 kV, equipped with either a Falcon 4i direct electron detector and a Selectris X energy filter set to a slit width of 10 eV or a Gatan K3 direct electron detector and a Gatan Quantum energy filter set to a slit width of 20 eV, and controlled by EPU. Per-dataset acquisition parameters (nominal magnification, pixel size, defocus range, dose and frame number) are summarised in Table S4.

Movies were motion-corrected in cryoSPARC with patch motion, and CTF parameters were estimated with patch CTF. Particles were picked with blob picker, extracted in a 384-pixel box binned three-fold to 128 pixels for initial alignment (FtsZ_Δ2-10_ was extracted in a 512-pixel box binned four-fold to 128 pixels), and an initial model was generated *de novo* from selected 2D classes without imposing symmetry. Particles were cleaned by 2D and 3D classifications. For each structure, the initial map was refined by helical refinement in cryoSPARC without imposing helical symmetry (C1, no helical parameters specified). Once the resolution approached the Nyquist limit of the binned pixel size, typically after this first unconstrained round, particles were re-extracted at full (unbinned) box size (384 pixels for FtsZ and FtsZ_Δ2-6_; 512 pixels for FtsZ_Δ2-10_). Helical twist and rise were then determined by searching a grid of parameters (twist −90° to 90°; rise 40–50 Å) and refining iteratively in cryoSPARC until the parameters converged (FtsZ: twist −1.1°, rise 45.36 Å; FtsZ d2-6: twist 1.09°, rise 45.71 Å; FtsZ d2-10: twist 89.67°, rise 11.35 Å). The resulting maps were refined with and without D1 point-group symmetry imposed and compared to confirm the correct point group, which was D1 for all three structures. The final particle sets were then refined with iterative per-particle CTF refinement (beam tilt, anisotropic magnification, higher-order aberrations, per-particle defocus and per-micrograph astigmatism) and Bayesian polishing^85,102^. All datasets were processed in cryoSPARC 4.7.0 using helical refinement (Punjani et al., 2017)^103^. Final resolutions were estimated from the Fourier shell correlation between independently refined half-maps at the 0.143 criterion^104^. Initial atomic models were generated by rigid-body fitting of PDB 8IBN into each cryo-EM map, followed by iterative rounds of manual model building in Coot^105^ and real-space refinement in Phenix^106^, until model geometry and fit to the density converged.

### *E. coli* plasmid construction and spot dilution assays

Plasmids were constructed using Gibson assembly and electroporated into electro competent E. cloni® cells (Lucigen). Single colonies were then mini-prepped and whole plasmid sequenced before use in further experiments. Mutations were made by replacing codons with the highest usage codon from a standard E. coli codon usage table. See Table S1 for all plasmids used, Table S5 for resulting strains and Table S7 for oligonucleotides.

For spot dilution assays and imaging, *E. coli* JKD7-1^51^ (strain harbouring an ectopic copy of *ftsz* on a temperature sensitive plasmid with chloramphenicol resistance) was transformed with the appropriate plasmid and grown on LB agar plates overnight with 100 μg/mL ampicillin and 1% glucose for catabolite repression of the pBAD plasmids containing *ftsZ* at 30°C. Chloramphenicol was avoided as it was found to exacerbate phenotypes. Unless indicated otherwise, the ampicillin glucose medium is the standard medium used for these experiments and growth of JKD7-1 was always done at 30°C to avoid loss of the temperature sensitive pKD3 plasmid harbouring the WT *ftsz* copy. The next day, single colonies were used to prepare cultures for glycerol stocks and overnight cultures for the next day. On the day of the experiment, the overnight cultures were diluted 1:100 into LB medium with 1% glucose and grown for 2 h. Next, the cultures were diluted 1:5 serially into 80 μL (total 100 μL) of LB medium distributed across 96 well plates 6 times. The dilution series of cells was then spotted onto LB agar plates containing indicated supplements with a stamp and grown at 30°C or 42°C. Cells grown at 42°C were imaged on the morning of the next day and cells grown at 30°C in the evening. Images were taken on a Bio-Rad Gel Doc XR+ Gel Documentation System by exposing the agar pads to white light for 0.5 s. Images were then contrast adjusted and inverted for visualisation. To confirm that the strains contain the correct plasmids, day cultures from which the spot tests were done were continued to grow over night and the plasmids in these strains verified by whole-plasmid sequencing.

### *E. coli* light microscopy, western blotting and SDS PAGE

Cultures of all temperature sensitive *E. coli* strains harbouring various FtsZ mutant plasmids (Table S1, S5) were grown overnight in LB supplemented with 100 µg/mL ampicillin at 30°C with shaking at 210 rpm. All subsequent cultures were also grown with shaking at 210 rpm. Overnight cultures were used to inoculate (1:100 dilution) 5 mL LB cultures with no antibiotic that were grown for 2 h at 30°C. Cultures were then back inoculated 1:20 in fresh LB and grown for 4 h at 42°C to deplete the plasmid containing WT *ftsz*. Finally, cultures were back inoculated again 1:20 and grown at 37°C for 2 h +/-0.1% arabinose, to test the phenotype of the FtsZ mutants. 2 µL of culture was applied to a 2% agarose LB pad to immobilise the cells and the spot was left to dry for 1 minute before a 22x22 mm coverslip was applied and the cells were imaged. Samples were imaged on a Zeiss LSM900 microscope with a 1.4 numerical-aperture (NA) Plan Apochromat x63 oil objective. Contrast and brightness of images was adjusted to improve clarity.

To verify cellular FtsZ levels by western blot (Figure S4), cells were grown as mentioned above and at chosen time points, 1 mL of cell culture were harvested by centrifugation at 21,000 xg for one minute, before resuspension in 200 μL of 1x sample buffer to OD_600_ 0.5. 100 µl of these samples was then heated for 5 min at 99°C and cooled down for 1 min in a metal block. Then, 1 µL of Benzonase® Nuclease (Sigma) was added and samples were incubated for another 5 min at 37°C. Subsequently, 5 µL of the samples was loaded on a SDS gradient gel (Criterion TGX Precast gels, 4-20%, Bio-Rad). The gel was then run for 10 min at 200 V and 400 mA to get sharp bands. Next, the protein from the gel was transferred onto a PVDF membrane using Trans-Blot Turbo Transfer Pack (Bio-Rad) and blotting performed in a Trans-Blot Turbo Transfer System (Bio-Rad) with the manufacturer preset mixed molecular weights transfer. The PVDF membrane was then blocked with 50 mL of 5% skimmed milk in PBS 0.05% Tween 20 (PBST) for 30 mins. Afterwards, 20 mL of the PBST milk solution were discarded, and FtsZ primary antibody was added to the PBST milk at 1:5.000 (0.2 ng/mL) and left to incubate for 30 mins. Following the incubation, the PBS milk solution was exchanged with 50 mL PBST thrice and incubated for 5 mins during the last wash step. After the last wash step, the PBST was exchanged with 30 mL of fresh PBST and anti-rabbit HRP secondary antibody was added at 1:10.000 (0.1 ng/mL) and incubated for 30 mins. The secondary antibody was then removed by washing thrice with 50 mL of PBST for 5 mins. Detection of the secondary antibody was achieved using 2 mL of the Amersham™ ECL Detection Reagents and a Bio-Rad Gel Doc XR+ Gel Documentation System and averaging the signal over 0.5 s. Lastly, an SDS-PAGE gel was run using 10 µL of all samples to verify that equal amounts of protein were loaded and transferred to the membrane.

### TmFtsZ single particle cryo-EM data collection, processing and model building

TmFtsZ SPA datasets were collected on a Titan Krios G4 (Thermo Fisher Scientific) 300 kV TEM microscope equipped with a Falcon 4i detector and Selectris X energy filter using the software EPU. Dataset parameters are listed in more detail in Table S4. Initial processing attempts revealed strong particle orientation preference, so additional micrographs were collected with alternating stage tilts between 30°, 45° and 60°, to improve the particle orientation sampling.

All processing was carried out using cryoSPARC^103^. Micrographs were patch motion-corrected and patch CTF-estimated before automated filament picking and 2D classification. Particles were initially classified in 2D to remove junk and classes showing single filaments. An initial model was made *ab initio* before a helical refinement without symmetry parameters imposed. Helical parameters were then estimated using the symmetry search tool and a successive helical refinement using these parameters and D1 symmetry was run. Helical parameters and alignments were improved iteratively through successive refinements while orientation preferences were corrected for between each refinement using the particle rebalance tool. Global and local CTF-refinement jobs were run before further particle cleaning with 2D classification and 3D classification in RELION^102^. A final reconstruction at 3.7 Å global resolution was sharpened with EMReady^107^. Despite the tilted data collections, the final reconstruction still suffered from anisotropy. An initial model was made from the EcFtsZ_WT_ filament model by using SWISSMODEL^108^ to thread the TmFtsZ sequence through the EcFtsZ model. The model was docked into the map in ChimeraX^99^, manually assessed for correctness before real space refinement using Phenix^106^.

### BsFtsZ single particle cryo-EM data collection, processing and model building

All data acquisition parameters are recorded in Table S4. BsFtsZ movies were collected using EPU software (Thermo Fisher Scientific) at the Institut Pasteur NCF core facility and processed within the CryoSPARC 4 environment. Motion correction and CTF estimation were performed patch-wise with default settings. Micrographs were filtered for further processing based on several quality requirements (CTF fit resolution better than 5 Å, full-frame motion distance below 30-pixel, relative ice thickness between the 1.0 and 1.3, astigmatism lower than 900 Å). FtsZ segments were picked using the Filament Tracer with the diameter set to 50 Å and using 2D-templates encompassing both single and double filaments classes. The traced filaments were filtered based on NCC and power scores to remove picks on the carbon support. Selected particles were extracted with a box size of 416 pixels and 2x binning, and subjected to several rounds of 2D classification to sort single and double filaments. The datasets collected at 0° (4697 movies) and 30° (5418 movies) stage tilt were processed separately up to this point, and particles were merged after the 2D classification cleaning. Selected particles were submitted to multi-class *ab initio* reconstruction (5 classes) with the following parameters, starting resolution set at 10 Å, maximum resolution set at 6 Å, initial and final minibatch size set at 300 and 1000, respectively. Similar ab initio models and the corresponding particles were combined and subjected to heterogeneous refinement, including decoy volumes corresponding to single filaments and junk classes. Segments assigned to classes with clearly resolved doublet features were taken forward to an initial helical refinement in C1, using starting helical parameters of 45 Å rise and 0°. After one round of 3D classification followed by helical refinement, Local and Global CTF refine, the volume was aligned to the symmetry axis, and D1 symmetry was enforced. The last helical refinement job produced a map with estimated resolution of 4.1 Å (FSC = 0.143), in which domain boundaries and secondary-structure elements were unambiguously resolved. A starting model for BsFtsZ was generated with AF2 in single-sequence mode, supplying the T-state EcFtsZ as a structural template to bias the prediction towards the tense conformation. The model was placed into the cryo-EM density by rigid-body fitting in ChimeraX^99^ and relaxed by interactive molecular dynamics flexible fitting in ISOLDE^109^. The starting model was then refined against the half-maps, applying jelly body and NCS symmetry restraints in Refmac Servalcat^110^. Model geometry was assessed with MolProbity^111^ and the validation statistics are reported in Table S4. Model vs. map FSCs were calculated with *phenix.mtriage*^106^.

### FtsZ knockdown in *B. subtilis* and complementation

WT *ftsZ* knockdown in *B. subtilis* was performed using a previously described CRISPR interference (CRISPRi) system based on dCas9^54^. Plasmids pJMP1 and pJMP3 were obtained from Addgene (accession nos. 79873 and 79875, respectively). A single-guide RNA (sgRNA) targeting the *ftsZ* open reading frame was designed using CRISPR@Pasteur (https://crispr-browser.pasteur.cloud)^112^. The sgRNA sequence 5′-TTCGCGATTTGTGCGATAAC-3′, which displayed a high overall design score, was selected. The sgRNA sequence was introduced into pJMP3 by PCR using oligonucleotides F_ftsZ_gRNA and R_ftsZ_gRNA (Table S7). The resulting pJMP3-ftsZ construct was integrated into the *thrC* locus of *B. subtilis 168* by natural transformation. Transformants were selected on spectinomycin-containing plates, and correct integration at the *thrC* locus was verified by colony PCR using oligonucleotides F_thrC_seq_genome and R_thrC_seq_genome (Table S7). The resulting strain (Table S6) was rendered competent and transformed with pJMP1 (Table S1), carrying dCas9 under control of the xylose-inducible promoter (Pxyl). Transformants were selected on spectinomycin and erythromycin/lincomycin-containing plates, and correct integration at the *ganA* locus was confirmed by colony PCR using oligonucleotides R_ganA_seq_genome and F_ganA_seq_genome (Table S7). Knockdown of native *ftsZ* expression was induced by supplementation of the growth medium with 1% (w/v) xylose.

For complementation experiments, an ectopic copy of *ftsZ* under the control of the same Pxyl promoter was integrated at the *amyE* locus. This ectopic *ftsZ* copy contained four synonymous substitutions within the sgRNA-target sequence, using codons of similar frequency, to render the strain “immune” to CRISPRi targeting. The complementation construct was assembled by Gibson assembly in the pSweet vector (Addgene plasmid no. 210272), the vector was linearised via PCR using oligonucleotides F_pSweet and R_pSweet and the *ftsZ* fragment was amplified using F_FtsZ and R_FtsZ (Table S7). Mutation of the sgRNA target was done via PCR with oligonucleotides F_FtsZ_mut and R_FtsZ_mut (Table S7). Plasmids were sequence verified and transformed into the *ftsZ* knockdown strain. Transformants were selected on chloramphenicol-containing plates, and correct integration at the *amyE* locus was verified by colony PCR using oligonucleotides F_AmyE_front and R_AmyE_back (Table S7). Complementation constructs encoded either WT FtsZ, or FtsZ variants carrying mutations at the interface of interest (Table S6). In all complementation experiments, 1% xylose was used to simultaneously reduce expression of the native *ftsZ* gene through CRISPRi and at the same time induce expression of the ectopic complementation construct. Mutants were constructed with the following oligonucleotides: Δ1-11 (F_FtsZ_ΔN1 and R_pSweet), K175E (F_FtsZ_K175E and R_FtsZ_K175E), E87K+E90K (F_FtsZ_E87K+E90K and R_FtsZ_E87K+E90K).

Western blotting with a commercial anti-FtsZ antibody (ABS2200, Sigma-Aldrich) was used to confirm that FtsZ levels following induction with 1% xylose were comparable to native FtsZ levels. All knockdown experiments were performed in LB medium at 37°C. Antibiotics were used at the following final concentrations: for *B. subtilis*, erythromycin, 1 µg/mL; lincomycin, 12.5 µg/mL; spectinomycin, 100 µg/mL; chloramphenicol, 6 µg/mL; and kanamycin, 7.5 µg/mL.

### BsFtsZ pelleting assay

Frozen aliquots of BsFtsZ were thawed, pre-cleaned by centrifugation at 16,000xg for 30 min to remove any large aggregate, and diluted in polymerisation buffer (100 mM KCl, 10 mM MgCl_2_, 25 mM Pipes pH 6.9) to a final concentration of 10 µM (∼ 0.5 mg/mL) in a final volume of 50 uL. Polymerisation was initiated by addition of GTP (2.5 mM) and the samples were pelleted by ultracentrifugation at 186,000xg for 1 h in a TLA 55 rotor at 4 °C in an Optima MAX-XP ultracentrifuge (Beckman). After supernatant removal, the pellet was resuspended in 50 uL of buffer and both fractions were mixed with equal volumes of Laemmli buffer and loaded onto a 12% SDS-PAGE gel. Bands were visualised by Coomassie staining and quantified using densitometric analysis within ImageLab software (Bio-Rad). The percentage of protein in the pellet fraction was calculated by dividing the intensity of the pellet fraction by the sum of the intensities of the pellet and supernatant fractions. Each condition was repeated at least in triplicate.

### Negative stain transmission electron microscopy (TEM)

FtsZ polymerisation was performed as described above. Carbon coated 400-mesh copper grids (CF 400-Cu; Electron Microscopy Sciences) were glow-discharged using an ELMO system for 30 s at 2 mA. A 5 µL aliquot of each reaction was applied to the grid and incubated for 1 min. Excess sample was removed by blotting, and the grids were washed three times with water, followed by two washes with 2 % aqueous uranyl acetate, before being air-dried. TEM images were acquired using a JEM-120i microscope equipped with an HC polepiece and a bottom-mounted CMOS SightSKY camera (JEOL, Japan) at the Ultrastructural BioImaging (UBI) Core Facility, Institut Pasteur.

### BsFtsZ treadmilling imaging and analysis by VerCINI

All *B. subtilis* strains used in VerCINI imaging experiments integrated an IPTG inducible copy of *ftsAZ*, containing wild type or mutant *ftsZs* (see Table S6), at the non-essential *amyE* locus, along with *ftsZ-gfp* at the native locus. This was achieved through inducing competency using standard methods described previously^113^. Double recombination at *amyE* was confirmed by testing alpha-amylase activity on starch plates. All point mutants and *ftsZ-gfp* fusion constructs were confirmed by DNA sequencing of the *amyE* and *ftsAZ* loci. To prepare slides for imaging, 650 µL of molten 6% ultrapure agarose (Invitrogen) dissolved in S750^Glucose^ media as previously described^114^ was pipetted over a silicone micropillar array prior to the addition of a slide containing a GeneFrame (Thermo), which cast the pillars^58^. Once set, excess agarose around the micropillars was removed to ensure sufficient oxygenation of cells during imaging. Overnight cultures were generated by inoculating colonies into LB media (Oxoid) supplemented with 0.4% glucose (Sigma), which were grown at 30°C with orbital agitation at 175 rpm. The next day, overnight cultures were washed to remove traces of LB, prior to being diluted to an optical density at 600 nm (OD_600_) of 0.0125-0.05 in freshly prepared S750^Glucose^ media. Liquid day cultures were grown in flasks at 30°C with 175 rpm shaking until OD_600_ reached 0.4 +/-0.05. Cultures were then back-diluted to OD_600_ = 0.25 in S750^Glucose^ supplemented with 100 µM IPTG and grown for a further 50 min to induce expression of the *ftsZ-*wild type or filament breaker/enhancer mutants. Induced cells were concentrated 40x and 8 µL were spotted over the microholes cast in the agarose pad, then slides were centrifuged for 5 min at 2204xg (Eppendorf Centrifuge 5430). Pads were washed to remove cells that did not load into the microholes prior to the addition of a coverslip (Thorlabs, 22 x 22 mm^2^, thickness no. 1.5).

Imaging was conducted on a custom-built microscope where cells were illuminated with a 488 nm laser (Vortran Stradus). A 100x TIRF objective was used (Nikon CFI Apochromat TIRF 100XC Oil), as well as a 200 mm tube lens (Thorlabs TTL200) and a Prime BSI sCMOS camera (Teledyne Photometrics) generating an apparent pixel size of 65 nm/pixel. Using a pair of galvanometer mirrors (Thorlabs) spinning at 200 Hz, image acquisition was conducted in ring-HiLO^115^, which enabled high signal-to-noise imaging with uniform illumination. FtsZ-dynamics were imaged at 1 frame/s continuous exposure for 2 min at a power density of 1.2 W/cm^2^. After each timelapse, a Z-stack was acquired (+/-300 nm from the imaging plane in 100 nm steps) to enable the exclusion of out-of-focus Z-rings and/or tilted cells during image analysis. The microscope was controlled using Micro-Manager (v 2.0 gamma).

Detailed protocols on image analysis have been published previously^31,58^ but in brief, FtsZ rings were marked and generated ROI’s saved using the verciniAnalysisJ plugin. Images were then denoised using the PureDenoise plugin, based on wavelet decomposition. Denoised videos were registered using StackReg, prior to cropping and saving of individual rings. Ring fitting was done on a frame-by-frame bases, generating fitted kymographs, using custom MATLAB scripts as described previously^31,58^. FtsZ-filament dynamics were manually annotated on the generated kymographs in ImageJ by drawing lines on FtsZ tracks. To aid in annotating FtsZ-filaments, particularly those that were dim, a ridge filter was applied to the kymograph using the verciniAnalysisJ plugin (set as 2 pixels, 130 nm). Using the length and angle of the marked kymograph traces, filament speeds were calculated. The raw data were plotted as violin plots, and manual interrogation revealed two populations: immobile and mobile filaments (Figure S19A). Based on this, filament speeds < 10 nm/s were classified as immobile, and filament speeds >10 nm/s were classed as mobile. The mobile filament speeds were plotted, and pairwise comparisons of the various strains were interrogated using a Mann Whitney U test. See Table S8 for P values. All experiments were conducted in biological duplicate. The n numbers of mobile and immobile filaments analysed in each repeat are shown in Table S9.

TIRF movies of each strain were acquired immediately prior to VerCINI imaging of cells lying on the top of the pad. Images were acquired 1 frame/sec continuous exposure for 2 mins at a power density of 1.2 W/cm^2^. Cropped images were upscaled 4x in x and y without interpolation. Movies are shown at 20 fps.

Spot dilutions of all strains used in the VerCINI imaging were done to assess the viability of each strain. Each were grown overnight in LB + 0.4 % glucose at 30°C with orbital agitation at 175 rpm. Day cultures were made by diluting overnights to OD_600_ = 0.05 in fresh LB, which were grown at 30°C until OD_600_ = 1. Each culture was then OD normalised to 0.5, prior to a series of 1:10 dilutions being made (down to 10^-6^). 3 µL of each dilution was spotted onto NA plates in the presence or absence of 100 µM IPTG. Plates were incubated at 30°C for 24 h prior to imaging.

### FtsZ AlphaFold analysis

To probe FtsZ doublet interface conservation, AlphaFold2 (AF2)^52^ models of monomeric FtsZ were aligned to either side of the doublet interface (a dimeric coordinate file was created with only one subunit per protofilament) and proximal residues annotated. To do this, the FtsZ sequence (EcFtsZ or BsFtsZ) was BLAST^116^ searched using the UniProt webserver^117^ against the UniProtKB database selecting for either Pseudomonadota [1224] (searched with EcFtsZ sequence) or Bacillota [1239] (searched with BsFtsZ sequence) phyla. UniProt ID numbers were downloaded and to overcome the server limits on search size and to search as wide as possible within the phylum, the least similar sequence in the search was used to re-search the same database. This was repeated once more to produce a list of UniProt IDs which were used to download pre-folded AF2 predictions (duplication events were avoided where identical UniProt ID models were overwritten) from the AF Database (AFDB)^118^. Of the total UniProt IDs with an AF2 prediction (770 Pseudomonadota and 1,414 Bacillota), a substantial proportion of sequences were identical or highly similar to at least one other UniProt ID. Here the dataset was split to minimise amplification of duplicated signal, where the models were filtered based on 90% sequence identity. A custom script was written which took the reference doublet dimer (labelled chain ‘A’ and ‘B’) and the AF2 predictions as input, removed the final 50 residues of the CTT (to prevent misfolded IDR from interfering with results). Models were then duplicated and structurally aligned to either chain A or chain B using US-align^119^, before being subject to *in vacuo* energy minimisation with OpenMM^119^ using the Amber14 forcefield^120^. Minimised proteins now had hydrogens so when measuring contacting residues, a short 3 Å cutoff was applied where residues from one chain within this distance from the other chain were counted as contacting. Histograms of per residue contact counts were plotted for chain A from which short regions with high contact counts selected to represent interfaces via a weblogo^121^. Full contact histograms pre and post screening for sequence identity can be found in Figure S8.

### Coarse-grained MD simulations and modelling

FtsZ filaments were modelled as coarse-grained polymers where each monomer consisted of three beads: a central backbone bead (∼5 nm as used previously^32^) and two lateral patch beads positioned at 90° to one another. The patches represented the C-terminal (C-C) and N-terminal (N-N) lateral interfaces and were placed at 0.5 σ and 0.9 σ from the central bead, respectively. Both interfaces were modelled as short-range attractions using the cosine/squared potential in LAMMPS^122^ (r_min = 0.1 σ, r_cut = 0.4 σ) with strengths of ε = 10 k_B_T for the C-C interface and ε = 5 k_B_T for the N-N interface. The Weeks-Chandler-Anderson (WCA) component of this potential was omitted to avoid numerical instabilities and all remaining interactions were purely volume exclusion. Monomer spacing and patch positions were maintained using harmonic bonds (k = 50,000 k_B_T) and the 90° patch geometry was enforced with a stiff angular potential (k = 50,000 k_B_T). Filament bending rigidity was set by an angular potential (κ = 1000 k_B_T), chosen to qualitatively reproduce zipping up of two filaments upon lateral contact. In contrast to previous work^32^, no aligning force towards the cell circumference was applied.

Treadmilling was implemented as published previously^32^. Wild-type rates were r_on_ = 16 s⁻¹ and r_off_ = 10 s⁻¹, with a hydrolysis time of τ_hydrolysis_ = 0.1 s, chosen by fitting simulated monomer lifetimes to *in vivo* single-molecule data from *B. subtilis* (Figure S18)^39^. Kinetic modulation was implemented by lowering r_off_ to 8 s⁻¹ when a C-C interface was formed on the terminal monomer of a filament. The treadmilling profile was as published^32^, but its width was increased from d = 10 to d = 20 to account for the wider filaments resulting from the addition of patches, and was multiplied by a Gaussian factor, 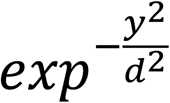 to concentrate the ring at the centre of the simulation box. The box circumference was 3000 nm, corresponding to a cell radius of ∼0.5 µm. In the wild-type condition, a reservoir of 4000 monomers was imposed such that no new monomers could be added until existing monomers depolymerised. Overexpression was modelled by removing this limit and the treadmilling mutant was modelled by increasing τ_hydrolysis_ tenfold to 1 s. All simulations represented 30 min of Z-ring condensation.

To test the importance of the antiparallel doublet interface, two control simulations were performed. In the first, only the C-C interface was attractive, while the N-N interface retained only WCA repulsion (ε = 5 k_BT_, r_cutoff_ = 0.1 σ, r_min_ = 0.1 σ). In the second, which tested a hypothetical parallel arrangement, the N-N interface was again repulsive and a short-range C-N attraction (ε = 10 k_BT_, r_cutoff_ = 0.4 σ, r_min_ = 0.1 σ) was introduced, coupling filaments treadmilling in the same direction.

Condensation trajectories were analysed in two ways. First, for every second of the trajectory, filaments were grouped into clusters connected by lateral interactions and the percentage of the circumference spanned by a continuous doublet cluster was calculated. Values were averaged over 15 replicas to give the mean trend. To account for strong temporal correlations, blocking analysis was performed on each replica between 20 and 29 min, and the mean of this period was taken as one sample per replica. Reported values are the mean and standard error of the mean across replicas. Second, ring thickness was measured by sampling the ring width at 10 positions around the circumference, filtering for laterally interacting filaments based on filament-filament distance and averaging across all slices. Mean thickness over time was calculated from 15 replicas for each simulation setup.

## Supplemental Information

**Figure S1.**
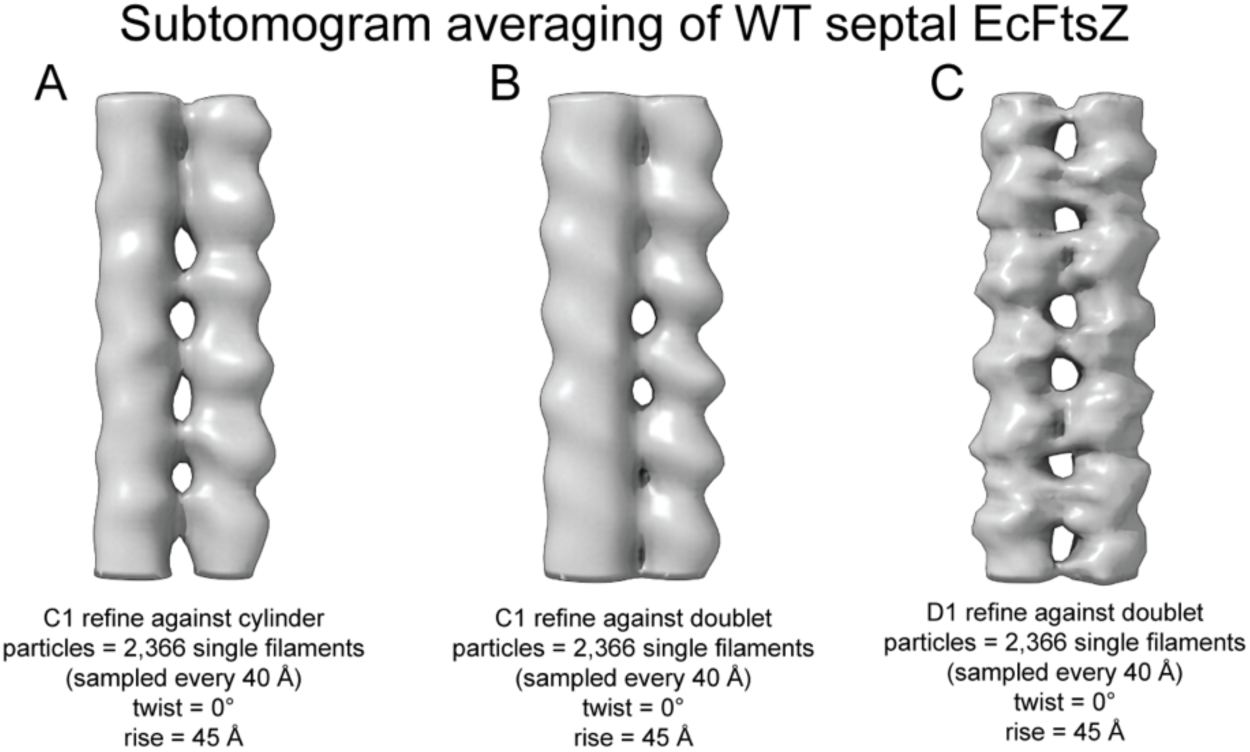
Subtomogram averaging FtsZ in unmodified “WT” C41(DE3) dividing *E. coli* cells. Maps are the results of refinements after picking all single filaments visible near invaginating septa. (A) Refinement against a cylinder as a reference model, applying a zero twist and 45 Å rise symmetry, demonstrates that filaments inside cells are mostly doublets. (B) Same helical parameters as for A but refined against a lowpass filtered double filament reconstruction from the FtsZ_D121A_ overexpression dataset to demonstrate that the EcFtsZ filaments are similar to the overexpressed filaments. (C) Same as B except D1 symmetry applied from the start to demonstrate that filaments in the non-overexpressed cells are antiparallel as well.

**Figure S2.**
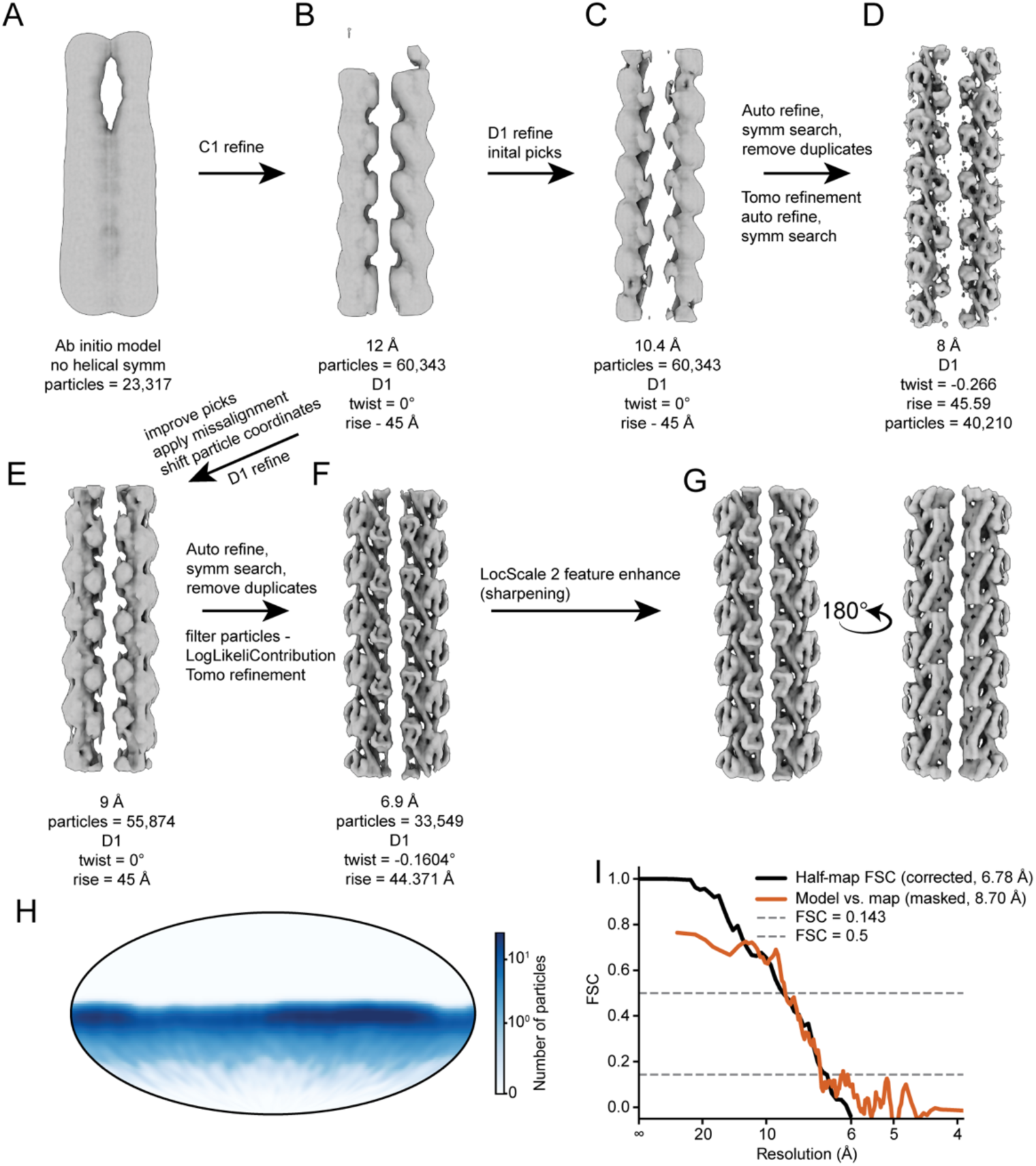
Subtomogram averaging (STA) processing workflow overview for FtsZ_D212A_. Corresponds to Figure 1E. (A) An initial model from a subset of traced filaments sampled every 40 Å and without helical or D1 symmetry applied. (B) First refinement in C1 with initial basic helical parameters of 0° twist and 45 Å rise applied. (C) First refinement applying D1 symmetry prior to helical symmetry search. (D) An 8 Å refinement prior to cleaning particle picks. The map shows secondary structure features but is noisy due to the large number of unaligned particles and duplication issues. The processing workflow suffered from a failure to identify suitable classes via 3D classification. (E) Map resulting from a first refinement following initial particle cleanup and extraction of coordinates from alignments produced by MissAlignment^1^. The volume shown in B was used as an initial reference. (F) The best reconstruction produced following multiple refinements and particle curations, and finally CTF refinement. No improvements could be made through polishing or further refinements. (G) LocScale-2.0^2^ – sharpened map (model-free). (H) Particle orientations shown on a Mollweide projection. Plotted using starparser^3^. (I) Corrected half-map FSC and model-map FSC curves.

**Figure S3.**
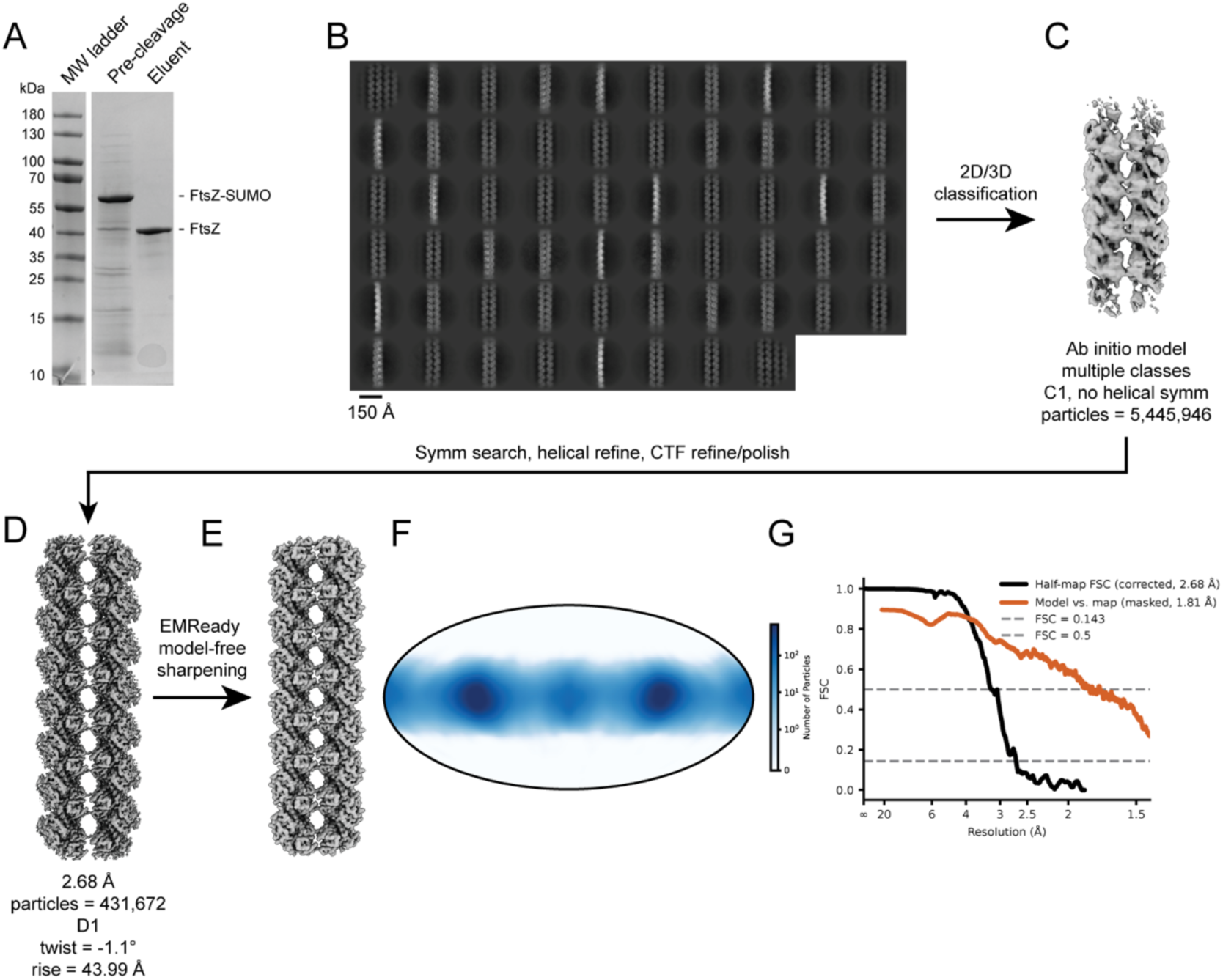
Overview of EcFtsZ single particle cryo-EM processing workflow. Corresponds to Figure 1J. (A) Example SDS-PAGE gel from protein purification. (B) 2D classes from the processing workflow. (C) *Ab initio* model used to initiate 3D refinement steps. (D) Final sharpened reconstruction. (E) EMReady^4^-sharpened final map (model-free). (F) Particle orientations shown on a Mollweide projection. Plotted using starparser^3^. (G) Corrected half-map FSC and model-map FSC curves.

**Figure S4.**
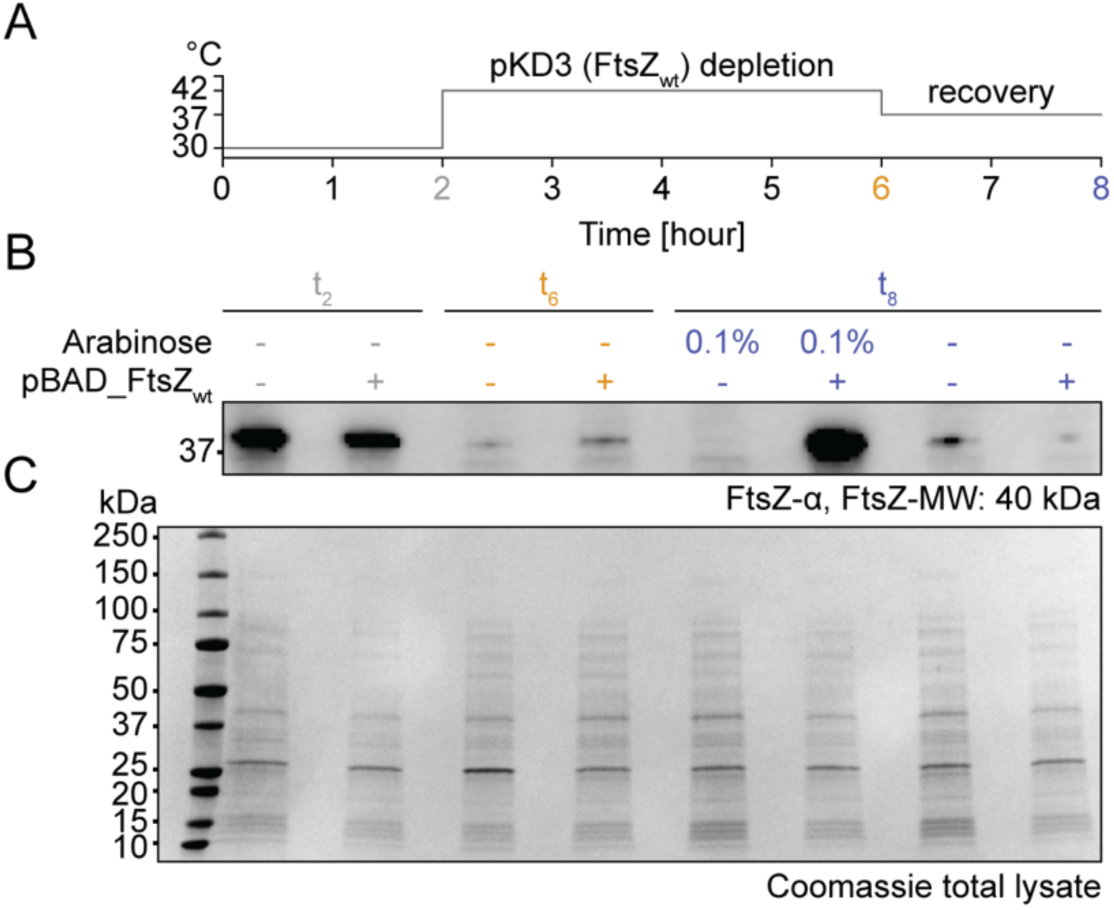
Western blot of FtsZ levels in *ftsZ*(WT) ts strain JKD7-1. (A) Experiment timeline. (B) Western blot showing depletion of FtsZ protein following 4 hours at the non-permissive temperature and a 2-hour recovery. (C) Loading control (Coomassie stained gel) of the samples used for western blotting in B. Corresponds to Figures 2B & 4C.

**Figure S5.**
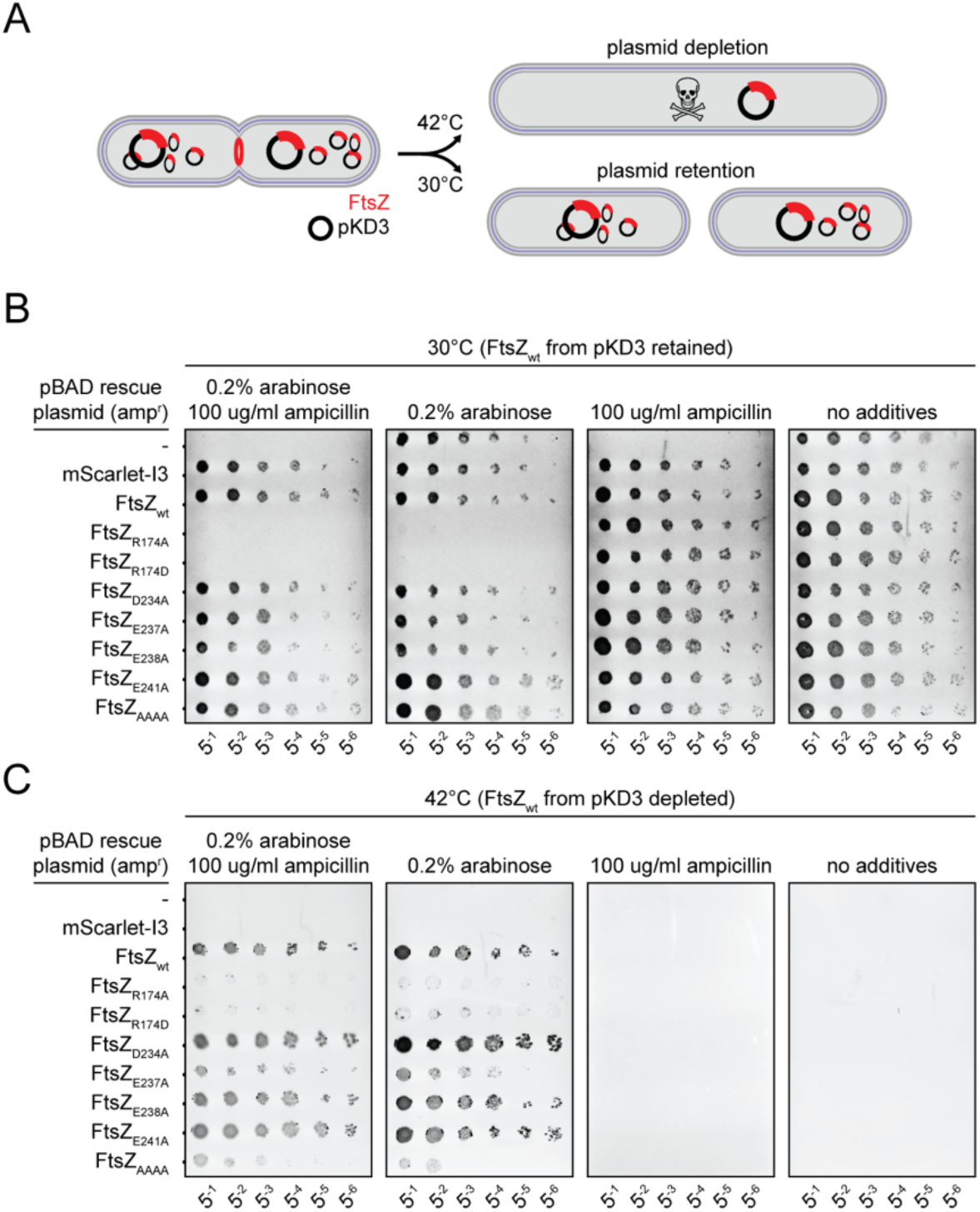
Extended dilution spot assay of EcFtsZ doublet interface mutants. Corresponds to Figure 2B. (A) Schematic of the genetic setup demonstrating the retention (B) and loss of the pKD3 *ftsZ(WT) ts* plasmid (C), which makes the pBAD rescue plasmid essential in the absence of pKD3 (Table S1).

**Figure S6.**
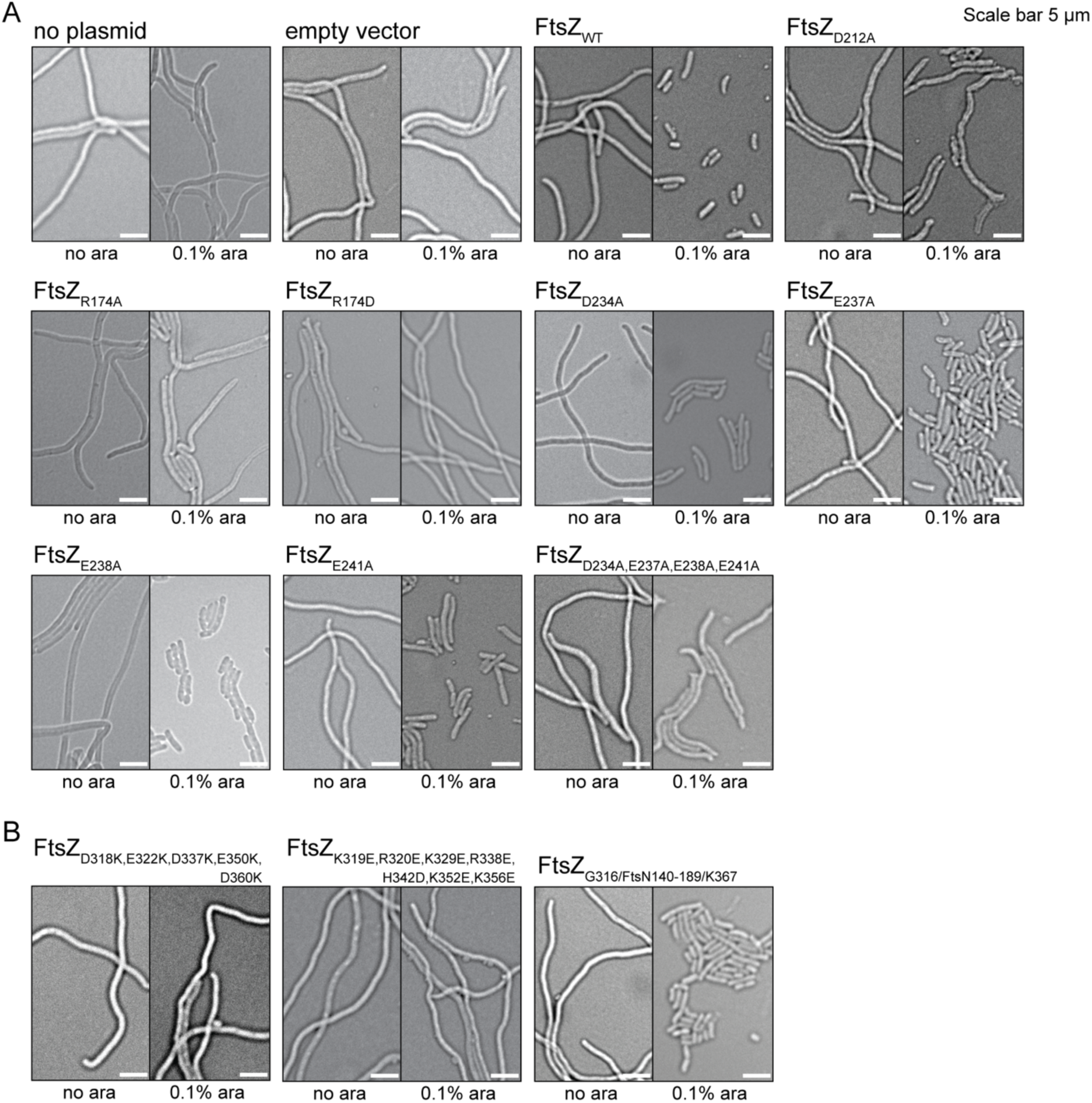
Phase contrast light microscopy of all EcFtsZ doublet interface mutants and some CTT mutants. Each condition is shown in two images – with and without arabinose induction of the pBAD rescue plasmid. Doublet mutants (A) correspond to Figure 2B and mutate residues either side of the doublet interface. (B) CTT mutants excluding the FtsA binding C-terminal Peptide (CTP) are equivalent to those shown in Figure S15. Mutations (from left to right) result in an entirely positive CTT, entirely negative or an FtsN insertion. This mutant does not rescue at 42°C but has no division phenotype when grown at 37°C following a 42°C heat-shock for 4 hours, suggesting this mutant is viable but temperature sensitive (see Figure S15D).

**Figure S7.**
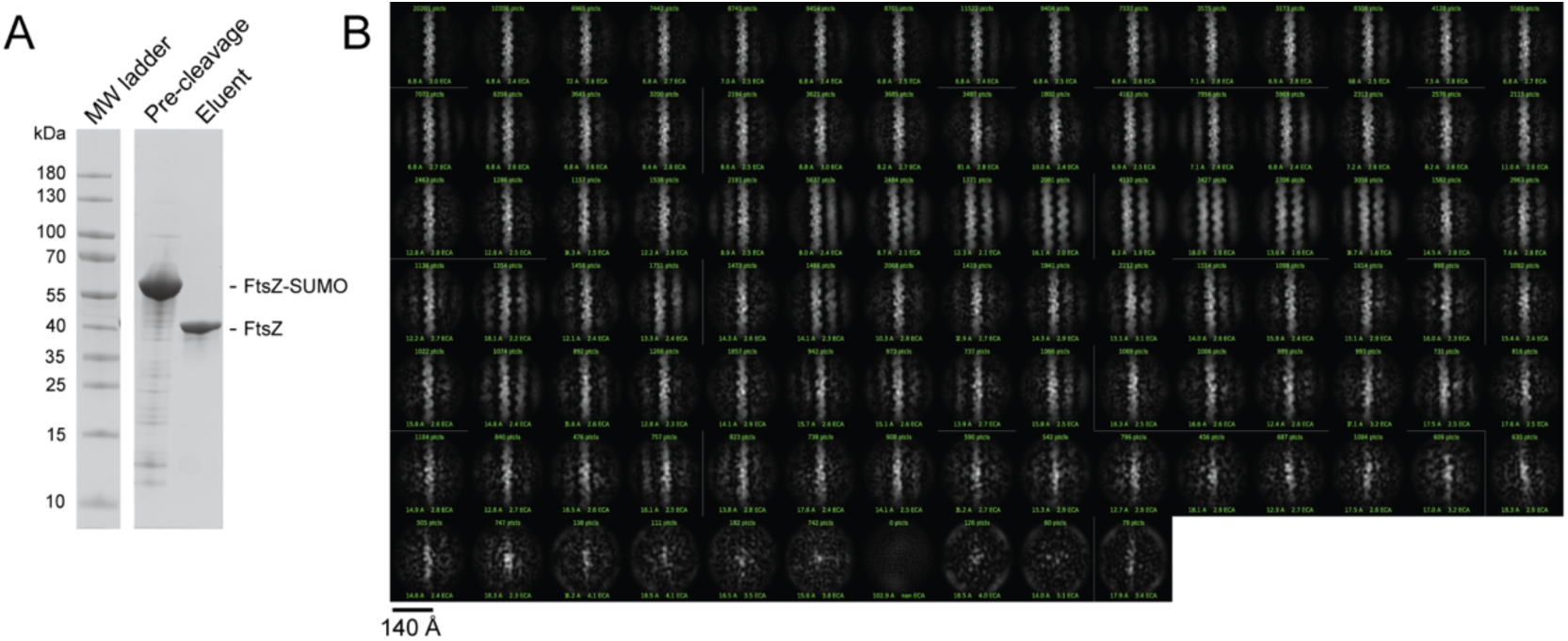
EcFtsZ_R174D_ single particle cryo-EM processing. Corresponds to Figure 2E. (A) SDS-PAGE gel from protein purification. (B) 2D classes from crowded EM micrographs. Neighbouring single filaments can be seen in some classes, likely due to nematic ordering at high densities, but no examples of double filaments could be resolved.

**Figure S8.**
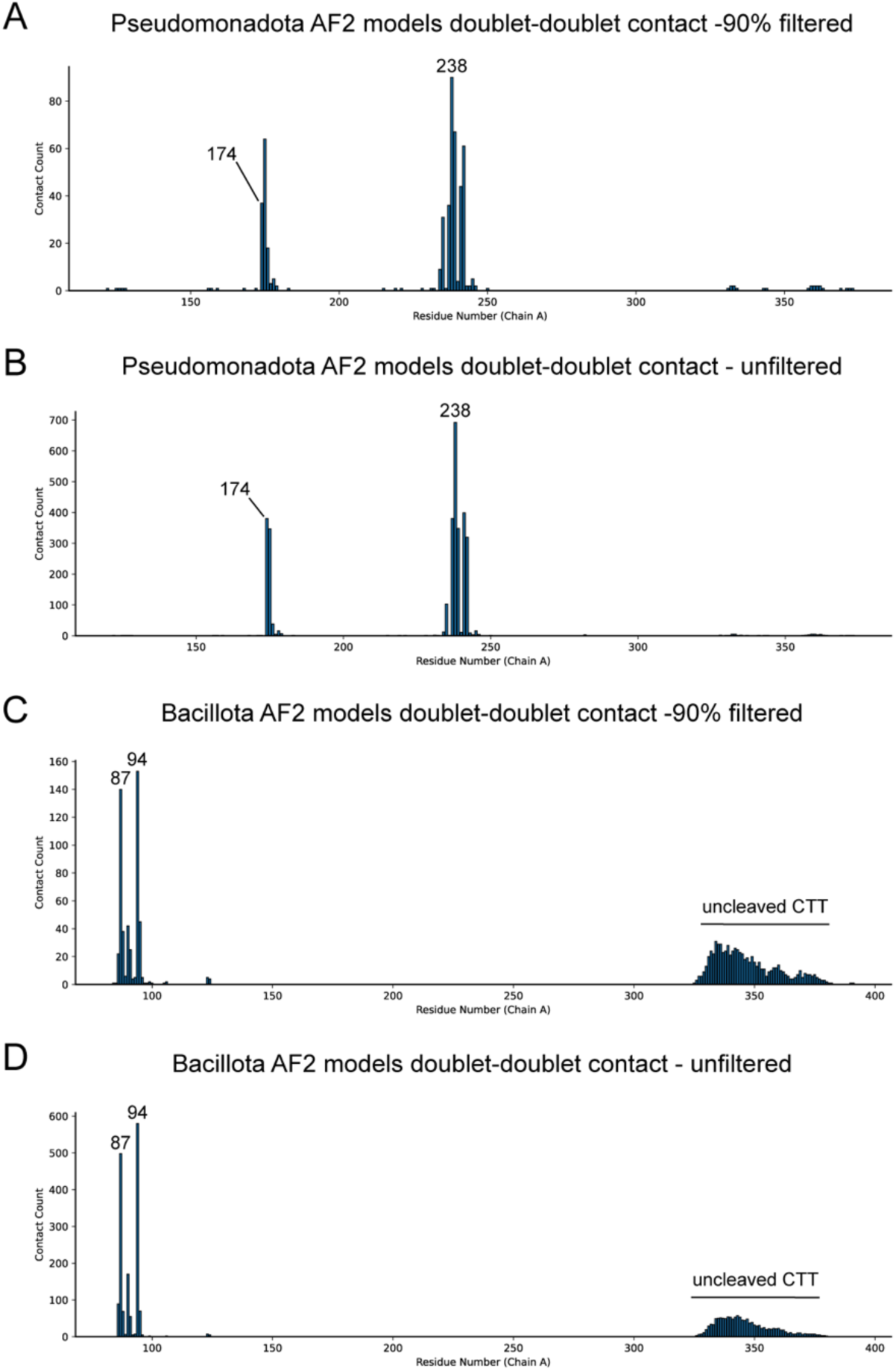
Histogram contact plots for the FtsZ AF2 analyses. Corresponds to Figure 2H-J (A, B) and Figure 3EG (C, D).

**Figure S9.**
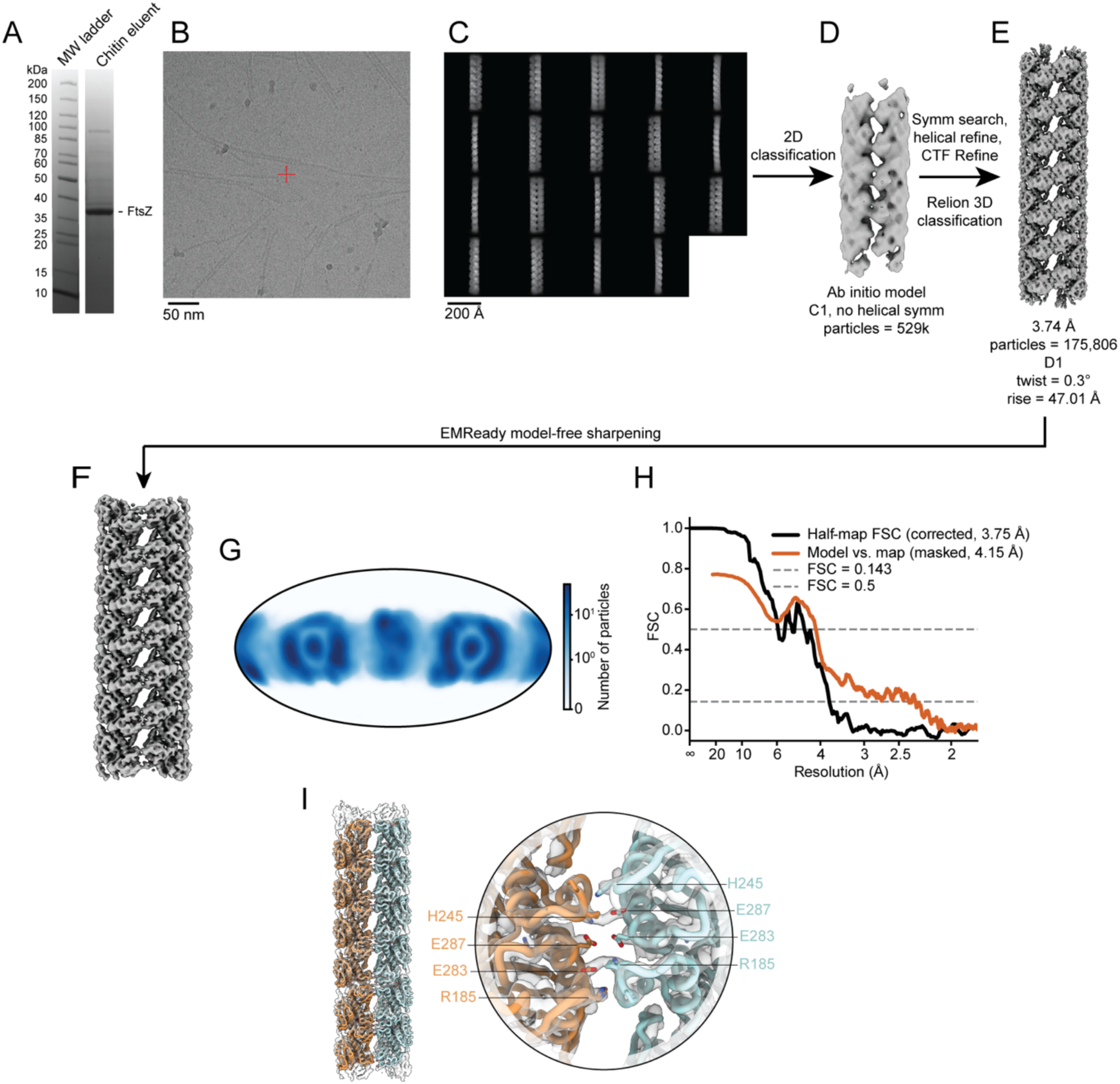
Overview of TmFtsZ single particle cryo-EM processing workflow. Corresponds to Figure 4F. (A) SDS-PAGE gel from protein purification of TmFtsZ. (B) Example micrograph showing double and single filaments used to extract particles for further processing. (C) Example 2D classes from the processing workflow. (D) Initial model used to align initial refinements. (E) Final and best reconstruction. (F) EMReady^4^ model-free sharpened map. (G) Orientations projected onto a Mollweide plot using starparser^3^. (H) Corrected half-map FSC and model-map FSC curves. (I) TmFtsZ atomic model shown in the transparent single particle cryo-EM map (left) beside a close-up of the doublet interface (right).

**Figure S10.**
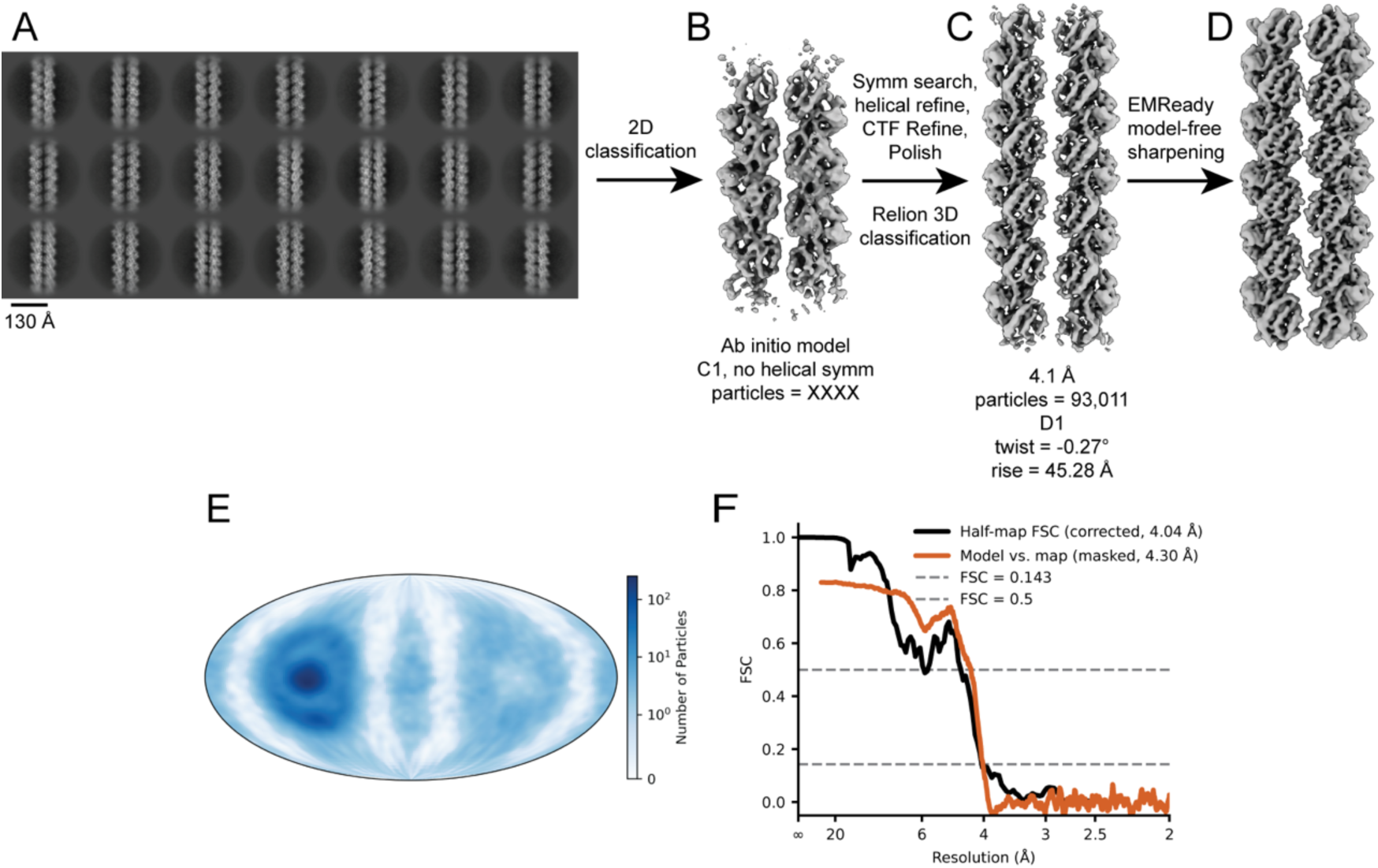
Overview of BsFtsZ single particle cryo-EM processing workflow. Corresponds to Figure 3B. (A) Example 2D classes from the processing workflow. (B) Initial model used for initial refinements. (C) Final and best reconstruction. (D) EMReady^4^ model-free sharpened map. (E) Orientations projected onto a Mollweide plot using starparser^3^. (F) Corrected half-map FSC and model-map FSC curves.

**Figure S11.**
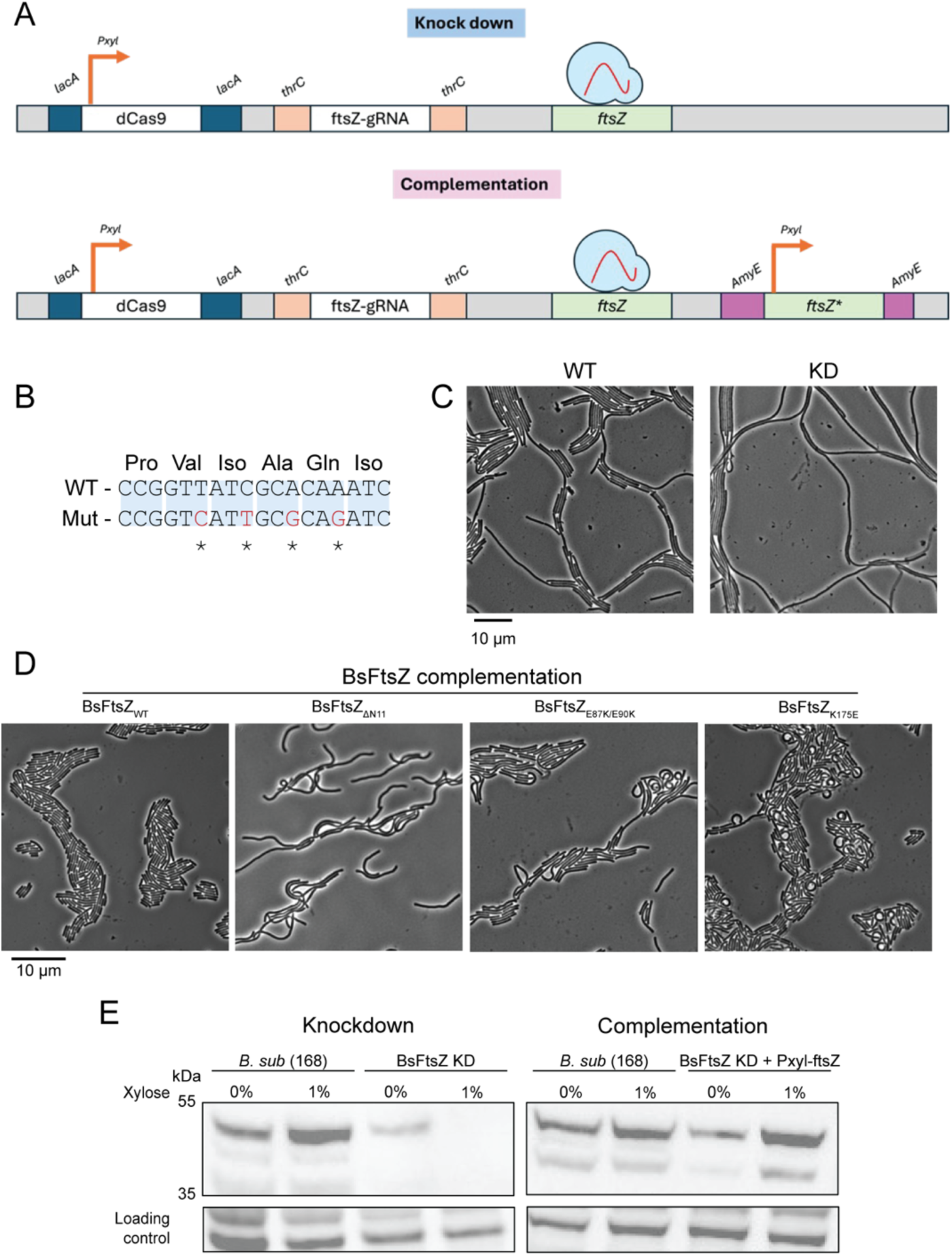
*B. subtilis* dCas9 *ftsZ* knockdown strains. A) Genetic setup. B) “Immunity” sequences used for *ftsZ\**, the copy that is expressed when xylose is added, which also repressed *ftsZ(WT)* through the action of dCas9 and the gRNA. C) KD test (no *ftsZ\**). D) Light microscopy of the strains used in the spot dilution assay (Figure 3F). E) Western blot showing the KD of FtsZ and the production of FtsZ*. Corresponds to Figure 3F.

**Figure S12.**
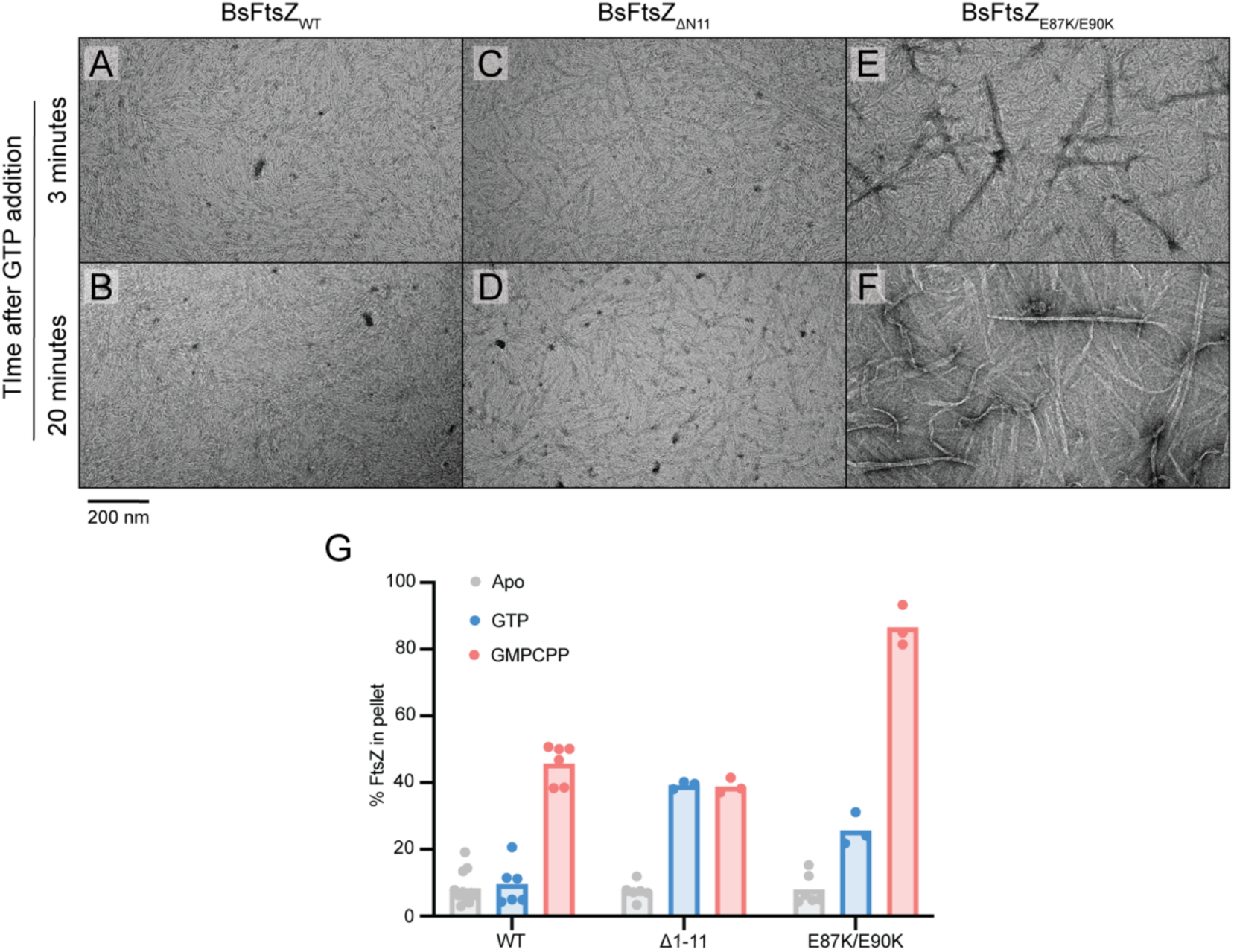
BsFtsZ negative stain electron microscopy imaging and pelleting assays. Corresponds to Figure 4FG. Purified BsFtsZ 3 minutes (A) and 20 minutes (B) after polymerisation with GTP. Purified BsFtsZ_Δ1-11_ 3 minutes (C) and 20 minutes (D) after polymerisation with GTP. Purified BsFtsZ_E87K/E90K_ 3 minutes (E) and 20 minutes (F) after polymerisation with GTP. (G) SDS-PAGE based pelleting assay of BsFtsZ WT and mutant filaments in the presence of GTP and GMPCPP.

**Figure S13.**
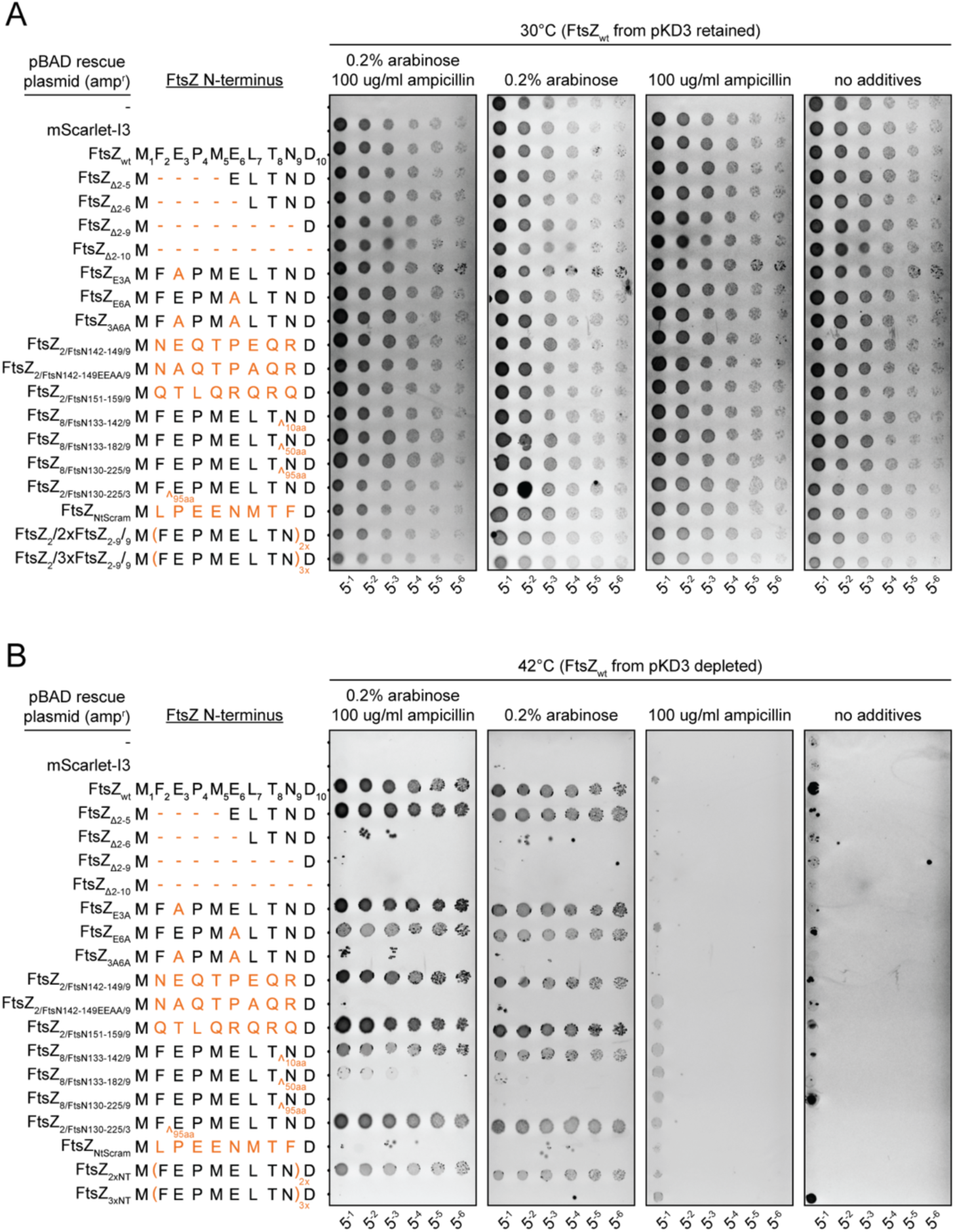
Extended dilution spot assay of EcFtsZ N-terminal tail (NTT) mutants. Corresponds to Figure 4C. See Figure S5A for schematic of the genetic setup. (A) Spot assays at permissive temperature with a retained pKD3 *ftsZ(WT)* ectopic plasmid. (B) Spot assays at a non-permissive temperature with a loss of pKD3 *ftsz(WT)* ectopic plasmid. A different FtsN IDR sequence (FtsN_151-159_) was used to replace the N-terminal residues 2-9 of EcFtsZ . The replacement spots at 42 but in Figure S14 (second from bottom, left side) the strain has a clear division phenotype. This is the second discrepancy between the light microscopy and spot assay that we have observed in this study, the other being the temperature sensitive nature of the the FtsZ_G316/FtsN140-189/K367_ in Figure S6 and S15D.

**Figure S14.**
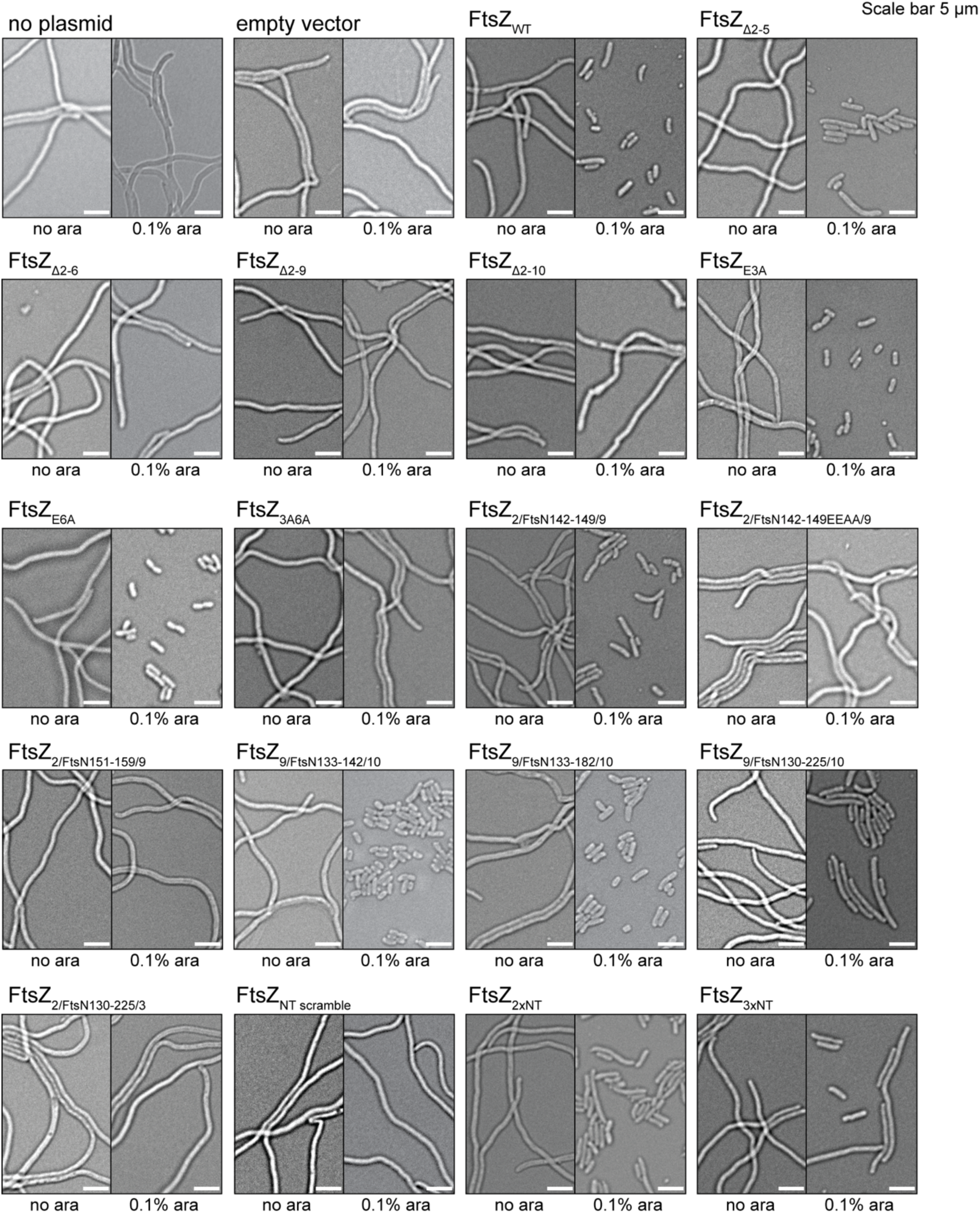
Phase contrast light microscopy of all EcFtsZ NTT mutants. The no plasmid, empty vector and FtsZ_WT_ are reused from Figure 2B and all strains correspond to Figure S13. Each condition is shown in two images – with and without arabinose induction. Refer to Figure S13 for description of mutants FtsZ_NT scramble_, FtsZ_2xNT_ and FtsZ_3xNT_. FtsZ_2/FtsN151-159/9_ (second from bottom, left side) has a division phenotype but in Figure S13, the same mutant complements FtsZ depletion.

**Figure S15.**
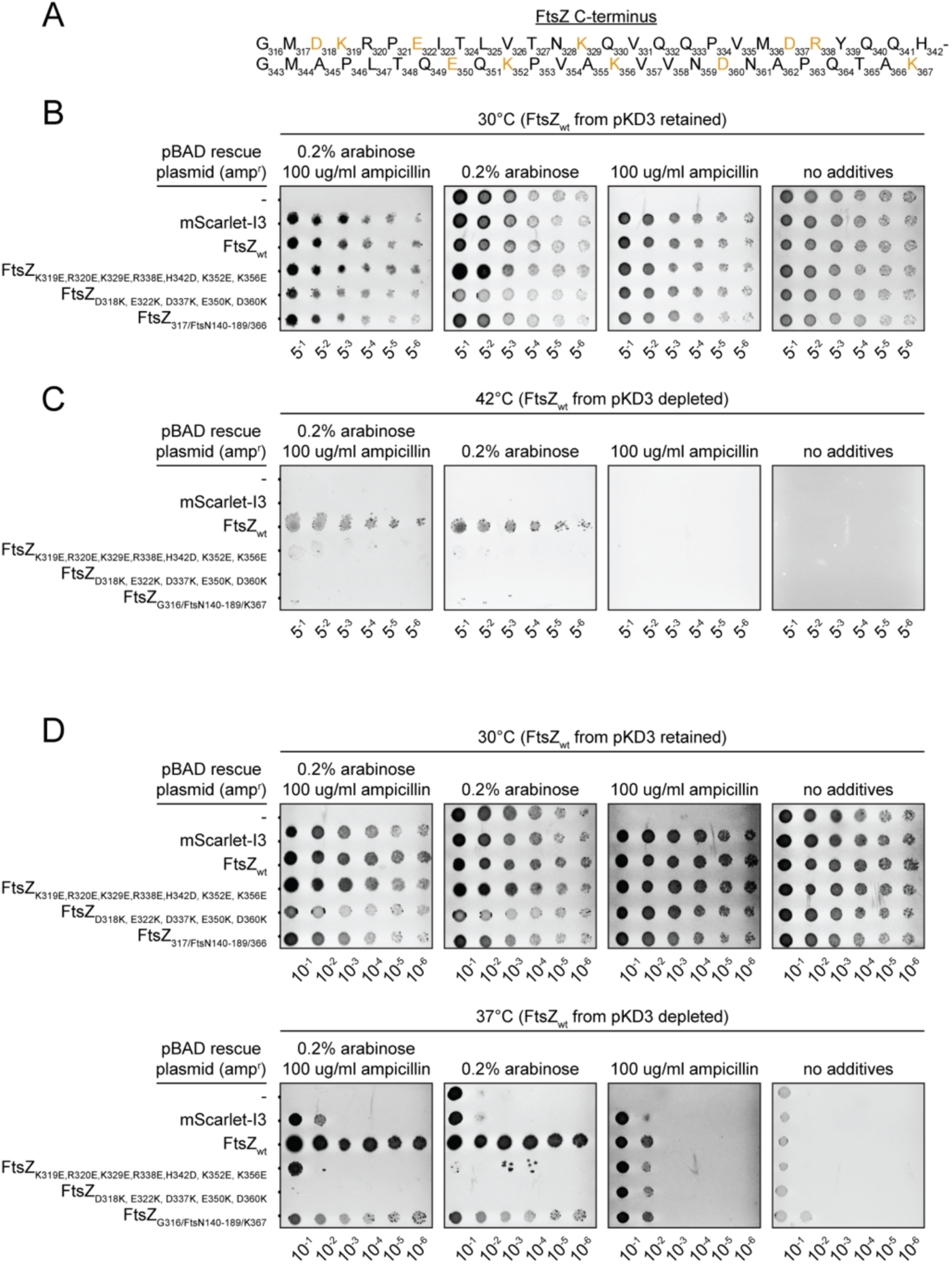
Dilution spot assay of EcFtsZ CTT mutants. Schematic of the genetic setup in Figure S5A. (A) EcFtsZ CTT with charged residues highlighted. (B) Spot assays at a permissive temperature with a retained pKD3 *ftsZ(WT)* ectopic plasmid. (C) Spot assays at a non-permissive temperature with a loss of pKD3 *ftsz(WT)* ectopic plasmid. (D) Spot tests done using the growth protocol used for light microscopy experiments using the same strains as in panels B and C. Note that FtsZ_G316/FtsN140-189/K367_ doesn’t grow when recovered at 42°C but does so at 37°C. The final spot shows a FtsN CTT replacement which cannot compliment at 42°C (panel C) in spot dilution assays but when heat-shocked at 42°C for 4 hours and allowed to recover at 37°C for a further two hours (see Figure S5A) shows no division phenotype (Figure S6B).

**Figure S16.**
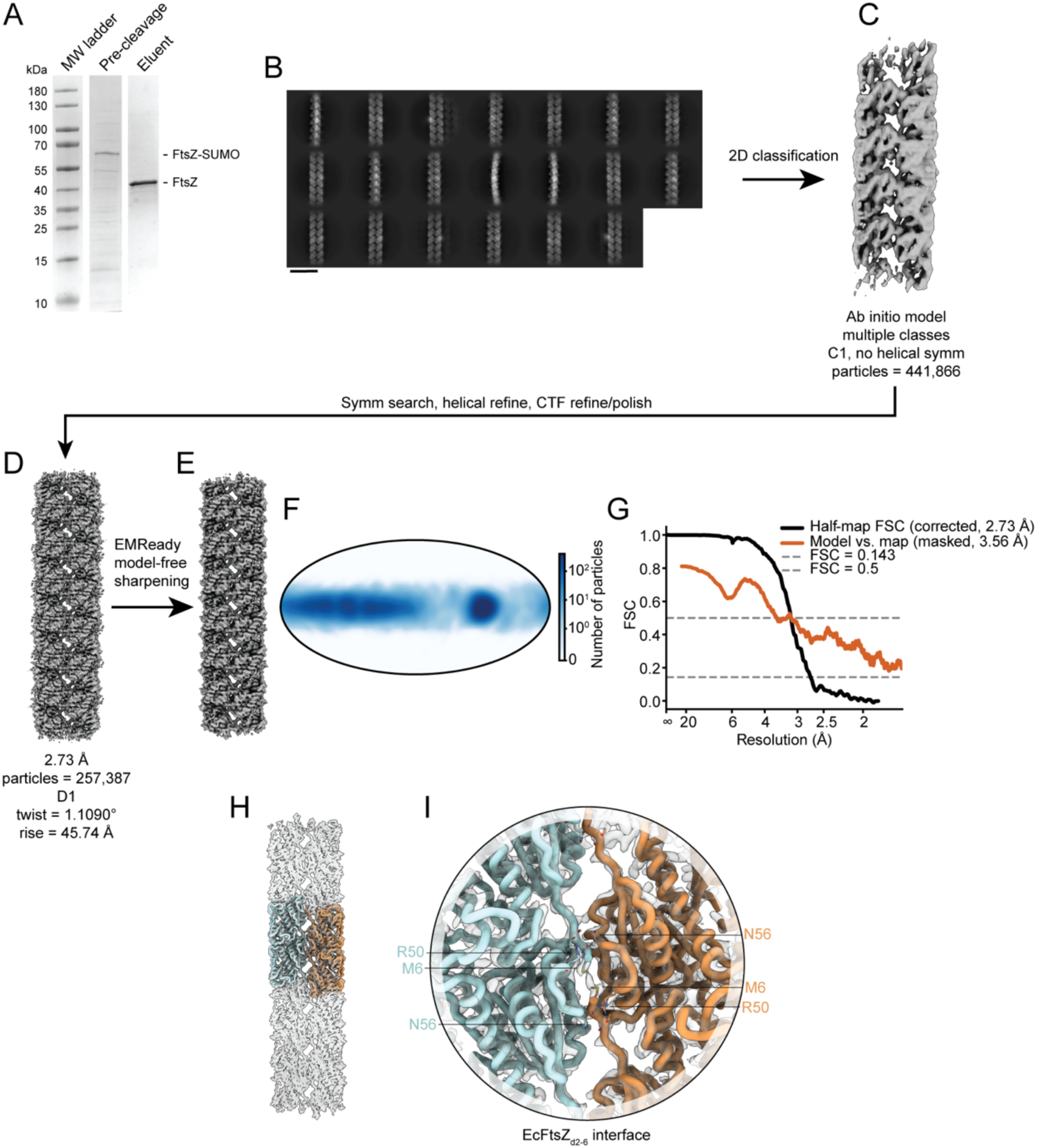
Overview of EcFtsZ_Δ2-6_ single particle cryo-EM processing workflow. Corresponds to Figure 4E. (A) SDS-PAGE gel from protein purification of EcFtsZ_Δ2-6_. (B) Example 2D classes from the processing workflow. (C) Initial model used for initial refinements. (D) Final and best reconstruction. (E) EMReady^4^ model-free sharpened map. (F) Orientations projected onto a Mollweide plot using starparser^3^. (G) Corrected half-map FSC and model-map FSC curves. (H) EcFtsZ_Δ2-6_ model shown in the transparent single particle cryo-EM map. (I) A close-up of the N-N doublet interface.

**Figure S17.**
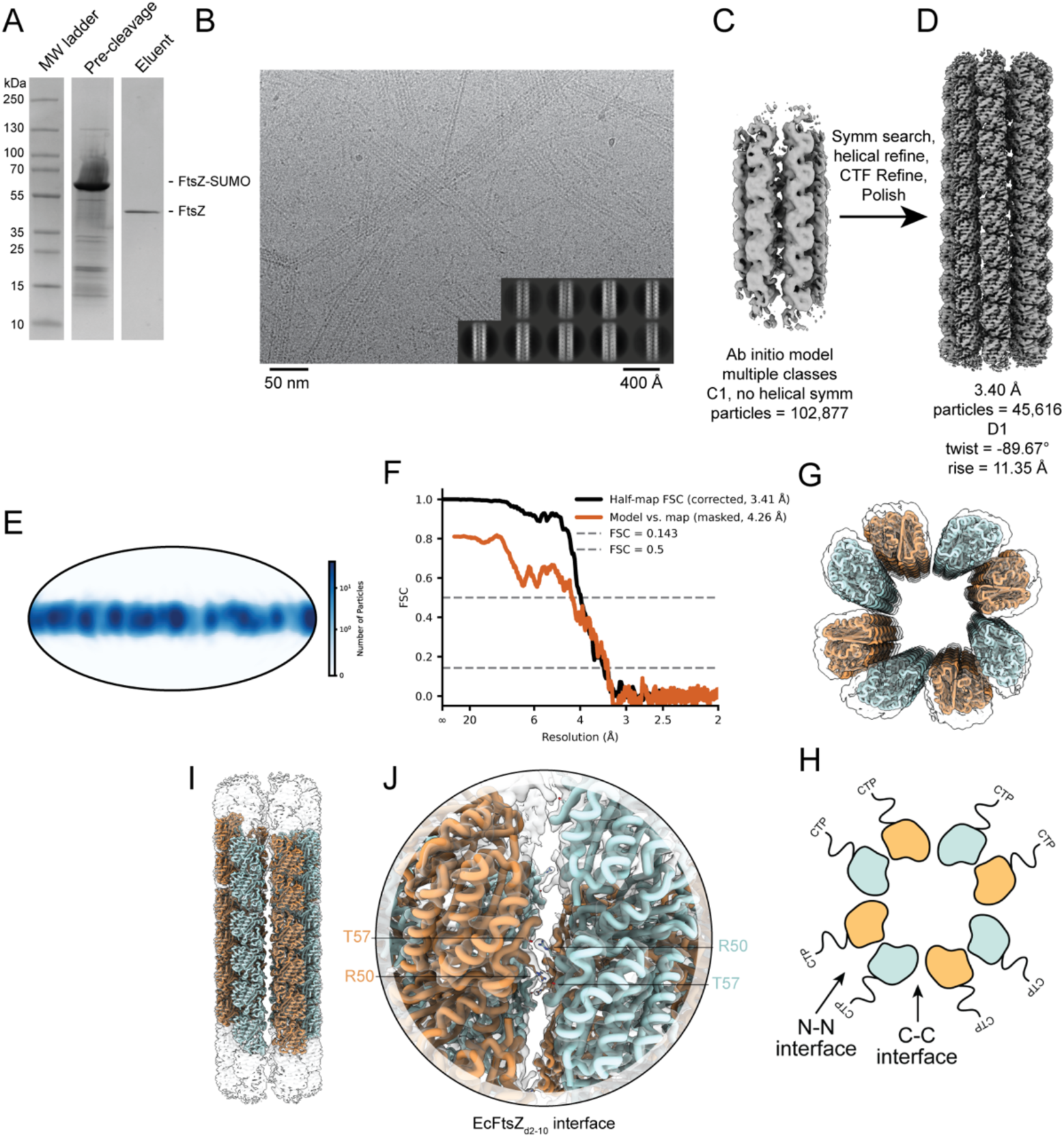
Overview of EcFtsZ_Δ2-10_ single particle cryo-EM processing workflow. Results not shown in main text figures. (A) SDS-PAGE gel from protein purification of EcFtsZ_Δ2-10_. (B) Example electron micrograph and 2D classes from the processing workflow. (C) Initial model used for initial refinements. (D) Final and best reconstruction. (E) Orientations projected onto a Mollweide plot using starparser^3^. (F) Corrected half-map FSC and model-map FSC curves. (G) EcFtsZ_Δ2-10_ model shown in the transparent single particle cryo-EM map (top view). (H) Schematic of top view shown in G, with direction of unresolved C-terminal tail (CTT) and both C-C and N-N interfaces holding the hollow cylinder together indicated. (I) Same as G but a side-view. (J) A close-up of the N-N interface.

**Figure S18.**
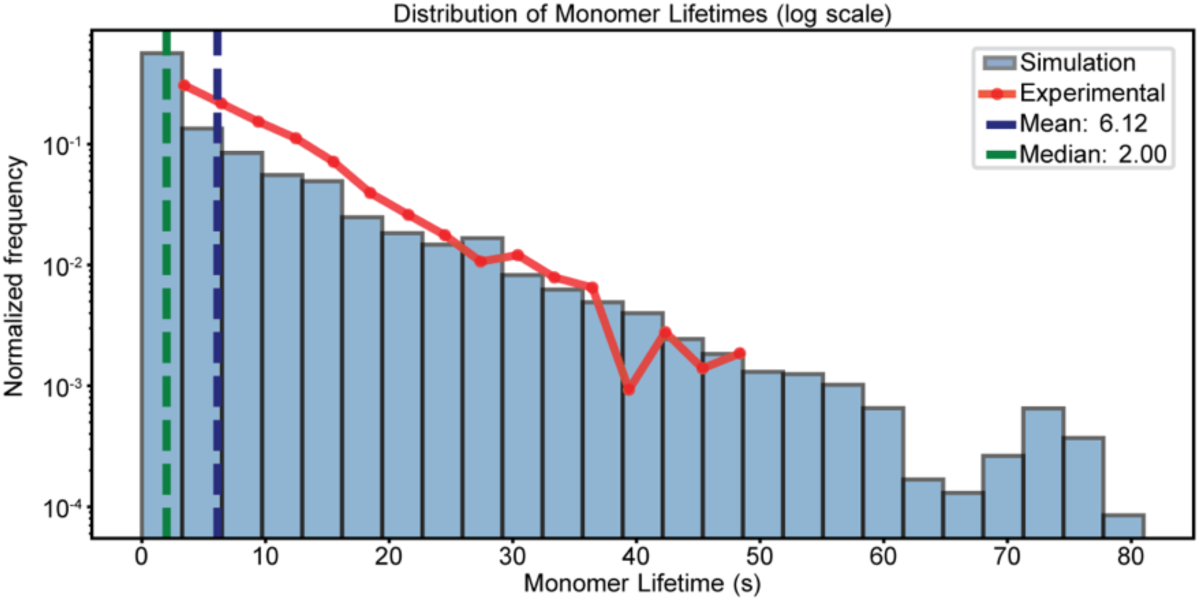
Monomer lifetimes in simulations closely match those in *in vivo* data from FtsZ treadmilling in *B. subtilis*. Corresponds to Figure 5. Experimental data taken from previous *in vivo* single molecule studies^5^.

**Figure S19.**
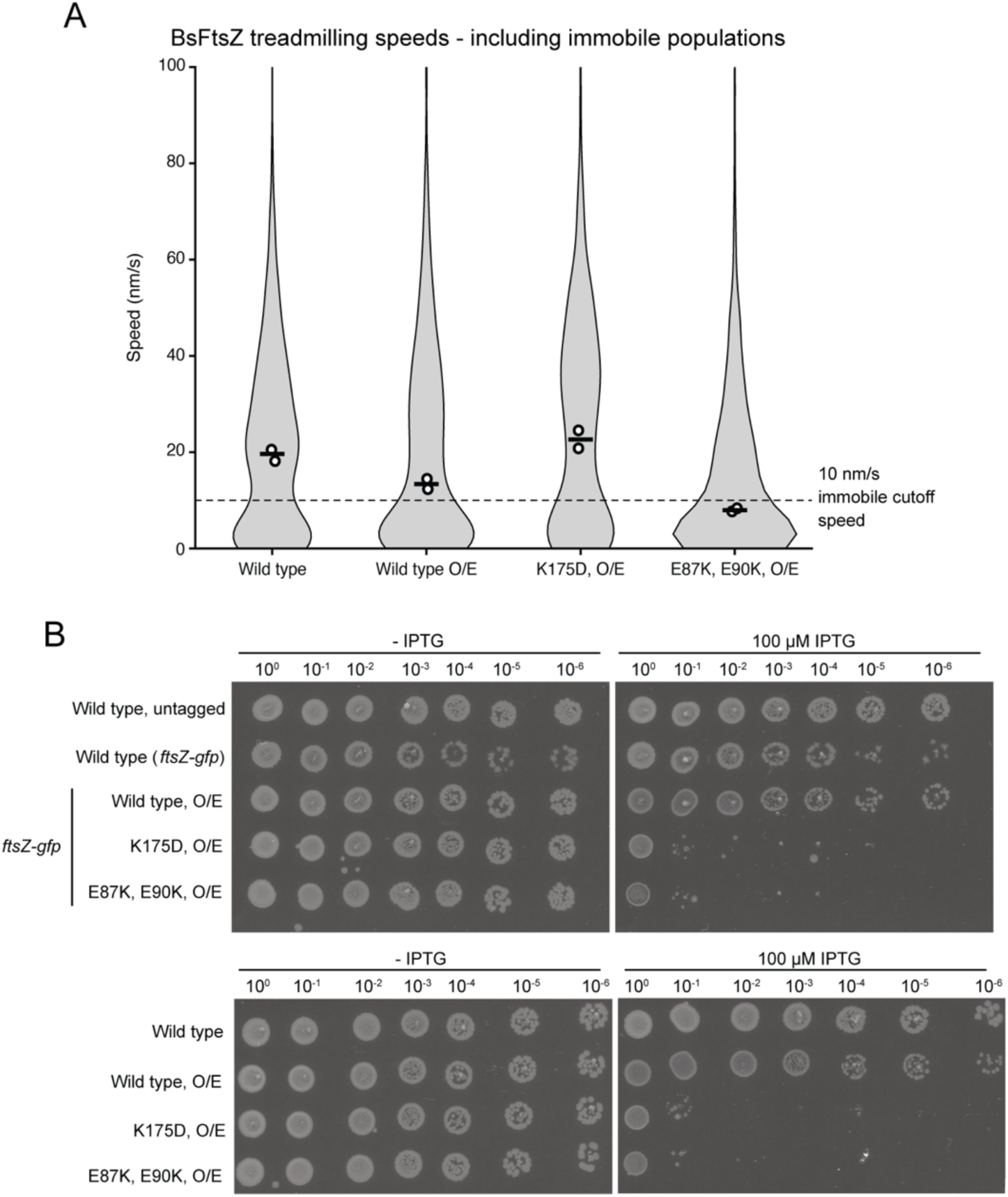
Measuring FtsZ treadmilling speeds in *B. subtilis* with VerCINI. A) The same treadmilling speed data as in Figure 5C, here also including tracks below the immobile cutoff speed (< 10 nm/s) not shown in Figure 5C. B) Dilution spot assay of the mutants used in Figure 5CD and panel A with and without GFP fusion showing dominant negative phenotypes of K175D and E87K/E90K are not due to GFP. Corresponds to Figure 5CD.

**Figure S20.**
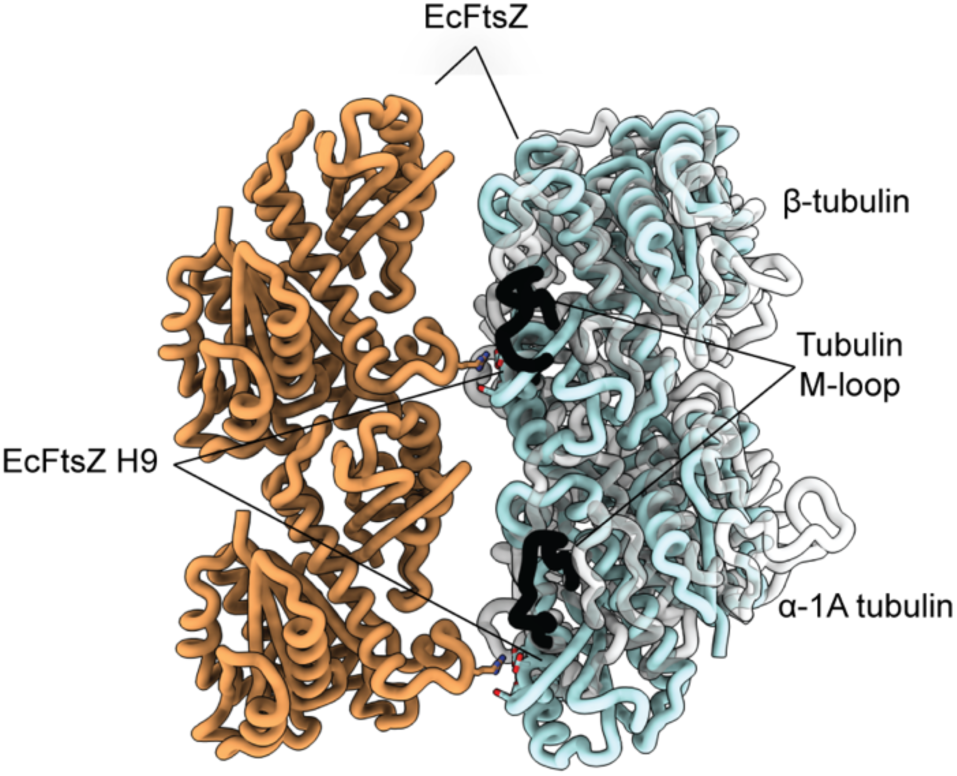
The position of the tubulin M-loop with respect to EcFtsZ lateral interface motifs. The first tubulin heterodimer structure (PDB 1TUB) from *Sus scrofa*^6^ was compared by aligning the Δ-tubulin chain to the top right subunit of the EcFtsZ doublet structure (this study). EcFtsZ interfacial residues are shown as sticks highlighting H9 which contributes negative charges to the interface. The Δ-tubulin subunit M-loop highlighted in black (responsible for holding microtubule protofilaments together laterally) protrudes out from the equivalent helix to H9. Both EcFtsZ and tubulin appear to use a very similar region of the tubulin fold for engaging in lateral interactions.

**Figure S21.**
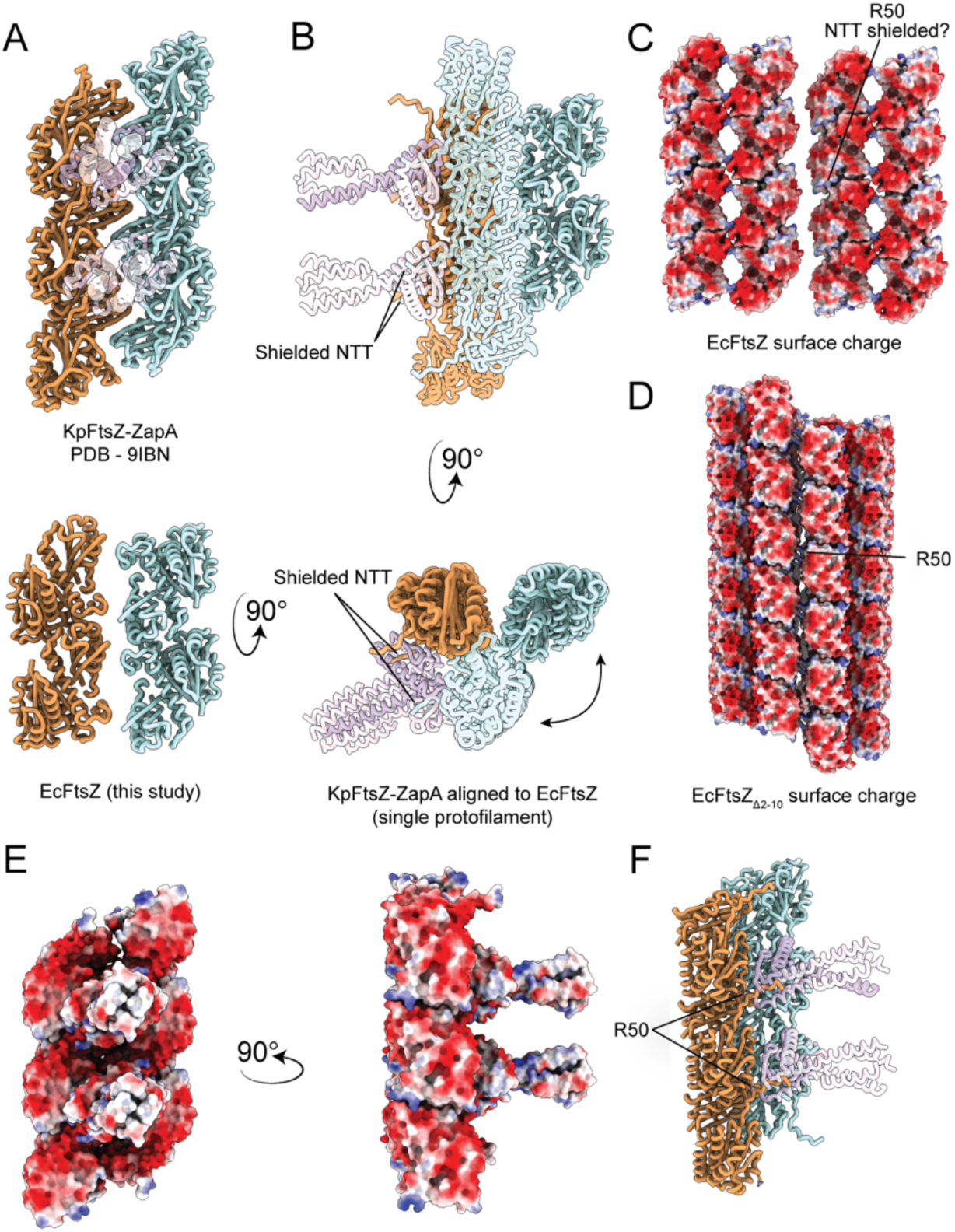
ZapA-bound FtsZ (PDB 9IBN)^7^ vs unbound (this study). (A) View of antiparallel double filament from KpFtsZ-ZapA (PDB 9IBN) top and EcFtsZ bottom. (B) KpFtsZ-ZapA (PDB 9IBN) aligned to EcFtsZ doublet left protofilament (orange) side and top view showing a substantial double filament rearrangement when bound to ZapA. (C) Surface charge of EcFtsZ doublet-doublet (Figure 4A) showing highly negatively charged surface with exposed R50 (presumably shielded by the NTT). (D) Same surface charge of EcFtsZ_Δ2-10_ mutant where R50 is no longer shielded and interfaces between protofilaments. (E) Surface charge of KpFtsZ-ZapA (PDB 9IBN) showing a similarly negatively charged surface area. (F) R50, which points towards neighbouring doublets in the EcFtsZ structure (this study), is rotated and points towards the cytoplasmic facing side of the double filament in the KpFtsZ-ZapA (PDB 9IBN) structure.

**Figure S22.**
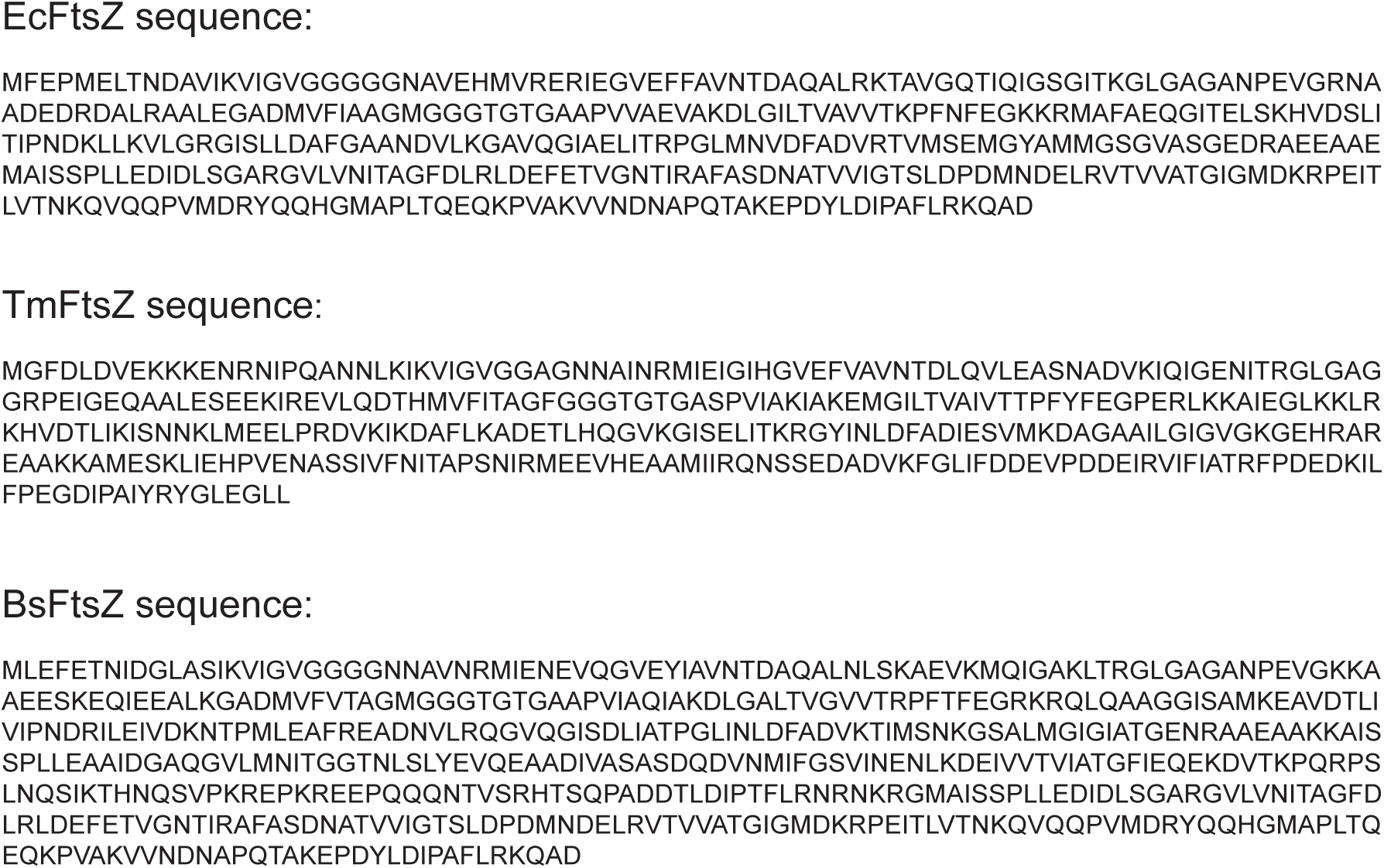
Protein sequences of FtsZs used in this study.

## Supplementary Tables

**Table S1.** Plasmids used in this study.

| Name | Content | Source |
|---|---|---|
| pLR241 | pET-24b(+)_10xHis_SUMO_FtsZ<sub>Δ2-6</sub> | Löwe group collection |
| pLR260 | pET-24a(+)_10xHis_SUMO_FtsZ | Löwe group collection |
| pLR263 | pET-24a(+)_10xHis_SUMO_FtsZR174D | Löwe group collection |
| pLR265 | pET-24(+)_10xHis_SUMO_FtsZ<sub>Δ2-10</sub> | Löwe group collection |
| pLR105 | pBAD_FtsZ | Löwe group collection |
| pLR140 | pBAD_FtsZ_R174A | Löwe group collection |
| pLR168 | pBAD_FtsZ_R174D | Löwe group collection |
| pLR182 | pBAD_ FtsZ_Δ2-10 | Löwe group collection |
| pLR186 | pBAD_ FtsZ_Δ2-6 | Löwe group collection |
| pLR231 | pBAD_FtsZ_E3A | Löwe group collection |
| pLR246 | pBAD_FtsZ_K319E_R320E_K329E_R338E_H342D_ K352E_K356E | Löwe group collection |
| pLR247 | pBAD_FtsZ<sub>8</sub>/FtsN<sub>130-225</sub>/<sub>9</sub> | Löwe group collection |
| pLR248 | pBAD_ FtsZ<sub>8</sub>/FtsN<sub>133-142</sub>/<sub>9</sub> | Löwe group collection |
| pLR250 | pBAD_FtsZ<sub>8</sub>/FtsN<sub>133-182</sub>/<sub>9</sub> | Löwe group collection |
| pLR253 | pBAD_FtsZ<sub>317</sub>/FtsN<sub>140-189</sub>/<sub>366</sub> | Löwe group collection |
| pLR257 | pBAD_FtsZ_D318K_E322K_D337K_E350K_D360K | Löwe group collection |
| pLR274 | pBAD_ FtsZ_E3A_E6A | Löwe group collection |
| pLR275 | pBAD_ FtsZ_E237A | Löwe group collection |
| pLR276 | pBAD_ FtsZ_E238A | Löwe group collection |
| pLR279 | pBAD_ FtsZ_E234A | Löwe group collection |
| pLR280 | pBAD_ FtsZ_E241A | Löwe group collection |
| pLR281 | pBAD_ FtsZ_E6A | Löwe group collection |
| pLR282 | pBAD_FtsZ<sub>2</sub>/2xFtsZ<sub>2-9</sub>/<sub>9</sub> | Löwe group collection |
| pLR283 | pBAD_ FtsZ<sub>2</sub>/3xFtsZ<sub>2-9</sub>/<sub>9</sub> | Löwe group collection |
| pLR284 | pBAD_FtsZ<sub>2</sub>/FtsN<sub>130-225</sub>/<sub>3</sub> | Löwe group collection |
| pLR285 | pBAD_ FtsZ<sub>2</sub>/FtsN<sub>142-149</sub>/<sub>9</sub> | Löwe group collection |
| pLR286 | pBAD_ FtsZ_Δ2-9 | Löwe group collection |
| pLR287 | pBAD_FtsZ<sub>2</sub>/scrambled/<sub>9</sub> | Löwe group collection |
| pLR288 | pBAD_ FtsZ_Δ2-5 | Löwe group collection |
| pLR289 | pBAD_ FtsZ_E234A_ E237A_E238A_E241A | Löwe group collection |
| pLR290 | pBAD_ FtsZ<sub>2</sub>_FtsN<sub>142-149</sub>_E143A_E147A/<sub>9</sub> | Löwe group collection |
| pLR292 | pBAD_ FtsZ<sub>2</sub>/FtsN<sub>151-158</sub>/<sub>9</sub> | Löwe group collection |
| pLR293 | pBAD_mScarlt-I3 | Löwe group collection |
| pMZ120 | pBAD_FtsZ_D212A | Löwe group collection |
| pSZ7 | pET15b_TmFtsZ | Szwedziak *et al.*,<sup>8</sup> Löwe group collection |
| pUMS | pET-SUMO | Wehenkel group collection |
| pUMS463 | pET-SUMO BsFtsZ | Wehenkel group collection |
| pUMS544 | pET-SUMO BsFtsZ<sub>E87K/E90K</sub> | Wehenkel group collection |
| pUMS494 | pET-SUMO BsFtsZ<sub>Δ1-11</sub> | Wehenkel group collection |
| pUMS500 | pJMP3-thrC::Pveg-sgRNA_ftsZ | Wehenkel group collection |
| pUMS522 | pSweet-Pxyl-FtsZ_WT | Wehenkel group collection |
| pUMS526 | pSweet-Pxyl-FtsZ_gRNAMut | Wehenkel group collection |
| pUMS530 | pSweet-Pxyl-FtsZ_ΔN11 | Wehenkel group collection |
| pUMS532 | pSweet-Pxyl-FtsZ_K175E | Wehenkel group collection |
| pUMS533 | pSweet-Pxyl-FtsZ_E87K+E90K | Wehenkel group collection |
| pGL004 | *(amyE::spc Phyperspank) bla* | Addgene ID: 215336<sup>9</sup>. Holden group |
| pDMR1 | *(amyE::spc Phyperspank-ftsA) bla* | NEBuilder reaction using oDMR17, oDMR18 and pGL004 template DNA (fragment 1); oDMR19, oDMR20 and MDS23 template DNA (fragment 2). Holden group |

**Table S2.** Tilt-series collections used for FtsZ_D212A_ subtomogram averaging.

| Dataset | Dataset 1 | Dataset 2 | Dataset 3 | Dataset 4 | Dataset 5 |
|---|---|---|---|---|---|
| Tilt scheme | Dose symmetric | Dose symmetric | Dose symmetric | Dose symmetric | Bidirectional |
| Energy filter slit width (eV) | 10 | 10 | 10 | 10 | 10 |
| Tilt span (°) | -65 to 55 ° | -65 to 55 ° | -48 to 72 ° | -52 to 68 ° | -42 to 66 ° |
| Tilt increment (°) | 3 | 3 | 3 | 3 | 3 |
| Dose per tilt | 3.66 | 3.66 | 3.7 | 3.7 | 4 |
| Total dose (eÅ<sup>-2</sup>) | 150 | 150 | 151.7 | 151.7 | 148 |
| Total tilts per series | 41 | 41 | 41 | 41 | 37 |
| Defocus range (μm) | -2 μm to -4 μm | -2 μm to -4 μm | -1.5 to -4 μm | -1.5 to -3.5 μm | -1 to -3.5 μm |
| Total tilt series | 21 | 25 | 164 | 136 | 195 |

**Table S3.** Tilt-series collections not used in FtsZ_D212A_ STA.

| Sample | Dividing C41 | Dividing C41 | FtsZ<sub>WT</sub> overexpression | FtsZ<sub>Δ2-10</sub> overexpression | FtsZ<sub>Δ2-6</sub> overexpression |
|---|---|---|---|---|---|
| Tilt scheme | Dose symmetric | Dose symmetric | Bidirectional | Dose symmetric | Bidirectional |
| Energy filter slit width (eV) | 10 | 6 | 4 | 10 | 6 |
| Tilt span (°) | -42 to 66 ° | -65 to 55 ° | -66 to 42 ° | -55 to 55 ° | -42 to 66 ° |
| Tilt increment (°) | 2 | 2 | 2 | 2 | 2 |
| Dose per tilt | 3.52 | 3.57 | 3.59 | 3.62 | 3.57 |
| Total dose (eÅ<sup>-2</sup>) | 193.6 | 196.35 | 197.45 | 202.72 | 196.35 |
| Total tilts per series | 55 | 55 | 55 | 56 | 55 |
| Defocus range (µm) | -2 µm to -4 µm | -2 µm to -4 µm | -2 µm to -4 µm | -2 µm to -4 µm | -2 µm to -4 µm |
| Total tilt series | 13 | 15 | 33 | 94 | 36 |

**Table S4.** SPA data collection settings.

| Sample | EcFtsZ<sub>WT</sub> | EcFtsZ<sub>R174D</sub> | EcFtsZ<sub>Δ2-6</sub> | EcFtsZ<sub>Δ2-10</sub> | TmFtsZ | BsFtsZ<sub>WT</sub> | BsFtsZ<sub>Δ1-11</sub> |
|---|---|---|---|---|---|---|---|
| EMDB | EMD-100276 | N/A | EMD-100278 | EMD-100277 | EMD-100783 | EMD-100880 | N/A |
| PDB | 34YW | N/A | 34YZ | 34YY | 50MZ | 50SS | N/A |
| Voltage (kV) | 300 | 300 | 300 | 300 | 300 | 300 | 200 |
| Magnification | 130,000x | 105,000x | 81,000x | 81,000x | 130,000x | 165,000x | 195,000x |
| Pixel size (Å pixel<sup>-1</sup>) | 0.926 | 0.8256 | 0.928 | 0.928 | 0.926 | 1.1945 | 1.4800 |
| Energy filter | Selectris X | Gatan Quantum | Gatan Quantum | Gatan Quantum | Selectris X | Selectris X | N/A |
| Energy filter slit width (eV) | 10 | 20 | 20 | 20 | 10 | 10 | N/A |
| Total dose (e<sup>-</sup> Å<sup>-2</sup>) | 40 | 40 | 40 | 40 | 40-80 | 50 | 50 |
| Defocus range (µm) | -0.9 to -2.4 | -0.8 to -2.8 | -0.9 to -2.4 | -0.9 to -2.4 | -1.2 to -2.8 µm | -0.6 to -2.2 | -0.6 to -2.2 |
| Total micrographs | 6,227 | 5,224 | 4,423 | 2,185 | 18,514 | 10,115 (0 + 30° tilt) | 6,118 |
| Initial particle images | 5,445,946 | 743,386 | 959,827 | 3,720,528 | 23M | 807,520 | 38,552 |
| Final particle images | 430,462 | 146,966 | 327,719 | 45,616 | 175,806 | 93,099 | 24,001 |
| Map resolution (FSC cutoff 0.143) | 2.69 | N/A | 2.73 | 3.39 | 3.7 | 4.1 | 8.0 |
| Symmetry imposed | D1 | N/A | D1 | D1 | D1 | D1 | N/A |
| Helical symmetry | Rise:43.99 Å<br>Twist: -1.1 ° | N/A | Rise: 45.71 Å Twist: 1.09 ° | Rise: 11.35 Å Twist: 89.67 ° | Rise: 47.01 Å<br>Twist: 0.3 ° | Rise: 45.28 Å<br>Twist: -0.27 ° | N/A |
| **Model refinement** |  |  |  |  |  |  |  |
| Starting model | 8IBN | N/A | 8IBN | 8IBN | EcFtsZ<sub>WT</sub> | AF2 | N/A |
| Model-map resolution (FSC cutoff 0.5) | 1.8 | N/A | 3.6 | 4.3 | 4.14 | 4.3 | N/A |
| **Monomer composition** |  |  |  |  |  |  |  |
| Non-hydrogen atoms | 2236 | N/A | 2276 | 2236 | 2362 | 2181 | N/A |
| Protein residues | 306 | N/A | 311 | 306 | 306 | 303 | N/A |
| Ligands | 1 G2P, 1 MG | N/A | 1 G2P, 1 MG | 1 G2P, 1 MG | 1 G2P, 1 MG | 1 GTP | N/A |
| RMSD bond lengths (Å) | 0.005 | N/A | 0.005 | 0.004 | 0.003 | 0.009 | N/A |
| RMSD bond angles (Å) | 1.542 | N/A | 1.072 | 0.851 | 0.8 | 1.538 | N/A |
| **Validation** |  |  |  |  |  |  |  |
| MolProbity score | 1.69 | N/A | 2.16 | 1.76 | 2.09 | 1.69 | N/A |
| Clashscore | 15.43 | N/A | 25.34 | 18.45 | 14.57 | 8.04 | N/A |
| Rotamer outlier (%) | 0.45 | N/A | 0.44 | 0 | 1.73 | 0.67 | N/A |
| Ramachandran favoured (%) | 99.01 | N/A | 96.12 | 98.36 | 96.38 | 96.22 | N/A |
| Ramachandran disallowed (%) | 0 | N/A | 0 | 0 | 0 | 0 | N/A |

**Table S5.**
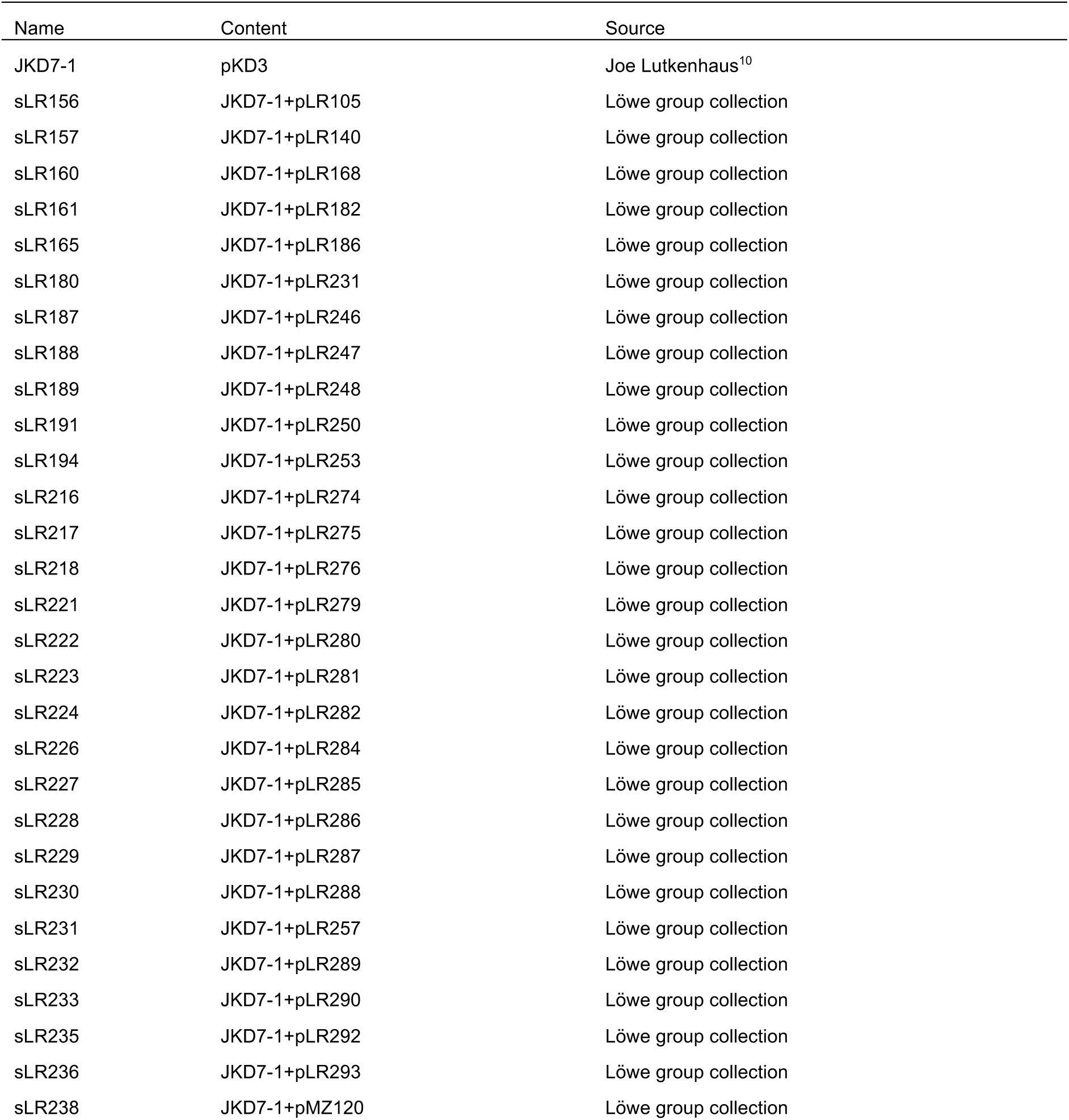
E. coli strains used in this study.

**Table S6.** *B. subtilis* strains used in this study.

| Strain | Genotype | Source/Construction |
|---|---|---|
| sUMS_403 | B. subtilis 168 (WT) | Wehenkel group |
| sUMS_459 | lacA::Pxyl-dcas9, xylR, erm; thrC::Pveg-sgRNA_FtsZ (SpcR) | Wehenkel group |
| sUMS_482 | lacA::Pxyl-dcas9, xylR, ermR; AmyE::Pxyl-FtsZ(gRNA mutated); thrC::Pveg-sgRNA_FtsZ (SpcR) | Wehenkel group |
| sUMS_486 | lacA::Pxyl-dcas9, xylR, ermR; AmyE::Pxyl-FtsZ-ΔN11(gRNA mutated); thrC::Pveg-sgRNA_FtsZ (SpcR) | Wehenkel group |
| sUMS_487 | lacA::Pxyl-dcas9, xylR, ermR; AmyE::Pxyl-FtsZ-K175E(gRNA mutated); thrC::Pveg-sgRNA_FtsZ (SpcR) | Wehenkel group |
| sUMS_488 | lacA::Pxyl-dcas9, xylR, ermR; AmyE::Pxyl-FtsZ-E87K+E90K(gRNA mutated); thrC::Pveg-sgRNA_FtsZ (SpcR) | Wehenkel group |
| MDS23 | Laboratory stock. A prototrophic version of *B. subtilis* 168 | Holden group collection, from Dakes-Stavrakakis *et al*., 2026.<sup>11</sup> |
| SH261 | *amyE::spc Phyperspank-ftsA-ftsZ* | Holden group collection. Transformation of MDS23 with an NEBuilder assembly of an *ftsZ* fragment from the MDS23 genome (amplified with oDMR39-40) and a fragment from pDMR1 (amplified with oDMR41-42). |
| SH262 | *amyE::spc Phyperspank-ftsA-ftsZ<sup>K175D</sup>* | Holden group collection. Transformation of MDS23 with an NEBuilder assembly of linear PCRs generated using oBB64, oDMR68 and SH261 template DNA (fragment 1); oBB65, oDMR69 and SH261 template DNA (fragment 2). |
| SH263 | *amyE::spc Phyperspank-ftsA-ftsZ<sup>E87K, E90K</sup>* | Holden group collection. Transformation of MDS23 with an NEBuilder assembly of linear PCRs generated using oBB64, oDMR84 and SH261 template DNA (fragment 1); oBB65, oDMR85 and SH261 template DNA (fragment 2). |
| SH264 | *ftsZ:ftsZ-gfp-cat* | Holden group collection. Transformation of MDS23 with a PCR fragment generated using oBB66, oBB67 and SH007 template DNA. |
| SH265 | *amyE::spc Phyperspank-ftsA-ftsZ, ftsZ:ftsZ-gfp-cat* | Holden group collection. Transformation of SH261 with a PCR fragment generated using oBB66, oBB67 and SH264 template DNA |
| SH266 | *amyE::spc Phyperspank-ftsA-ftsZ<sup>K175D</sup>, ftsZ:ftsZ-gfp-cat* | Holden group collection. Transformation of SH262 with a PCR fragment generated using oBB66, oBB67 and SH264 template DNA |
| SH267 | *amyE::spc Phyperspank-ftsA-ftsZ<sup>E87K, E90K</sup>, ftsZ:ftsZ-gfp-cat* | Holden group collection. Transformation of SH264 with an NEBuilder assembly of linear PCRs generated using oBB64, oDMR84 and SH261 template DNA (fragment 1); oBB65, oDMR85 and SH261 template DNA (fragment 2). |

**Table S7.**
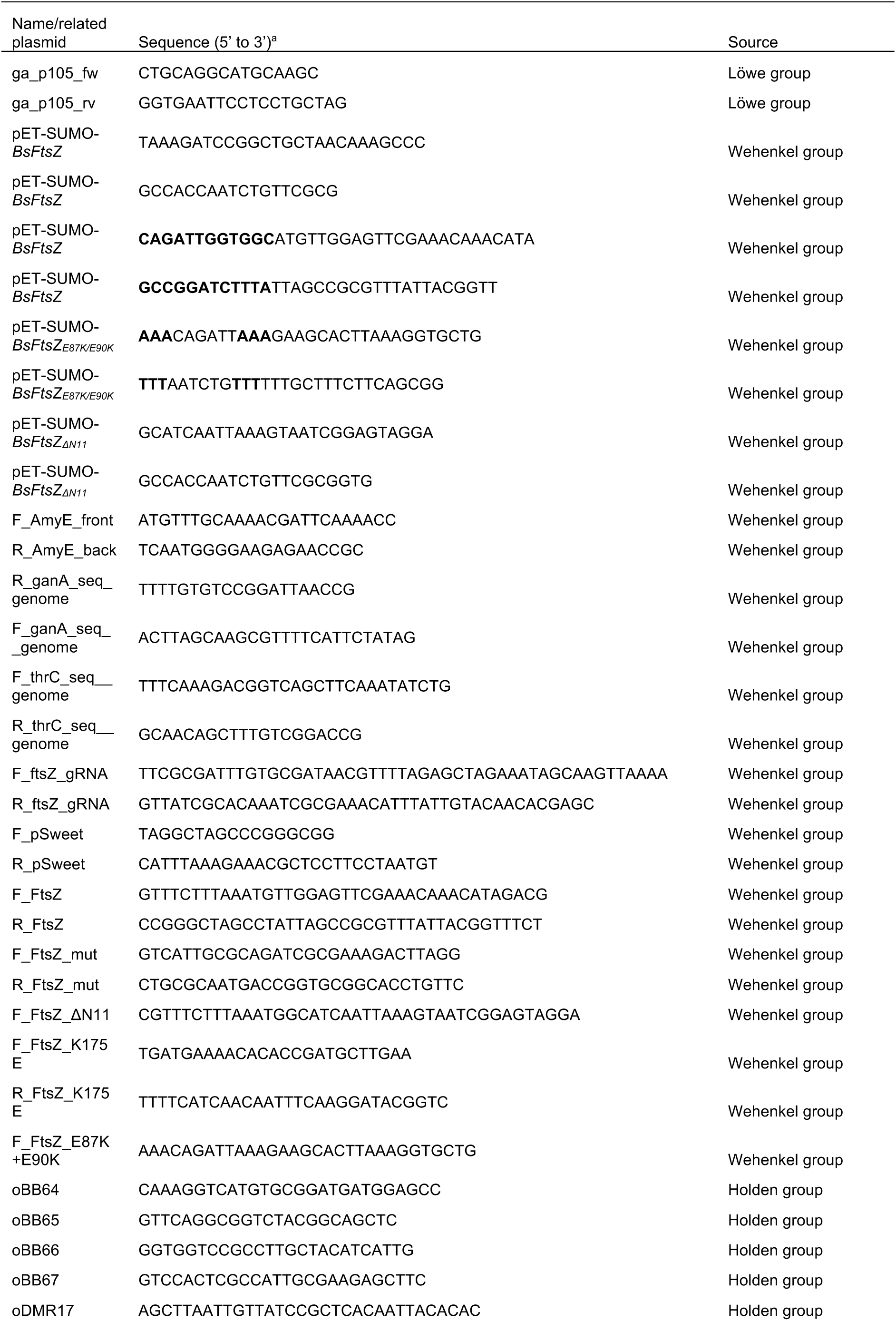
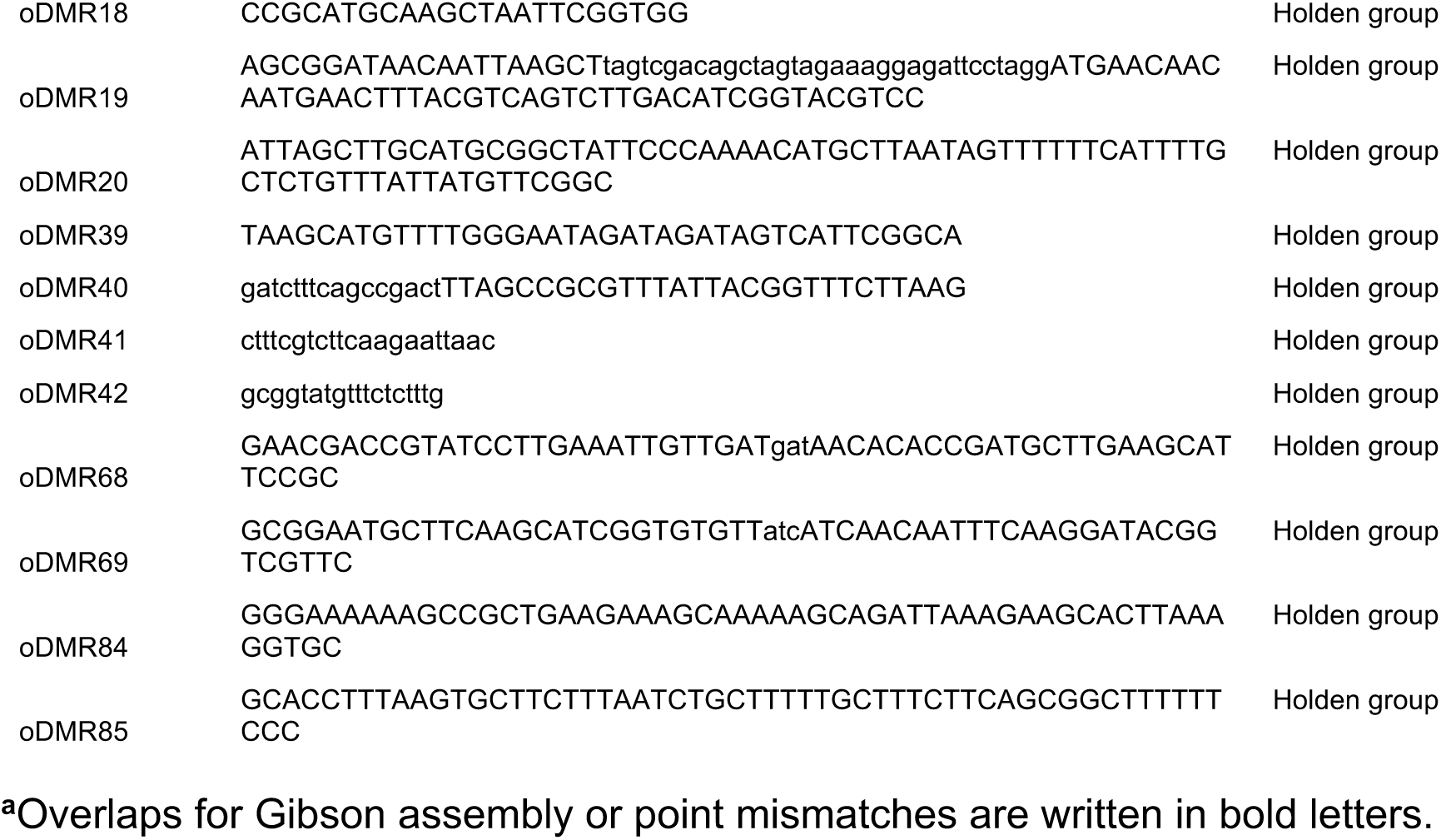
Oligonucleotides.

**Table S8.** Live cell imaging P values from Mann Whitney U test.

| **Dataset comparison** | **p value** |
|---|---|
| Wild type vs. Wild type, O/E | 0.011 |
| Wild type, O/E vs. K175D, O/E | 4.45 x 10<sup>-7</sup> |
| Wild type O/E vs. E87K, E90K, O/E | 6.5 x 10<sup>-19</sup> |

**Table S9.**
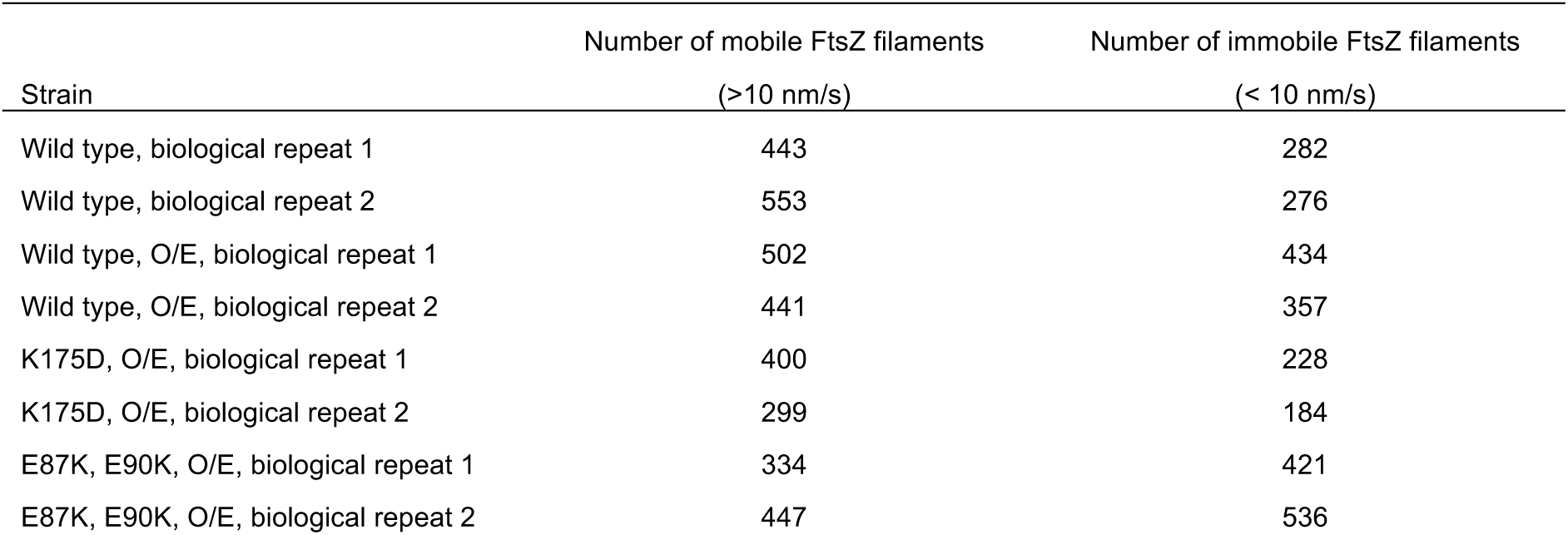
Number of FtZ-GFP filaments analysed in live-cell imaging.

| Strain | Number of mobile FtsZ filaments<br>(>10 nm/s) | Number of immobile FtsZ filaments<br>(< 10 nm/s) |
|---|---|---|
| Wild type, biological repeat 1 | 443 | 282 |
| Wild type, biological repeat 2 | 553 | 276 |
| Wild type, O/E, biological repeat 1 | 502 | 434 |
| Wild type, O/E, biological repeat 2 | 441 | 357 |
| K175D, O/E, biological repeat 1 | 400 | 228 |
| K175D, O/E, biological repeat 2 | 299 | 184 |
| E87K, E90K, O/E, biological repeat 1 | 334 | 421 |
| E87K, E90K, O/E, biological repeat 2 | 447 | 536 |

## Supplementary Movies

**Movie S1.** Tomograms of dividing *E. coli* C41(DE3) cells shown in Figure 1AB. The movie plays through Z-planes before revealing 3D segmented membrane, ribosomes and FtsZ. These segmentations are shown as 2D slices in Figure 1AB.

**Movie S2.** Tomogram of an *E. coli* C41(DE3) cell overexpressing FtsZ_D212A_ shown in Figure 1C. The movie plays through Z-planes before revealing 3D segmented membrane, ribosomes and FtsZ. These segmentations are shown as 2D slices in Figure 1C.

**Movie S3.** Orientations of refined particle coordinates shown in Figure 1H. Refined coordinates are show with respect to the 6.9 Å STA map. The red arrow represents the direction of the unresolved CTT. Particles are shown alongside the membrane segmentation (also shown in Movie S2) to visualise FtsZ orientation within the cell.

**Movie S4.** Tomogram of an *E. coli* C41(DE3) cell overexpressing FtsZ (shown in Figure 2C) followed by a tomogram of an *E. coli* C41(DE3) overexpressing FtsZ_R174A_ (shown in Figure 2D). The movie plays through Z-planes before revealing 3D segmented membrane, ribosomes and FtsZ. These segmentations are shown as 2D slices in Figure 2CD.

**Movie S5.** Tomograms of *E. coli* C41(DE3) cells overexpressing FtsZ_Δ2-6_ and FtsZ_Δ2-10_. The first tomogram corresponds to Figure 4D and shows 3D segmentations of membrane, ribosomes and FtsZ. Two other tomograms that were not segmented or shown in the main text follow and are of *E. coli* cells overexpressing FtsZ_Δ2-10_.

**Movie S6.** Coarse-grained MD simulations of treadmilling FtsZ with lateral interactions modelled. Corresponds to Figure 5 and shows an example trajectory for each condition tested.

